# Engineered orthogonal translation systems from metagenomic libraries expand the genetic code

**DOI:** 10.1101/2025.10.30.685624

**Authors:** Kosuke Seki, Michael T. A. Nguyen, Petar I. Penev, Jillian F. Banfield, Farren J. Isaacs, Michael C. Jewett

**Author notes:** Department of Bioengineering and Therapeutic Sciences, University of California, San Francisco; San Francisco, CA 94143, USA.

## Abstract

Genetic code expansion with non-canonical amino acids (ncAAs) opens new opportunities for the function and design of proteins by broadening their chemical repertoire. Unfortunately, ncAA incorporation is limited both by a small collection of orthogonal aminoacyl-tRNA synthetases (aaRSs) and tRNAs and by low-throughput methods to discover them. Here, we report the discovery, characterization, and engineering of a UGA suppressing orthogonal translation system mined from metagenomic data. We developed an integrated computational and experimental pipeline to profile the orthogonality of >200 tRNAs, test >1,250 combinations of aaRS:tRNA pairs, and identify the AP1 TrpRS:tRNA^Trp^_UCA_ as an orthogonal pair that natively encodes tryptophan at the UGA codon. We demonstrate that the AP1 TrpRS:tRNA^Trp^_UCA_ is highly active in cell-free and cellular contexts. We then use *Ochre*, a genomically recoded *Escherichia coli* strain that lacks UAG and UGA codons, to engineer an AP1 TrpRS variant capable of 5-hydroxytryptophan incorporation at an open UGA codon. We anticipate that our strategy of integrating metagenomic bioprospecting with cell-free screening and cell-based engineering will accelerate the discovery and optimization of orthogonal translation systems for genetic code expansion.

## Introduction

The site-specific incorporation of non-canonical amino acids (ncAAs) expands protein properties, structures, and functions.^1–3^ This functional expansion enables the study of post-translational modifications and the development of next-generation therapeutics and biomaterials.^4–6^ Over 200 different ncAAs have been incorporated into proteins using an orthogonal aminoacyl-tRNA synthetase (aaRS) and tRNA pair called an orthogonal translation system (OTS).^7^ Yet, identifying an OTS suitable for a ncAA and codon of interest is a key bottleneck in genetic code expansion. Traditional OTSs such as the *Methanocaldococcus jannaschii* TyrRS:tRNA^Tyr^_CUA_ and the *Methanosarcina barkeri, Methanosarcina mazei,* and *Methanomethylophilus alvus* PylRSs:tRNAs^Pyl^_CUA_ are limited by aaRS substrate promiscuity and often must be laboriously evolved to accept ncAAs with diverse chemistries.^8–10^

A promising approach to expand the chemistries and codons available for genetic code expansion is to discover new OTSs.^11–14^ The primary challenge is to find an aaRS:tRNA pair that is orthogonal to endogenous aaRSs, tRNAs, and amino acids in the host of interest. In some cases, aaRS:tRNA pairs from heterologous organisms are orthogonal due to the evolution of divergent tRNA recognition mechanisms as in PylRS:tRNA^Pyl^_CUA_.^15,16^ In general, however, OTS discovery campaigns are challenging because aaRS:tRNA pairs that sufficiently satisfy orthogonality are rare and because the combinatorial space of candidate aaRS:tRNA pairs is vast. Therefore, developing methods that quickly identify functional OTSs and finding sources of aaRSs and tRNAs that are enriched for orthogonality would be broadly useful for genetic code expansion.

To overcome these challenges, we develop an integrated workflow that combines metagenomic prospecting, high-throughput cell-free approaches to screen and characterize orthogonal aaRSs and tRNAs, and aaRS engineering in recoded *Escherichia coli* strains (**Fig. 1**). We specifically characterize metagenome-derived suppressor tRNAs and aaRSs from bacteriophages which, despite having garnered recent attention for widespread stop codon reassignment, are untapped and uncharacterized sources of translational machinery (**Fig. 1**).^17,18^ We curated a set of > 200 suppressor tRNAs for all three stop codons and characterized their activity and orthogonality in *E. coli* based cell-free reactions. We then evaluated a panel of aaRSs, tested > 1,250 combinations of aaRS:tRNA pairs, and identified two candidate OTSs. We characterized the AP1 TrpRS:tRNA_UCA_, which orthogonally suppresses the UGA codon as Trp in *E. coli*. Finally, we engineered the AP1 TrpRS to incorporate 5-hydroxytryptophan (5HTP) at UGA using *Ochre*, a genomically engineered *E. coli* strain optimized for UAG and UGA suppression.^19^ In total, our work shows how integrated computational and experimental workflows can be used to discover new OTSs.

**Figure 1:**
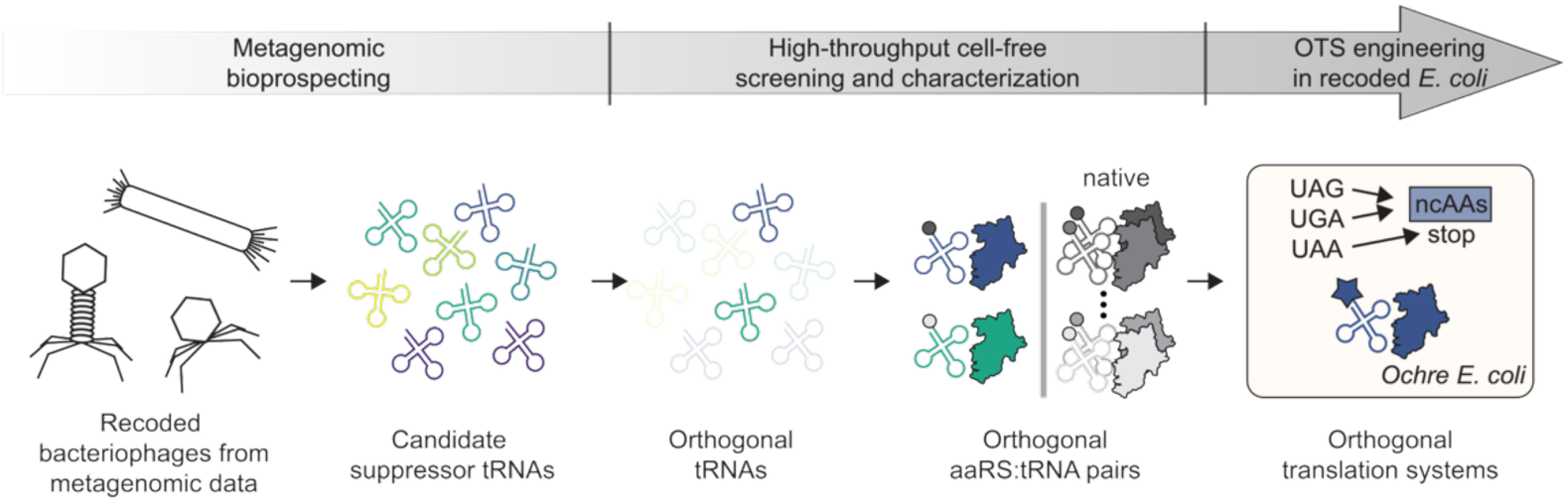
Integrated computational and experimental workflows enable the identification of new OTSs. Metagenomic prospecting, high-throughput screening of tRNAs and aaRSs in cell-free expression systems, and OTS engineering in recoded *E. coli* strains enables the identification and discovery of new OTSs.

## Results

### Identification and phylogeny of suppressor tRNAs in phage genomes

Our goal was to develop a robust, high-throughput, and generalizable workflow to identify aaRS:tRNA candidates from genomic libraries, starting with putative suppressor tRNAs. We hypothesized that bacteriophages would be privileged sources of suppressor tRNAs due to recent literature showing widespread recoding of stop codons in their genomes.^17,18^ We curated a library of 213 suppressor tRNAs from bacteriophage genomes that were assembled from metagenomic datasets (**Fig. 2**, **Table S1**).^18^ We refer to each tRNA using standard tRNA nomenclature along with a unique integer identifier (i.e. tRNA_CUA_-1 for the first tRNA suppressor for UAG) (**Table S1**). Most of the library consists of tRNAs_CUA_ (76%, 162/213), while 9% are tRNAs_UUA_ (20/213), and 15% are tRNAs_UCA_ (31/213). This is consistent with previous observations that UAG recoding is widespread in bacteriophages.^18^ The library contains tRNA sequences that are predicted with high and low confidence, and we retained low confidence sequences (11/20 tRNA_UUA_ and 9/31 tRNAs_UCA_) as they could be easily accommodated in our high-throughput screen (described below). Phylogenetic analysis of these tRNA sequences reveals that tRNA sequences segregate into distinct clades according to their predicted anticodon (**Fig. 2**). This may suggest that suppressor tRNA sequences evolve under the constraint of accurately decoding their respective codons.

**Figure 2:**
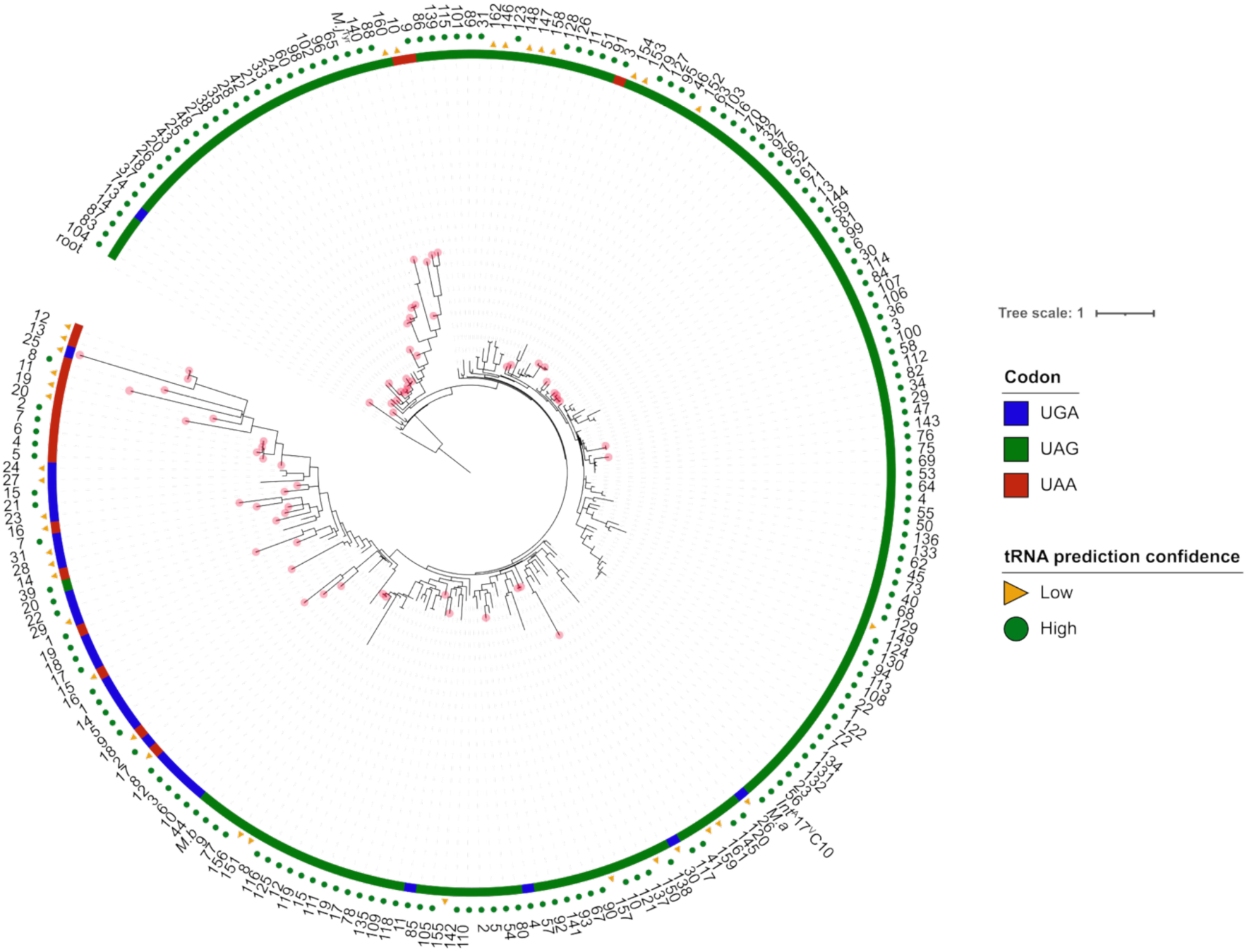
A library of suppressor tRNA sequences from bacteriophages in metagenomic data provides candidate orthogonal tRNA sequences. A phylogenetic tree of suppressor tRNA sequences. Colors denote predicted codon for each tRNA, and shapes indicate confidence scores. High confidence scores are those > 30 as scored by tRNAscan-SE^20^. Specific tRNA sequences highlighted with a red circle on the branches are those identified to be orthogonal based on Figure 2. *M.j* = *M. jannaschii*, *M.a* = *M. alvus* and *M.b* = *M. barkeri*.

### Development of a high-throughput tRNA expression platform

With putative suppressor tRNAs in hand, we next developed a screening platform to transcribe and evaluate suppressor tRNAs in high-throughput (**Fig. 3a**). Current methods for tRNA expression are neither generalizable nor scalable; *in vivo* expression can cause cellular toxicity and requires laborious plasmid cloning workflows,^11^ and *in vitro* tRNA transcription requires either a +1 G on the tRNA or a 5’ hammerhead ribozyme that must be cleaved and removed from mature tRNA.^21,22^ We circumvented these limitations by leveraging tRNA processing activity in cell-free gene expression (CFE) reactions, which activate transcription and translation in the crude extracts of cells, to robustly transcribe and mature functional tRNAs.^23^ Candidate tRNAs were first designed as fusions to a 5’ RNase P tag.^24^ The RNase P tag is a strong substrate for T7 RNA polymerase and standardizes transcription of tRNAs which would otherwise be poor substrates. RNase P-tagged tRNAs were then synthesized by *in vitro* transcription (IVT). Crude tRNAs were added directly to CFE reactions, where tRNAs are simultaneously matured by RNase P in cell extracts through cleavage of the RNaseP tag and evaluated for function by measuring suppression of a premature stop codon in a protein of interest. We used superfolder Green Fluorescent Protein (sfGFP) with a stop codon at T216 (216X-sfGFP, where X is UAG, UAA, or UGA) as a reporter for suppression activity. This method has several advantages; it is (i) entirely cell-free, requiring no cloning or transformation steps, (ii) rapid (PCR: 1.5 hrs, *in vitro* transcription: 4 hrs, CFE: ∼4 hrs), and (iii) high-throughput, enabling the functional characterization of hundreds of suppressor tRNAs in a single 384-well plate. A similar method implemented in a PURE cell-free translation system to transcribe tRNAs was reported during the preparation of our work;^25^ our workflow avoids the need to purify RNase P by leveraging endogenous RNase P activity in cell extracts and thus simplifies the assay.

**Figure 3.**
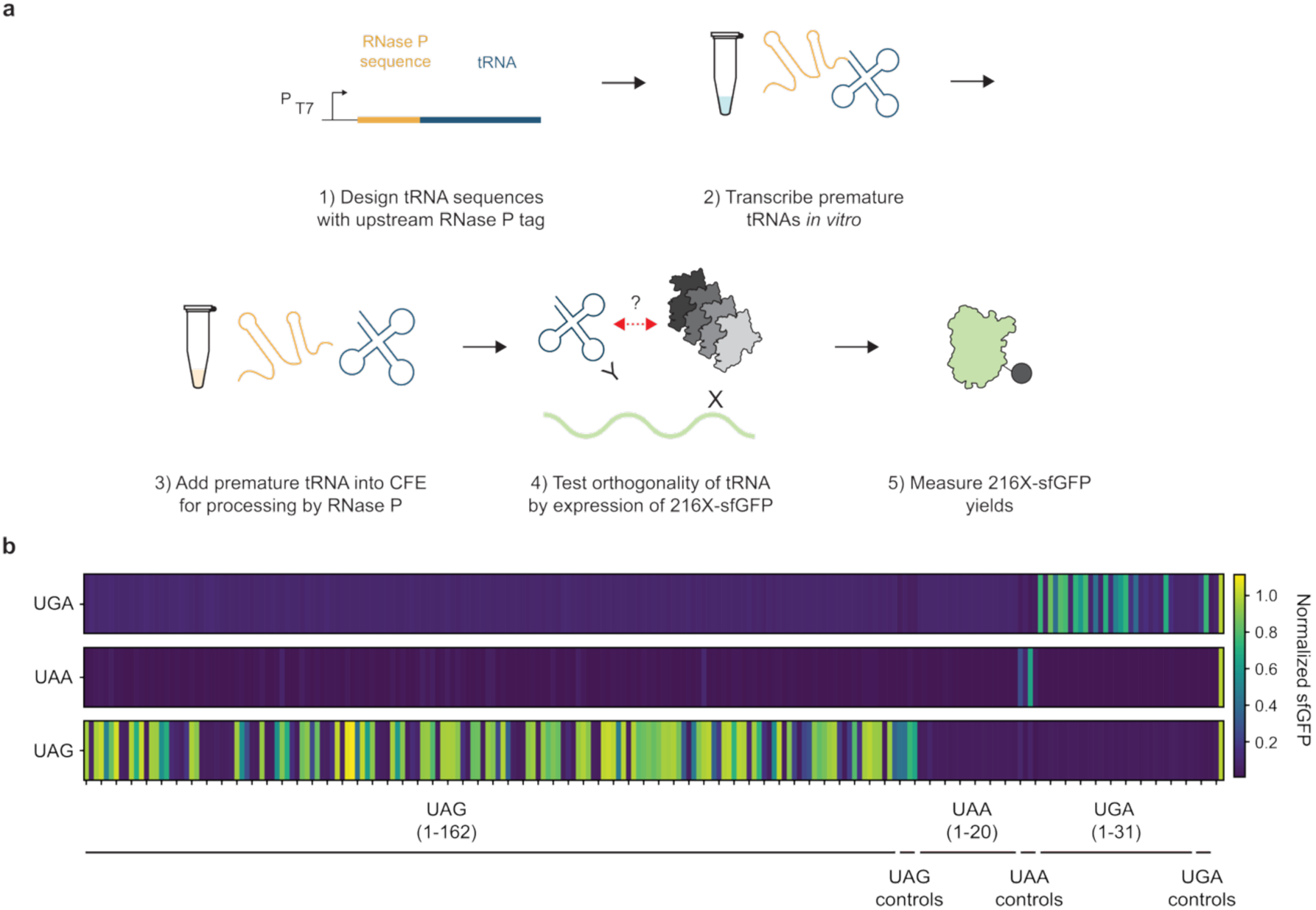
A high-throughput cell-free workflow for tRNA expression and characterization enables rapid profiling of tRNA orthogonality. (**a**) Schematic of high-throughput tRNA evaluation method. tRNAs are purchased as short linear DNA sequences, transcribed *in vitro* with T7 RNA polymerase, matured into functional tRNAs by endogenous RNase P in cell extracts, and evaluated for stop codon suppression activity. (**b**) A high-throughput screen of bioinformatically identified suppressor tRNAs revealed 73 orthogonal tRNA candidates. Suppressor tRNAs are grouped by codon along with internal controls of the *M. jannaschii* tRNA^Tyr,^ the *M. barkeri* tRNA^Pyl^, the *M. alvus* tRNA^Pyl^, and the ^A^17 ^V^C10 *Int* tRNA^Pyl^. Heatmap shows representative data (see **Supplementary Files**) from three independent experiments with similar results. All signal is normalized to WT sfGFP (far right).

Our CFE platform enables robust expression and characterization of orthogonal tRNAs. Proof-of-concept experiments with the *M. alvus* tRNA^Pyl^_CUA_, a model orthogonal tRNA, showed that the 5’ RNase P tag was efficiently cleaved in *E. coli* cell extracts and that the resulting tRNA was functional for UAG suppression in CFE in the presence of its cognate aaRS and azidolysine, a compatible ncAA (**Fig. S1a,b,** see *M. alvus* column). We evaluated the generalizability of this platform by testing phage-derived tRNAs_CUA_ which are known substrates for *E. coli* GlnRS.^26,27^ All six tRNAs_CUA_ suppressed UAG efficiently when expressed with the RNase P tag (**Fig. S1b**). In contrast, tRNAs expressed using plasmids or RNase P tag-less templates showed inconsistent 216UAG-sfGFP synthesis, which we speculate is due to either poor transcription or poor tRNA maturation.

### High-throughput screening of suppressor tRNAs from phage genomes

Using our high-throughput tRNA expression and characterization platform, we next screened our library of 213 suppressor tRNAs for orthogonal tRNAs. We defined an orthogonal tRNA as one that is unable to support 216X-sfGFP expression, as we expected that a non-orthogonal tRNA would be aminoacylated by endogenous aaRSs and would support expression of full-length sfGFP. We validated this criterion by showing that (i) the *M.barkeri* tRNA_CUA_^Pyl^, a known orthogonal tRNA, does not support 216X-sfGFP expression in the absence of its cognate aaRS (**Fig. S2a**) and that (ii) T216X substitutions, which could occur by suppression from non-orthogonal aminoacylated tRNAs, do not disrupt sfGFP fluorescence (**Fig. S2b**). Loss of fluorescence in the presence of a candidate suppressor tRNA should therefore indicate orthogonality.

We screened the activity and orthogonality of all suppressor tRNAs against all three stop codons, resulting in 639 combinations of tRNAs and codons (**Fig. 3b**). We included a set of commonly used OTSs (the *M. jannaschii* tRNA^Tyr^:BpyRS,^28^ the *M. barkeri* tRNA^Pyl^:chPylRS (IPYE),^9^ the *M. alvus* tRNA^Pyl^:PylRS, and the ^A^17 ^V^C10 *Int* tRNA^Pyl^:Lum1 PylRS^29^) as controls. Within the UAG and UGA codons, we observed a wide range of activity for suppressor tRNAs (**Fig. 3b**). We found that 90/162 UAG suppressors and 9/31 UGA suppressors achieved protein yields of at least 50% of WT sfGFP. We binned these as active, non-orthogonal tRNAs. Notably, we identified 42 UAG suppressors, 20 UAA suppressors, and 11 UGA suppressors with near-background levels of 216X-sfGFP. These 73 tRNAs were categorized as orthogonal tRNA candidates. As it is possible that some of these tRNAs were not correctly transcribed or processed, we randomly selected a subset of these tRNAs and confirmed that their DNA templates were amplified by PCR, transcribed in IVT, and processed by RNase P to mature tRNAs (**Fig. S3a-c**). We therefore moved these tRNAs forward as orthogonal candidates.

Apart from identifying orthogonal tRNA candidates, our dataset highlights failure modes for orthogonality in suppressor tRNAs. Using representative non-orthogonal tRNAs, we analyzed suppression events in 216UAG-sfGFP and 216UGA-sfGFP using intact protein electrospray ionization mass spectrometry (ESI-MS). Of the tested tRNAs_CUA_, 28/29 incorporated glutamine at UAG and the remaining tRNA_CUA_ (tRNA_CUA_-151) incorporated alanine (**Fig. S4**). Although glutamine and lysine are nearly isobaric, we hypothesize that UAG is reassigned to glutamine because this reassignment is commonly observed in nature^30,31^ and because *E. coli* GlnRS recognizes tRNA^Gln^_CUA_ found in some *E. coli* strains.^32^ Reassignment to alanine, however, has been observed infrequently.^33,34^ Analysis of 13 tRNAs_UCA_ showed incorporation of tryptophan in all cases (**Fig. S5**), a frequently observed stop codon reassignment mechanism in nature.^31^ The scale of our dataset highlights general patterns for non-specific aminoacylation of suppressor tRNAs_CUA_ and tRNAs_UCA_. In the future, avoiding tRNA identity elements for GlnRS and TrpRS may provide a favorable route to identifying or engineering orthogonal tRNAs.

### Identification and characterization of cognate aaRSs for orthogonal tRNAs

Next, we bioinformatically screened for cognate aaRSs to our orthogonal tRNAs to discover orthogonal aaRS:tRNA pairs. First, we hypothesized that bacterial hosts may harbor cognate aaRSs for orthogonal tRNAs found in bacteriophage genomes. We searched for bacterial hosts of the phage whose high-quality genome assembly contained the orthogonal tRNA_CUA_-103. A potential host from the *Prevotella* clade was identified through CRISPR spacer analysis. We predicted 15 putative aaRSs from this genome: MetRS, LeuRS, GluRS, AspRS, CysRS, TyrRS, GlnRS, LysRS, PheRS, AsnRS, GlyRS, and SerRS (**Table S2**). The SerRS contained an in-frame UGA codon which we recoded as UGG based on the common UGA > W reassignment. Second, in a broader approach, we identified aaRSs that co-localized with orthogonal tRNAs in putative “code change” operons responsible for stop codon suppression in bacteriophage genomes. We found two such TrpRS:tRNA candidate pairs: the AP1 TrpRS:tRNA^Trp^_UCA_ (tRNA_UCA_-11, hereafter referred to as AP1 tRNA^Trp^_UCA_) and the HF2 TrpRS:tRNA^Trp^_UCA_ (tRNA_UCA_-13, hereafter referred to as HF2 tRNA^Trp^_UCA_).^18^ We hypothesized that these aaRS:tRNA pairs mediate reassignment of stop codons and are functional and orthogonal in *E. coli* extracts.

Enabled by the high-throughput nature of cell-free reactions, we screened >1,250 combinations of aaRS:tRNA pairs for 216X-sfGFP expression using co-expression assays. Within the 15 aaRSs identified in the broad sequence search, we found that the putative GlnRS and TyrRS enzymes were functional only with tRNAs_CUA_ (**Fig. 4a, S6-S8**). The best performing tRNAs for the GlnRS and TyrRS were tRNA_CUA_-59 and tRNA_CUA_-27, respectively (**Fig. 4a**, red boxes). Yields of 216UAG-sfGFP using the GlnRS: tRNA_CUA_-59 pair were comparable to the WT sfGFP control and showed 4-fold greater activity compared to the TyrRS: tRNA_CUA_-27 pair. We next screened the AP1 and HF2 TrpRSs against orthogonal tRNAs_UCA_ (**Fig. 4b**). As expected, they were most active in the presence of their cognate tRNA (**Fig. 4b**, red boxes). Their tRNA specificities are unique but not mutually orthogonal; the AP1 TrpRS recognized tRNA_UCA_-29 while the HF2 TrpRS did not, and non-cognate pairs (AP1 TrpRS:HF2 tRNA_UCA_ and vice versa) were functional. Differences in tRNA identity elements, such as the G73 discriminator base in AP1 tRNA_UCA_ and tRNA_UCA_-29 compared to the A73 discriminator base in HF2 tRNA_UCA_, may contribute to the observed specificities (**Fig. S9**).^35^ In total, these results identified functional aaRS:tRNA pairs for the UAG and UGA codons.

**Figure 4:**
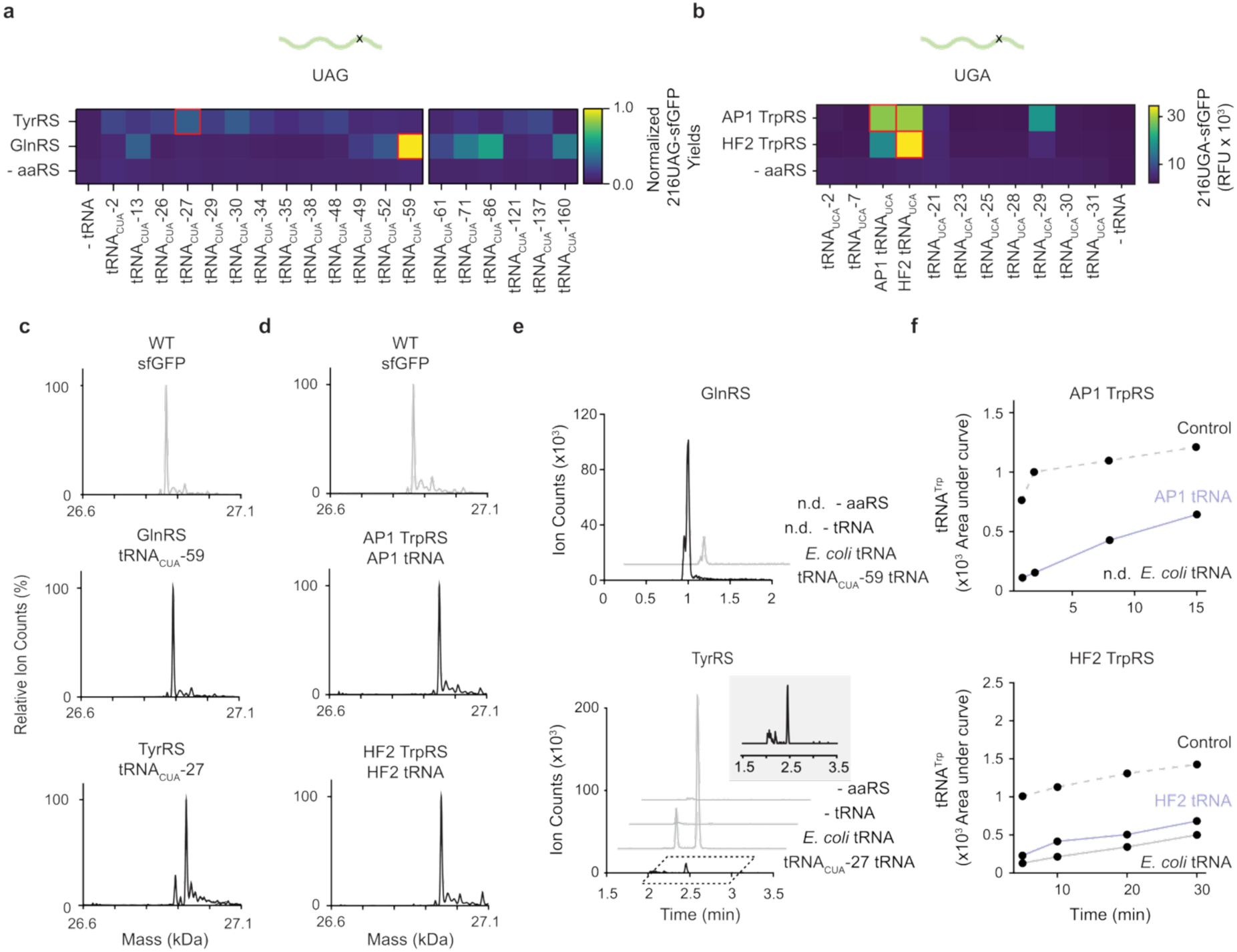
Identification of orthogonal aaRS:tRNA pairs from high-throughput screens of thousands of aaRS:tRNA combinations. (**a**) Broad screening of a panel of 16 aaRSs against all orthogonal tRNAs identifies several active aaRS:tRNA_CUA_ pairs. CFE was performed by coexpressing the aaRS of interest and the 216X-sfGFP reporter supplemented with an *in vitro* transcribed tRNA of interest. Data shown are curated from the full dataset presented in **Fig. S6**. Data are representative of two independent experiments and is normalized to a WT sfGFP control. (**b**) Focused screening of TrpRSs identifies functional TrpRS:tRNA_UCA_ pairs and highlights distinct tRNA recognition mechanisms. Experiments used purified AP1 and HF2 TrpRSs. Red boxes in (a-b) highlight aaRS:tRNA pairs selected for further characterization (**c-d**) ESI-MS of intact proteins confirms assignment of putative aaRSs. Compared to the WT (26864.10 Da), the mass shifts of the products from GlnRS: tRNA_CUA_-59 (26891.05 Da), TyrRS: tRNA_CUA_-27 (26925.65 Da), AP1 (26949.60 Da), and HF2 (26949.50 Da) are consistent with incorporation of Gln, Tyr, Trp, and Trp, respectively. (**e-f**). *in vitro* aminoacylation reactions highlight orthogonality of aaRSs. GlnRS: tRNA_CUA_-59 and the AP1 TrpRS:tRNA^Trp^_UCA_ pairs appear functionally orthogonal in *E. coli*, while the TyrRS: tRNA_CUA_-27 and HF2 TrpRS:tRNA^Trp^_UCA_ pairs are not orthogonal in *E. coli*. n.d. = not detected. Control in **f** is *E. coli* TrpRS with total *E. coli* tRNA. Data in (**c-f**) were collected twice; a representative dataset is shown.

We confirmed the functional assignment of these aaRSs. ESI-MS of purified 216UAG-sfGFP from reactions containing the GlnRS: tRNA_CUA_-59 pair and the TyrRS: tRNA_CUA_-27 pair displayed mass shifts consistent with incorporation of Gln and Tyr, respectively (**Fig. 4c**). We observed products corresponding to Gln readthrough from the TyrRS: tRNA_CUA_-27 pair, which is likely a result of competition with endogenous *E. coli* tRNA^Gln^.^27,36^ Analysis of purified 216UGA-sfGFP from reactions containing the AP1 TrpRS:tRNA^Trp^_UCA_ and the HF2 TrpRS:tRNA^Trp^_UCA_ also confirmed that the TrpRSs enable Trp incorporation at UGA (**Fig. 4d**).

To evaluate the orthogonality of the aaRS:tRNA pairs, we measured aaRS activity against either their cognate tRNA or total *E. coli* tRNA at estimated physiological concentrations using an *in vitro* aminoacylation assay. This assay uses LC-MS to detect the aminoacylated adenosine (aa-A, where aa is the three-letter code for an amino acid) produced after RNase-A digestion of aminoacylation reactions.^37^ The Gln-A and Tyr-A products were first identified using positive control reactions with *E. coli* aaRSs and total *E. coli* tRNA (**Fig. S10**). We re-tested the orthogonality of the tRNA_CUA_-27 and tRNA_CUA_-59 tRNAs *in vitro* with *E. coli* aaRSs and found that, although tRNA_CUA_-27 remained orthogonal to the *E. coli* TyrRS, tRNA_CUA_-59 was aminoacylated by *E. coli* GlnRS (**Fig. S10**). However, tRNA_CUA_-59 was found to be orthogonal in CFE which better mimics the competitive translation environment (**Fig. 4a**). We then tested the aaRSs for orthogonality. We found that the GlnRS was highly active with tRNA_CUA_-59 and showed weak activity with total *E. coli* tRNA (**Fig. 4e**). On the other hand, the putative TyrRS produced a trace amount of Tyr-A product in the presence of tRNA_CUA_-27 (**Fig. 4e**, see inset) but had significant cross-reactivity with total *E. coli* tRNA. We hypothesize that the GlnRS: tRNA_CUA_-59 pair, but not the TyrRS: tRNA_CUA_-27 pair, may function orthogonally for the UAG codon. Weak non-specific aminoacylation of total *E. coli* tRNA has been observed even with established OTSs, suggesting that a small amount of non-specificity may not break orthogonality due to competition with endogenous aaRSs during translation.^11,38^

We then assessed the orthogonality of the AP1 TrpRS:tRNA_UCA_ pair and the HF2 TrpRS:tRNA_UCA_ pair. As before, the Trp-A product was identified by aminoacylating total *E. coli* tRNA with *E. coli* TrpRS (**Fig. S11**). Consistent with the ESI-MS analysis, aminoacylation assays showed production of a Trp-A product in the presence of the AP1 TrpRS:tRNA_UCA_ pair and the HF2 TrpRS:tRNA_UCA_ pair. Encouraged by this result, we analyzed Trp-A formation over time (**Fig. 4f**). The HF2 TrpRS synthesized similar amounts of Trp-A using either total *E. coli* tRNA or its cognate tRNA_UCA_. We found that the HF2 TrpRS nonspecifically aminoacylated *E. coli* tRNA^Trp^_CCA_ (**Fig. S12**). In contrast, the AP1 TrpRS showed robust activity against its cognate tRNA_UCA_ and did not synthesize detectable Trp-A product in the presence of total *E. coli* tRNA (**Fig. 4f**). This suggests that the AP1 TrpRS:tRNA_UCA_ pair is a natural UGA suppressing translation system that is orthogonal to *E. coli* translation machinery. These cell-free workflows show how aaRS:tRNA pairs can be rapidly identified and characterized for orthogonality.

### In vivo assessment of the AP1 TrpRS:tRNA_UCA_

We next assessed the function of the AP1 TrpRS:tRNA_UCA_ pair in living *E. coli* cells. We tested three common *E. coli* laboratory strains (MG1655, BL21, and DH10β) as well as *Ochre*, an *E. coli* strain where all genomic UAG and UGA stop codons are synonymously recoded to UAA.^19^ These strains also differ in RF2 termination strength; mutations in RF2 weaken termination at UGA in MG1655 (T246A), DH10β (T246A), and *Ochre* (E170K and S205P), while BL21 has a wild-type RF2.^19,39^ First, we assessed UGA suppression activity *in vivo* by measuring expression of GFP containing an N-terminal 3x UGA elastin-like-polypeptide (ELP-3xUGA-GFP) in the *Ochre* strain. Expression was observed only when the tRNA and TrpRS were coexpressed, showing that the AP1 translation system has robust UGA suppression activity *in vivo* and that the AP1 tRNA_UCA_ remained orthogonal (**Fig. 5a**). We tested the performance of the AP1 translation system in all four *E. coli* strains and found that expression was dependent on the expected RF2 strength, suggesting competition against RF2 as a key variable (**Fig. 5b**). We then assessed its toxicity by measuring *E. coli* growth. MG1655 and DH10β displayed a reduced final OD_600_ as a function of induction level, although the doubling time of all strains during exponential phase was unaffected (**Fig. 5c**, **S13**). These data point towards an RF2-mediated tradeoff where RF2 prevents suppression of UGA by the AP1 tRNA_UCA_ but relieves toxicity resulting from readthrough of endogenous UGA codons. *Ochre* is therefore an ideal strain for UGA suppression due to its attenuated RF2 and lack of endogenous UGA codons. In total, these results showed that the AP1 TrpRS:tRNA_UCA_ pair is functional *in vivo* and identified key determinants for effective UGA suppression.

**Figure 5:**
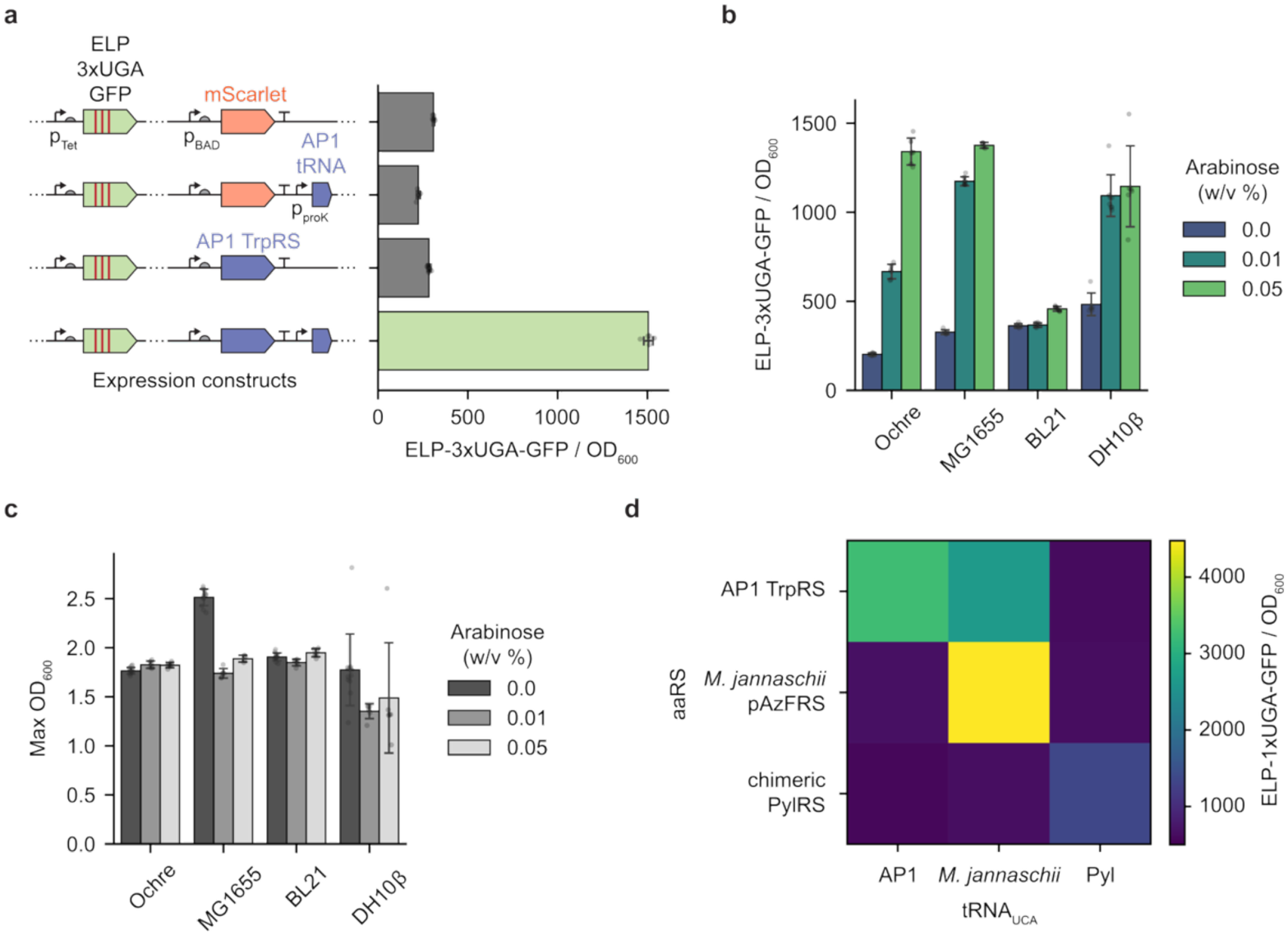
*In vivo* characterization of the AP1 TrpRS:AP1 tRNA_UCA_ pair. (**a**) Bar chart of ELP-3xUGA-GFP normalized by OD_600_ shows that coexpression of the AP1 tRNA_UCA_ and TrpRS is required for UGA suppression. Expression cassettes for AP1 TrpRS:tRNA_UCA_ pair and ELP-3xUGA-GFP reporter construct are labelled on the y axis. The AP1 tRNA is constitutively expressed under the *proK* promoter, the ELP-3xUGA-GFP is controlled by TetR promoter and can be induced with anhydrotetracycline (aTc), and the AP1 TrpRS is controlled by pBAD and can be induced with arabinose. mCherry is used as a negative control. Bars represent the average of n = 6 replicates for all conditions except the + AP1 TrpRS / - tRNA condition, where n = 4 replicates. Error bars represent one standard deviation. (**b**) Bar chart of ELP-3xUGA-GFP signal normalized by OD_600_ in *E. coli* strains Ochre, MG1655, BL21, and DH10β. UGA is efficiently suppressed in DH10β, MG1655, and Ochre, while BL21 shows decreased GFP expression. (**c**) Bar chart of final OD_600_ after growth in the presence and absence of the AP1 TrpRS:tRNA_UCA_ pair. *E. coli* show reduced OD_600_ in stationary phase in strains MG1655 and DH10β, but not in BL21 or Ochre. (**d**) The AP1 TrpRS:tRNA_UCA_ pair is mutually orthogonal with the PylRS:tRNA^Pyl^_UCA_ but not the *M. jannaschii* tRNA^Tyr^. Heatmap shows average ELP-1xUGA-GFP / OD_600_ signal of n = 3 replicates. The ncAAs used for this experiment were *p*-acetylphenylalanine for the *M. jannaschii* pAzFRS, N^ε^-tert-butyloxycarbonyl-lysine (BocK) for the PylRS, and no exogenous ncAAs were used for the AP1 TrpRS.

We then evaluated the mutual orthogonality of the AP1 TrpRS:tRNA_UCA_ pair with an evolved *M. jannaschii* TyrRS-based OTS (pAzFRS) and with the chimeric Pyl OTS by measuring UGA suppression using all pairwise combinations of tRNAs_UCA_ and aaRSs (**Fig. 5d**).^9,10^ The AP1 tRNA_UCA_^Trp^ was orthogonal with respect to both aaRSs and the AP1 TrpRS was orthogonal to the tRNA_UCA_^Pyl^, showing that the AP1 TrpRS:tRNA_UCA_ pair is mutually orthogonal to the PylRS systems. However, the AP1 TrpRS nonspecifically recognizes *M. jannaschii* tRNA_UCA_^Tyr^ and therefore may require further engineering for orthogonality in dual ncAA incorporation applications when used alongside *M. jannaschii* TyrRS-based OTSs.

### Structure-based engineering of AP1 TrpRS

Finally, we aimed to engineer the AP1 TrpRS to incorporate an ncAA at the UGA codon. To inform our engineering efforts, we predicted a structure of the AP1 TrpRS: AP1 tRNA_UCA_ complex using AlphaFold3 (**Fig. S14**).^40^ The structural model shows a homodimeric TrpRS in which two tRNAs bridge individual subunits, consistent with a previously solved structure (PDBID: 2ake, **Fig. S14a**).^41^ Confidence metrics for the AP1 TrpRS (pLDDT = 89.8), the AP1 tRNA (pLDDT = 72.0), and the interfaces (ipTM = 0.88) are reasonably strong (**Fig. S14b**). Importantly, the AP1 TrpRS aligns well with a previously solved structure of the *E. coli* TrpRS bound to ATP and Trp (PDBID: 5v0i, RMSD = 2.24 Å) (**Fig. S14c**). As the *E. coli* TrpRS and other homologues have been engineered for ncAA incorporation, we hypothesized that structure-based engineering may successfully alter AP1 TrpRS substrate specificity.

We generated two AP1 TrpRS variants containing mutations that we hypothesized would enable selective incorporation of 5-hydroxytryptophan (5HTP, **Fig 6a**). Previous works have shown that mutations to the positions structurally aligning to T7, I145, and V147 in AP1 TrpRS (S8, V144, and V146 in *E. coli* TrpRS) enable selective incorporation of 5HTP in *E. coli* and *P. horikoshii* TrpRSs, and we further identified L6 and F40 in the AP1 TrpRS as positions in the tryptophan binding pocket that deviated from the *E. coli* TrpRS (F7 and C40, respectively) (**Fig. 6b**).^42,43^ We therefore designed AP1.1, containing mutations L6F, T7A, F40C, I145G, and V147C, as well as AP1.2, containing mutations L6F, T7A, F40C, I145A, and V147A. We then tested UGA suppression in the presence and absence of 5HTP with the hypothesis that selective variants which recognize 5HTP but discriminate against Trp should show improved ELP-1xUGA-GFP expression in the presence of 5HTP. Both variants showed improved expression of ELP-1xUGA-GFP in the presence of 5HTP (**Fig. 6c**). AP1.2 showed overall greater activity than AP1.1 based on ELP-1xUGA-GFP fluorescence data. Intermediate variants (T7A, I145G, V147C; T7A, I145A, V147A; and L6F, F40C) indicated no selectivity for 5HTP, showing that changing substrate specificity requires the combination of these mutations (**Fig. 6c**).

**Figure 6:**
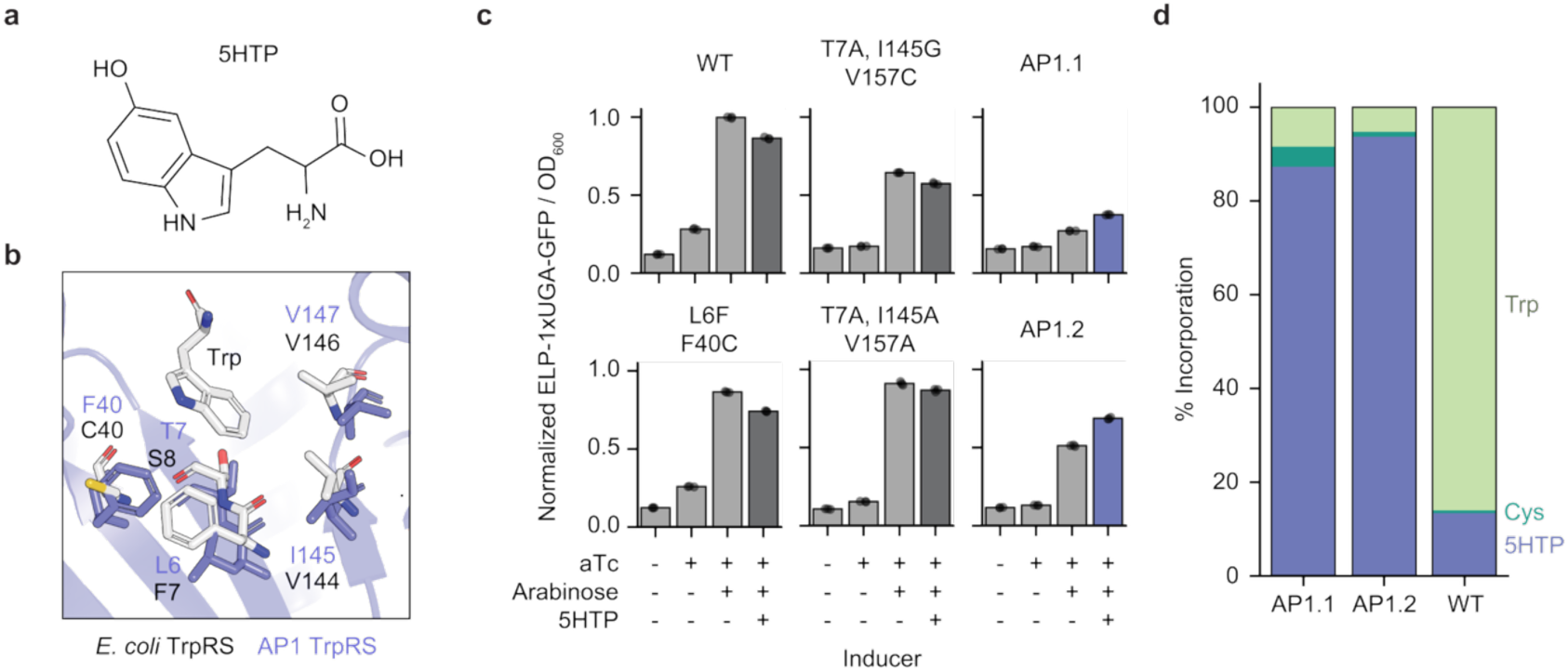
Structure-based engineering of AP1 TrpRS for 5HTP incorporation. (**a**) Chemical structure of 5HTP. (**b**) Residues involved in substrate specificity of AP1 TrpRS (blue) and *E. coli* TrpRS (gray). The structure of the *E. coli* TrpRS is from PDB ID 5v0i, and the structure of the AP1 TrpRS was predicting using AlphaFold3. AP1 TrpRS residues are labelled in blue, *E. coli* TrpRS residues are labelled in grey/black, and the AP1 TrpRS is represented as cartoon diagram in blue. (**c**) Bar chart of ELP-1xUGA-GFP / OD_600_ signal shows increased UGA suppression for AP1.1 and AP1.2 in the presence of 5HTP. X axis shows different induction conditions, where aTc is used to induce ELP-1xUGA-GFP, arabinose is used to induce AP1 TrpRS expression, and 5HTP is added to test for selectively. AP1.2 shows improved activity compared to AP1.1. Bars represent the average of n = 3 replicates and error bars represent one standard deviation. All data is normalized to the ELP-1xUGA-GFP / OD_600_ of WT AP1 TrpRS in the presence of aTc and arabinose. (**d**) MS-READ assays shows that both AP1.1 and AP1.2 selectively incorporate 5HTP compared to the WT AP1 TrpRS. Bars represent the average of n = 3 replicates.

Quantitative analysis of amino acid incorporation at UGA shows that AP1.1 and AP1.2 selectively incorporate 5HTP. We used mass spectrometry to measure incorporation events at UGA within the ELP peptide released after tryptic digestion of the ELP-1xUGA-GFP, which we call the mass spectrometry reporter for exact amino acid decoding (MS-READ, **Fig. 6d**).^44^ Strikingly, 5HTP accounts for greater than 90% of incorporation events at UGA when using AP1.1 and AP1.2 but only accounts for ∼12% of incorporation events at UGA when using the WT AP1 TrpRS. These results show that the AP1 TrpRS can be engineered to selectively incorporate ncAAs at the UGA codon, and we anticipate that improved selectivity and activity could be readily achievable with traditional directed evolution workflows using the recoded *Ochre* strain.^9,10,42^ Taken together, we engineered a novel AP1 OTS capable of 5HTP incorporation at the UGA codon.

## Discussion

In this work, we developed an integrated computational and experimental pipeline to discover OTSs from metagenomic datasets. The integration of these approaches is critical to our workflow; metagenomic bioprospecting allowed us to screen a natural source for suppressor tRNAs and aaRSs, high-throughput cell-free methods allowed us to rapidly identify orthogonal tRNAs and functional aaRS:tRNA pairs, and cellular aaRS engineering allowed us to characterize determinants of UGA suppression efficiency and identify AP1 TrpRS variants that are selective for 5HTP. We anticipate that this workflow will be a powerful strategy for genetic code expansion. By screening additional sequence databases, we hope to discover additional OTSs for genetic code expansion, which will be needed as genome engineering efforts in *E. coli* and other organisms construct alternative genetic codes with dedicated codons for ncAAs.^19,45–49^

Our study introduces three important advances for OTS discovery. First, the cell-free tRNA expression platform enables rapid, parallel assessment of hundreds of tRNAs, paving the way for large datasets that can be used to train machine learning models for rational tRNA engineering. While the platform does not yet distinguish inactive tRNAs from orthogonal ones, integrating predictive algorithms to rescue inactive variants could improve dataset quality.^43^ Second, we harness natural diversity in bacteriophages as a resource for genetic code expansion.^17,18^ Screening natural suppressor tRNAs avoids pitfalls associated with converting sense codon tRNAs into suppressors, which can disrupt aaRS recognition or compromise orthogonality.^11,43^ Third, our strategy enables discovery of rare genetic code deviations, exemplified by tRNA_CUA_-151, which mediates an unusual UAG→Ala reassignment. Although its biological relevance remains unclear, such findings challenge assumptions about the structural rigidity of the genetic code.

The AP1 TrpRS:tRNA^Trp^_UCA_ is an efficient and naturally UGA-suppressing OTS that functions *in vitro* and *in vivo*. The AP1 TrpRS:tRNA^Trp^_UCA_ adds to the growing repertoire of OTSs used to incorporate Trp derivatives in *E. coli*, such as the *Saccharomyces cerevisiae* TrpRS:tRNA^Trp^_CUA_, the chimeric PylRS:tRNA^Pyl^_CUA_ systems, the *E. coli* TrpRS:tRNA^Trp^_UCA_ systems used in ATMW strains, and others.^8,42,43,50,51^ The AP1 OTS, however, consolidates multiple desirable properties, such as efficient UGA suppression, portability into common *E. coli* strains, and mutual orthogonality with other OTSs. We also highlight the synergy between genomically recoded strains like *Ochre* and OTSs specific for UGA. This synergy establishes new paradigms to use genomically recoded organisms to screen and evolve OTSs for codons that have been previously inaccessible.

Looking forward, we anticipate that our platform for identifying, characterizing, and engineering tRNAs and aaRSs will be used to elucidate fundamental principles underpinning protein biosynthesis and recoding genomes. Along with continuing advances in synthetic biology to liberate codons,^19,45–49^ our cell-free workflow and AP1 TrpRS:tRNA^Trp^_UCA_ pair is poised to impact efforts to expand the genetic code.

## Methods

### Bioinformatic identification of tRNAs

To identify potential suppressor tRNAs from metagenomes we used tRNAScanSE (v. 2.0.9)^20^ on 736 metagenomes,^18^ previously analyzed for stop-codon recoding, and 400 publicly available phage contigs. We used tRNAScanSE preset parameters relevant for phages (-B; -O; -G) and dereplicated the final results. Predicted suppressor tRNAs were determined to be low confidence if the tRNAScanSE score was below 30 or their length was greater than 100 nucleotides.

Phylogenetic trees of tRNAs were built with IQ-TREE (v. 1.6.12) and parameters with 1000 ultrafast bootstraps and 5000 maximum iterations with the K3P+I+G4 model chosen through the Bayesian Information Criterion with the integrated model finder.^52^

### Bioinformatic identification of aaRSs

To identify aaRSs possibly involved in functional OTSs we searched for potential hosts by identifying genomes from the same sequencing projects as the phages that have CRISPR-Cas systems. We extracted the CRISPR spacers and searched for their presence in the phage genomes with BLAST.^53^ After identifying hosts, we used HMMER 3.3 with TIGRFAM HMMs to identify aaRSs.^54^

### In vitro transcription of RNase-P-based tRNA templates

DNA templates encoding the tRNAs were purchased as eBlocks from IDT in a linear template:

~~~
tttcgccacctctgacttgagcgtcgatttttgtgatgctcgtcaggggggcggagcctatggaaaaacgccagcaacgcgatcccgcgaaattaatacgactcactatagggagaccacaacggtttccctctaga[tR NA]gtcgaccggctgctaacaaagcccgaaaggaagctgagttggctgctgccaccgctgagcaataactagcataaccccttggggcctctaaacgg
~~~

where [tRNA] denotes the tRNA sequence as listed in **Table S1**. Templates were amplified using Q5 DNA Polymerase (NEB) following manufacturer’s protocols using a T_m_ of 50 °C, with a T7-specific forward primer (5’-TAATACGACTCACTATAGGG-3’) and a tRNA-specific reverse primer (**Table S3**). Reverse primers were purchased from IDT with a C2’-methoxy modification on the penultimate 5’ base to reduce 3’ non-templated nucleotide addition by T7 RNA polymerase.^55^ To verify amplification, 2 μL of PCR product was mixed with 1 μL of 6X Gel Loading Dye (NEB) and 3 μL of nuclease free (NF) water (Ambion) and loaded into a 1% w/v agarose gel containing SYBR Safe Dye (Apex Bio). The gel was run in 1X TAE buffer at 120V for 20 minutes and imaged using a Bio-Rad Gel Doc XR+.

tRNA templates were *in vitro* transcribed using HiScribe T7 RNA Synthesis Kit (NEB). 1.8 μL of crude PCR product was mixed with 0.75 μL each of 10X Buffer, 100 mM NTPs, T7 RNA Polymerase, and 3.7 μL NF water. Reactions were set up at 0.75X concentration as recommended by NEB for products < 0.3 kb. Reactions were incubated for 4-6 hours at 37 °C and stored at −20 °C or −80 °C. Crude *in vitro* transcription products were either analyzed by denaturing PAGE or used in CFPS reactions.

### Denaturing PAGE

12% Urea-PAGE gels were cast using the SequaGel UreaGel 19:1 Denaturing Gel System (National Diagnostics) using appropriate spacers (1 mm for analytical gels, 3 mm for preparative gels) and an appropriate comb. Gels were allowed to polymerize for 1 - 2 hours at room temperature. *In vitro* transcription products or cleaved tRNAs (see tRNA aminoacylation assays section) were mixed with an equal volume of 2X RNA Loading Dye (8 M Urea, 2 mM Tris pH 7.5, 2 mM EDTA, 0.004% Bromophenol Blue), denatured at 70 °C for 15 minutes, and cooled on ice for 5 minutes. Samples were loaded onto the gel, along with a low range ssRNA ladder (NEB). The gel was run in 1X TBE buffer at 230 V for 2.5 hours at 4 °C or until the dye front had reached the bottom of the gel. For imaging, gels were stained with 1X SYBR Gold (Thermo Fisher) for 10 min with gentle shaking.

### CFE reaction setup

CFE reactions were set up in 5 μL reactions using previously described conditions.^56^ Reactions consisted of 8 mM magnesium glutamate, 10 mM ammonium glutamate, 130 mM potassium glutamate, 1.2 mM ATP, 0.85 mM GTP, 0.85 mM UTP, 0.85 mM CTP, 0.03 mg/mL folinic acid, 0.17 mg/mL total *E. coli* tRNA, 0.4 mM NAD, 0.27 mM CoA, 4 mM oxalic acid, 1 mM putrescine, 1.5 mM spermidine, 57 mM HEPES pH 7.2, 2 mM total amino acids, 0.03 M phosphoenolpyruvate, 5 ng/μL plasmid DNA in pJL1 backbone (https://www.addgene.org/102634/, purified by Zymo DNA Midiprep),^57^ 30% v/v cell extract, and water. To express tRNAs from crude IVT reactions, 10% v/v crude tRNA product was added. For the *M. jannaschii* tRNA^Tyr^, the BpyRS was added at 1 mg/mL and Bpy was added at 1 mM. For tRNAs^Pyl^, the corresponding PylRSs were added at 1 mg/mL and AzK was added at 1 mM. The *M. barkeri* tRNA^Pyl^ was paired with the IPYE chimeric PylRS,^9^ the *M. Alvus* tRNA^Pyl^ was paired with the *M. alvus* PylRS,^58^ and the ^A^17 ^V^C10 *Int* tRNA^Pyl^ was paired with the *Lum1* PylRS.^29^ To screen aaRSs for activity with orthogonal tRNAs, we supplemented midiprepped pJL1 plasmids encoding each aaRS into CFE reactions at a final concentration of 5 ng/µL. AP1 TrpRS and HF2 TrpRS were purified and added at concentrations of 1 mg/mL. aaRS purification protocols are described below. CFPS reactions were incubated at 37 °C for UAG suppression or at 30 °C for UAA and UGA suppression.^27^

### Cell extract preparation

759.T7, derived from 759, was used as the chassis strain for cell extract preparation for all experiments because it has been optimized for efficient ncAA incorporation at the UAG codon. 759.T7 was streaked onto an LB-agar plate from a glycerol stock and incubated at 34 °C for ∼ 20 hrs. A single colony was inoculated into 100 mL of LB and incubated at 34 °C for ∼ 20 hrs with 220 rpm shaking. The next day, 5 L of 2xYTPG (16 g/L tryptone, 10 g/L yeast extract, 5 g/L NaCl, 7 g/L dibasic potassium phosphate, 3 g/L monobasic potassium phosphate, 18 g/L glucose) was inoculated at OD_600_ = 0.075 with the overnight culture and grown at 34 °C with 220 rpm shaking. At OD_600_ = 0.6, IPTG was added to a final concentration of 1 mM to induce expression of T7 RNA polymerase. Cells were grown until OD_600_ = 3.0. Cells were pelleted by centrifugation at 5k x g for 10 min at 4 °C and were washed three times with cold S30 buffer (10 mM Tris-Acetate pH 8.2, 14 mM Mg Acetate, 60 mM K Acetate, 2 mM DTT). Cells were flash frozen in liquid nitrogen and stored at −80 °C.

Extract preparation follows previously developed approaches.^59,60^ Cells were thawed on ice and resuspended in 0.8 mL S30 buffer / g cells. Cells were sonicated in 1.4 mL aliquots using three 45 sec on / 59 sec off cycles at 50% amplitude for a total of 950 J in a Q125 Sonicator (Qsonica). Lysed cells were centrifuged for 10 min at 12 k x g at 4 °C. Supernatant was removed and a run-off reaction was performed by incubating the supernatant at 37 °C for 1 hour. Extracts were centrifuged for 10 min at 10 k x g at 4 °C to remove insoluble components. Supernatant was transferred to a Slide-a-Lyzer (10 kDa MWCo) and dialyzed into 200X volumes of S30 buffer for three hours. Clarified extract was isolated by centrifuging for 10 min at 10 k x g at 4 °C, put into single use aliquots, flash-frozen in liquid nitrogen, and stored at −80 °C.

### sfGFP quantification

WT sfGFP and 216X-sfGFP fluorescence was measured using a Bio-Rad Synergy 2 plate reader. Yields were calculated using a standard curve of sfGFP fluorescence vs. sfGFP yields as measured by ^14^C-leucine scintillation counts. WT sfGFP containing ^14^C-leucine was synthesized by adding ^14^C-leucine (Perkin Elmer) at a concentration of 10 μM in CFPS. An equal volume of 0.5 M KOH was added to CFPS reactions to hydrolyze ^14^C-leucine-acylated tRNA. Samples were pipetted onto two Filtermat A (Perkin Elmer) fiberglass paper sheets. After drying, one filtermat was washed 3X with 5% w/v trichloroacetic acid and 1X with 100% ethanol. Meltilex A (Perkin Elmer) was applied to both sheets, and scintillation counts were measured using the MicroBeta^2^. Soluble yields were calculated as described previously.^61,62^ Dilutions were made in 1X PBS and a standard curve was built using linear regression in Microsoft Excel.

### aaRS purification

Overexpression vectors containing aaRSs with C-terminal 10X His tags were either purchased from Twist Biosciences in the pET.BCS backbone or were cloned in-house. If cloned, aaRS sequences were purchased as gBlocks from IDT containing appropriate overhangs for Gibson Assembly into a pET.BCS overexpression vector. Linearized pET.BCS vector was synthesized by PCR using Q5 DNA Polymerase following the manufacturer’s instructions using forward primer (5’-GCAGTAGTGGTCATCATC-3’) and reverse primer (5’-atgTCCTCCTTATGTGTG-3’) at a T_m_ of 59 °C. PCR amplification was confirmed by agarose gel electrophoresis as described previously. PCR products were column purified using Zymo DNA Clean and Concentrate, resuspended in 1X CutSmart Buffer, and digested with DpnI (NEB) overnight at 37 °C. Gibson Assembly reactions were set up with 25 ng of backbone and a 3X molar excess of the insert, along with 75 mM Tris-HCl pH 7.5, 7.5 mM MgCl_2_, 0.15 mM dNTPs, 7.5 mM DTT, 0.75 mM NAD, 0.004 U/μL T5 Exonuclease, 0.025 U/μL Phusion Polymerase, 4 U/μL Taq DNA Ligase, and 3.125 μg/mL ET SSB. Reactions were incubated at 50 °C for one hour. The entire reaction was transformed into chemically competent NEB 5α cells following the manufacturer’s protocol, plated onto LB-Carb[100] agar plates, and incubated overnight at 37 °C. Single colonies were inoculated into 5 mL of LB-Carb[100] and grown overnight at 37 °C with 250 RPM shaking. Plasmid DNA was purified using Zymo Miniprep Kits and sequence confirmed.

Plasmids encoding aaRSs of interest were transformed into BL21 (DE3) Star following manufacturer’s protocols and plated onto LB-Carb[100]. The next day, a single colony was inoculated into 3 mL of LB-Carb[100] and grown at 37 °C until saturated. 250 mL of Overnight Express TB Media (Millipore) was prepared by mixing 15 g of powder, 2.5 mL of glycerol, and 250 mL of water and was sterilized by microwaving until bubbles started to appear. After cooling, 250 μL of Carb[100] was added and mixed. 250 μL of saturated culture was mixed into the Overnight Express TB Media and incubated at 37 °C overnight with 250 rpm shaking.

Cells were pelleted by centrifugation at 5k x g for 10 min in an Avanti J-25 centrifuge and washed with Buffer 1 (300 mM NaCl, 50 mM monobasic sodium phosphate pH 8.0) containing 10 mM imidazole pH 8.0. Cells were resuspended in Buffer 1 + 10 mM imidazole pH 8.0 by vortexing and supplemented with Benzonase (Thermo Fisher). Cells were pulled through an 18-gauge syringe needle and lysed by homogenization at ∼20,000 PSI in an Avestin B3 homogenizer. Cellular debris was pelleted by centrifugation at 20k x g. Supernatant was added to pre-equilibrated Ni-NTA resin (Qiagen) and incubated with end-over-end shaking at 4 °C for 1 hour. Supernatant was removed by centrifugation, and the resin was washed five times with Buffer 1 + 20 mM imidazole pH 8.0. Resin was then packed into a gravity column. Proteins were eluted with 20 mL of Buffer 1 + 0.5 M imidazole pH 8.0 in 1 mL fractions. Protein-containing fractions were identified by measuring A280 on nanodrop and analyzed by SDS-PAGE. 1 uL of eluted protein was mixed with 3.75 μL 4X LDS Sample Buffer, 1.5 μL 1M DTT, and water to 15 μL. Samples were denatured at 95 °C for 10 minutes, and 10 μL of sample were loaded onto a 4-12% Bis-Tris NuPAGE gel (Invitrogen). Gel was run in 1X MES Buffer at 180 V for 45 minutes. Gels were stained in AcquaStain Protein Gel (Bulldog Bio) for 15 minutes and then imaged to confirm protein size and purity. aaRS-containing fractions were pooled and dialyzed into Buffer 1 + 40% v/v glycerol, with three buffer changes. After dialysis, proteins were quantified by Nanodrop (using molecular weights and exctinction coefficients calculated by ExPasy ProtParam), put into single-use aliquots, flash-frozen in liquid nitrogen, and stored in −80C.

### Intact protein ESI-MS

216X-sfGFP was synthesized in 50 μL CFPS reactions and was purified with Strep-Tactin XT Resin and spin columns as recommended by the manufacturer (IBA Life Sciences). Eluted proteins were buffer exchanged into 100 mM ammonium acetate using Amicon Ultra 0.5 mL Centrifugal Filters (10 kDa MWCO). Protein purity was confirmed by SDS-PAGE.

Samples were injected on a 1200 HPLC System (Agilent Technologies Inc., Santa Clara, California, USA) onto a Thermo Hypersil-C18 column (3.0 μm, 30 × 2.1 mm) for reverse-phase separation which was maintained at 35 °C with a constant flow rate at 0.400 ml/min, using a gradient of mobile phase A (water, 0.1 % formic acid (v/v)) and mobile phase B (acetonitrile, 0.1% formic acid (v/v)). The gradient program was as follows: 0 – 0.5 min, 1%B; 0.5 – 5 min, 1 – 100%B; 5 – 7.25 min, 100%B; 7.25-7.5 mins, 100 – 1%B; 7.5-10 min, 1%B. “MS-Only”, positive ion mode acquisition was utilized on an Agilent 6230 time-of-flight mass spectrometer equipped Electrospray ionization source (Agilent Technologies Inc., Santa Clara, California, USA). The source conditions were as follows: Gas Temperature, 320 °C; Drying Gas flow, 5 L/ min; Nebulizer, 20 psi; VCap, 4500 V; Fragmentor, 210 V; Skimmer, 65 V; and Oct 1 RF, 750 V. The acquisition rate in MS-Only mode was 3 spectra/second, utilizing m/z 922.009798 as reference masses. Data was plotted using a custom python script.

### tRNA aminoacylation assays

To prepare for tRNA aminoacylation assays, tRNAs were first synthesized in scaled-up *in vitro* transcription reactions from a DNA template containing a 5’ hammerhead ribozyme construct.^22^ DNA templates were amplified using PCR and purified using the Zymo DNA Clean and Concentrate kit to ensure efficient transcription. After IVT, crude IVT products were cleaved by adding 5X volumes of NF water and incubating at 60 °C for between 2-8 hours. 0.1X volumes of 3M NaCl and 2.5X volumes of 100% ice-cold ethanol were added to cleavage reactions and incubated at −20 °C for at least 2 hours to precipitate products. Crude product was pelleted by centrifugation at 21k x g for 10 min at 4 °C. Products were resuspended in 200 μL 1X RNA Loading Dye, denatured by incubating at 70 °C for 10 min, loaded onto a denaturing Urea-PAGE gel (made with 3 mm spacers), and run in 1X TBE at 230V at 4 °C for 2.5 hours. Bands containing cleaved tRNA were visualized by UV shadowing, excised from the gel, crushed using a clean pestle, and passive eluted into 0.3 M NaCl overnight with end-over-end shaking at 4 °C. Supernatant from passive elution was isolated by centrifugation and was ethanol precipitated. tRNA concentrations were quantified by Nanodrop 2000c.

For the AP1 and HF2 aaRSs, aminoacylation reactions were set up by in 30 μL reactions consisting of 100 mM HEPES pH 7.4, 4 mM DTT, 10 mM MgCl_2_, 10 mM ATP, 7.5 nM aaRS, 2.5 mM amino acid, 4 U/mL PPIase (NEB), and 15.6 μM tRNA candidate or 106 μM total *E. coli* tRNA.^63^ We chose a concentration of 15.6 μM tRNA because it approximates *in vivo* concentrations of tRNA^Trp^ in *E. coli* cells (943 tRNA^Trp^ molecules/cell^64^ assuming a cell volume of 1 fL (https://ecmdb.ca/e_coli_stats)). A 10-fold excess was chosen to ensure observable signal in LC-MS and because *in vivo* tRNA expression is typically done with a pEVOL vector containing a p15A origin of replication, which has a copy number ∼ 10. 106 μM total *E. coli* tRNA was calculated by assuming 64,274 tRNA molecules/cell^64^ and a cell volume of 1 fL (https://ecmdb.ca/e_coli_stats). For the metagenomically-identified GlnRS and TyrRS, aminoacylation reactions were set up in 30 μL reactions mimicking CFPS conditions, which consisted of 130 mM potassium glutamate, 1.2 mM ATP, 0.85 mM GTP, 0.85 mM UTP, 0.85 mM CTP, 0.03 mg/mL folinic acid, 0.4 mM NAD, 0.27 mM CoA, 4 mM oxalic acid, 1 mM putrescine, 1.5 mM spermidine, 57 mM HEPES pH 7.2, 2 mM total amino acids, 0.03 M phosphoenolpyruvate, 15.6 μM tRNA or 106 μM *E. coli* tRNA, 7.5 nM aaRS, and 30% v/v cell extract that was filtered through an Amicon Ultra 0.5 mL Centrifugal Filters (3 kDa MWCO) to remove any aaRSs and tRNAs. The GlnRS and TyrRS were found to be inactive in the previous conditions. For kinetics, 5 μL at each timepoint were removed and quenched with 1.1X volumes of RNase A solution (200 mM sodium acetate pH 5.2, 1.5 U/μL RNase A (NEB)) and incubated for 5 min at room temperature. Reactions were precipitated with 0.1X volumes of 50% w/v trichloroacetic acid and incubated at −80 °C for at least 30 min. Samples were spun down at 21,000 x g for 10 min at 4 °C, and supernatant was transferred to autosampler vials and analyzed by LC-MS.

Samples were injected on a 1290 Infinity II UHPLC System (Agilent Technologies Inc., Santa Clara, California, USA) onto a Poroshell 120 EC-C18 column (1.9 μm, 50 × 2.1 mm) (Agilent Technologies Inc., Santa Clara, California, USA) for reverse-phase separation which was maintained at 30 °C with a constant flow rate at 0.500 ml/min, using a gradient of mobile phase A (water, 0.1 % formic acid (v/v)) and mobile phase B (acetonitrile, 0.1% formic acid (v/v)). The gradient program was as follows: 0 – 1 min, 2%B; 1 – 5 min, 2 – 40%B; 5 – 6 min, 40 – 99%B; 6 – 8 mins, 99%B; 8 – 8.10 min, 99 – 2%B; 8.10 – 14 min, 2%B. “MS-Only”, positive ion mode acquisition was utilized on an Agilent 6545 quadrupole time-of-flight mass spectrometer equipped with a JetStream ionization source (Agilent Technologies Inc., Santa Clara, California, USA). The source conditions were as follows: Gas Temperature, 300 °C; Drying Gas flow, 12 L/ min; Nebulizer, 45 psi; Sheath Gas Temperature, 350 °C; Sheath Gas Flow, 12 L/ min; VCap, 3500 V; Fragmentor, 110 V; Skimmer, 65 V; and Oct 1 RF, 750 V. The acquisition rate in MS-Only mode was 3 spectra/second, utilizing *m/z* 121.050873 and *m/z* 922.009798 as reference masses.

### Characterizing sfGFP mutations

Mutations at T216 in sfGFP were installed using a previously described method.^65^ 20 pairs of mutagenic primers were designed for site-directed mutagenesis of sfGFP T216 and ordered from IDT. All following T_m_s are calculated by Benchling. The primers were designed to overlap with a T_m_ of 40-45°C. The forward primer containing the mutation was designed to have a T_m_ of 60-62 °C and the reverse primer was designed to have a T_m_ of 58 °C. Linear templates containing the desired mutation were amplified by Q5 DNA Polymerase in 10 μL PCR reactions using a touchdown PCR, starting at T_m_ of 72 °C and stepping down −1 °C/cycle until a T_m_ of 62 °C was reached. Cycles were repeated for a total of 25 cycles.

PCR products were digested by adding 1 μL of Dpn1 to the crude PCR reactions and were incubated for 2 hrs at 37 °C. Reactions were diluted 4X, and 1 μL was added into a 4 μL Gibson Assembly Reaction, as described previously. Gibson Assembly reactions were incubated at 50 °C for one hour. Reactions were diluted 10X and used in a second PCR reaction to prepare linear expression templates using Q5 DNA Polymerase following the manufacturer’s protocol using forward primer (5’-ctgagatacctacagcgtgagc-3’) and reverse primer (5’-cgtcactcatggtgatttctcacttg-3’).

Linear templates were then added into CFPS reactions along with 1 μM GamS to protect linear templates and incubated at 37 °C. sfGFP was quantified as previously described.

### RNase P cleavage assays

*In vitro* transcription reactions were set up as previously described. 1 μL of IVT reaction was diluted with 80 μL NF water and was split into 2 x 9 μL aliquots. One aliquot was treated with 1 μL of cell extract (diluted as necessary), and the other was treated with 1 μL of water. Reactions were incubated at 37 °C for one hour and were quenched by addition of RNA Loading Dye. Denaturing Urea PAGE was run as previously described.

### Structure predictions and analysis

tRNA secondary structure predictions were done using R2DT (https://rnacentral.org/r2dt).^66^ All tRNA structures except tRNA_UCA_-29 were predicted using the default settings. tRNA_UCA_-29 structure was predicted using the Constrained Folding setting on the Full Molecule.

The AP1 TrpRS:AP1 tRNA_UCA_ complex was predicted using AlphaFold3 (alphafoldserver.com). Two copies of the tRNA, protein, and ATP were used as inputs. AlphaFold3 was run 15 times with different starting seeds and the model with the best ipTM and pTM scores was selected for analysis. Average atom pLDDT was calculated using a custom Python script.

### In vivo characterization of AP1 OTS

Unless otherwise stated, all cultures were grown in Lysogeny Broth Lennox (LB) composed of 10 g/L bacto tryptone, 5 g/L yeast extract, and 5 g/L NaCl (Sigma-Aldrich # L3022). LB agar plates were composed of LB plus 15 g/L bacto agar. N^ε^-Boc-L-lysine (BocK) was purchased from Chem-Impex (#00363) and dissolved in LB to a final concentration of 10 mM. p-acetylphenylalanine (pAcF) was purchased from Chem-Impex (#24756), dissolved in sterile water to a concentration of 50 mM, filter-sterilized with a 0.22 µm filter, and used at a final concentration of 1 mM. 5HTP was purchased from Chem-Impex (#00607), dissolved in 15% DMSO, 0.2 M NaOH to a concentration of 100 mM, filter-sterilized with a 0.22 µm filter, and used at a final concentration of 1 mM.

Plasmids for *in vivo* expression of the AP1 OTS were cloned using Golden Gate Assembly. Gene fragments for the aaRS and tRNA were synthesized by Twist Bioscience. Plasmids were sequence verified by whole-plasmid sequencing (Plasmidsaurus or Quintarabio). All cloning was made in Mach1(Thermofisher #C862003) or DH10β (NEB #C3019H) using standard protocols.

Strains were transformed with OTS-reporter plasmids by standard electroporation protocols. Electroporated strains were recovered in 2 mL SOC (2% tryptone, 0.5% yeast extract, 10 mM NaCl, 2.5 mM KCl, 10 mM MgCl_2_, 10 mM MgSO_4_, and 20 mM glucose) for at least 2 hours before plating onto LB-agar plates with kanamycin (50 µg/mL) and incubated at 37 °C overnight. Single colonies from each plate were picked and grown in 800 µL LB (10 g/L bacto tryptone, 5 g/L yeast extract) supplemented with 50 µg/mL kanamycin in a 96 deep-well plate sealed with a Breathe-Easy film (Sigma-Aldrich) and incubated at 37 °C with shaking at 220 rpm for 20-24 hours. After overnight growth the cultures were back-diluted 1:50 onto a clear-bottom black 96-well plate (Costar) in a total of 150 µL of LB supplemented with kanamycin (50 µg/mL), aTc (100 ng/mL), L-arabinose (0.05 % w/v). Cell growth (absorbance at OD_600_) and GFP fluorescence (excitation 485 nm, emission 525, gain 70, bottom measurement) were measured in a BioTek Synergy H1 plate reader (Agilent) for 24 hours at 10 minutes intervals with linear shaking. Data was analyzed with a custom Python script. GFP fluorescence was taken from the maximum value after 10 hours and plotted as bar plots, first normalized by OD_600_ and then to CGC native codon control construct GFP fluorescence.

For the mutual orthogonality OTS experiments, 1 mM pAcF and 10 mM BocK were added to cultures containing the pAzFRS or the PylRS, respectively, in addition to other inducers. For the 5HTP incorporation assay, 1 mM 5HTP was added in addition to other inducers.

### Doubling time analysis

Kinetic growth (OD_600_) curves were obtained via monitoring strain growth within a BioTek Synergy HT1 plate reader from the expression experiments. The absorbance obtained by the BioTek Synergy H1 plate reader was calibrated to OD_600_ (absorbance at 600 nm through a 1 cm path length) using a standard curve *y* = 2.082*x* + 1.123*x*^2^. The OD_600_ of an overnight LB culture of MG1655 was measured by a Biochrom Libra S4 Spectrophotometer at 600 nm wavelength in a Semi-micro cuvette (1 cm pathway) after 1:10 dilution of culture into 1mL LB. To generate the calibration curve, a series of cultures with OD_600_ ranging from 0 to 6 were prepared by diluting in LB media. These cultures were then measured by the BioTek Synergy H1 plate reader, with the same settings as growth cultures. The average values were then fitted to a polynomial standard curve for recalibration. The effects of media evaporation in plate wells are not considered. The recalibrated growth curve was used to calculate the doubling time and MaxOD using a custom Python script. Linear fitting of *log*_2_*OD*_600_ was performed using a sliding window method, where the window size is 50 minutes in the early log phase.^67^ The doubling time was calculated as the reciprocal of the slope. MaxOD was obtained within the 24-h growth period.

### MS-READ Analysis

Single colonies of Ochre containing the AP1-OTS-ELP-TGA-GFP plasmid were inoculated in 2 mL LB with 50 µg/mL kanamycin in a 14 mL falcon tube overnight at 37 °C with shaking at 220 rpm. After overnight growth the cultures were diluted 1:100 into a 250 mL baffled flask with 50 mL LB-kan supplemented with 1 mM 5HTP, 100 ng/mL aTc, and 0.05 % w/v L-arabinose. The cultures were grown at 37°C overnight after which the cells were harvested by centrifugation at 3,200 g for 20 minutes in a 50 mL centrifuge tube and stored at −20 °C until protein purification.

Frozen *E. coli* cell pellets (approximately 1 g each) were thawed on ice, and lysed by chemical lysis in by resuspension in 5 mL of lysis buffer consisting of Bugbuster reagent (Sigma-Aldrich #70921) supplemented with 1X PBS, 2.5 units/mL benzonase (Sigma-Aldrich #E8263), 20 mM imidazole, 1 tablet pr 25 mL cOmplete™, Mini, EDTA-free Protease Inhibitor Cocktail (Sigma-Aldrich #11836170001).The resuspended pellets were incubated on a rotator for 20 minutes at room temperature, followed by centrifugation at 3,200 g in swinging bucket rotor for 20 minutes to pellet cellular debris. The resulting supernatant was filter-sterilized using 10 mL syringes into 15 mL conical tubes. For each purification, 200 µL bed volume (400 μL slurry volume) of Ni-NTA agarose resin was used. To equilibrate the column 400 µL of slurry was centrifuged in a 1.7 mL tube at 11,000 for 1 minute. The supernatant was removed, and the resin pellet was resuspended in 1 mL of 1X PBS, centrifuged at 11,000 for 1 minute, and the supernatant discarded. The equilibrated resin was then resuspended in 1 mL of the clarified lysate and transferred to the 15 mL lysate tubes for a 20-minute incubation on a rotator at room temperature. The protein-resin mixture was applied to pierce spin columns (Thermo-Fischer #69705) using a syringe. Columns were washed with 4 mL of wash buffer (PBS, 20 mM imidazole), and bound proteins were eluted with 4 mL of elution buffer (PBS, 500 mM imidazole). The eluate was buffer exchanged and concentrated with PBS using Amicon 10 kDa molecular weight cut-off spin columns (Sigma-Aldrich, #UFC8010) following the manufacturer’s protocol. Protein concentration was determined by nanodrop spectrophotometry, and purified samples were stored at −20° C until further use.

Affinity purified, buffer exchanged ELP-GFP reporter protein were digested and analyzed by mass spectrometry by the MS & Proteomics Resource at Yale University. MS data were searched using MaxQuant version 1.6.10.43 with Deamidation (NQ), Oxidation (MW) as variable modifications and Carbamidomethyl (C) as a fixed modification with up to 3 missed cleavages, 5 AA minimum length, and 1% FDR against a modified Uniprot *E. coli* database containing custom MS-READ reporter proteins. MS-READ search results were analyzed using MaxQuant and Perseus version 1.6.2.2.

## Supporting information

tRNA orthogonality data

## Acknowledgements

We would like to thank Dr Jessica Willi, Dr. Antje Kruger, and Dr. Camila Kofman for discussions about tRNAs and the genetic code, Dr. Steve Fleming for help with *in vitro* transcription and tRNA purification, Dr. Grant Landwehr for help with site saturation mutagenesis, Kathryn Myers for experimental assistance, Prof. Julius Lucks for his insight into RNase P function, Dr. Fernando Tobias for his assistance regarding LC-MS methods, and Kebron Gurara and Prof. Jesse Rinehart for discussions about OTSs and assistance with MS-READ. We would also like to thank Prof. Ashty Karim for careful editing of the manuscript. This work made use of the IMSERC MS facility at Northwestern University, which has received support from the Soft and Hybrid Nanotechnology Experimental (SHyNE) Resource (NSF ECCS-2025633), and Northwestern University. We also thank the MS & Proteomics Resource at Yale University for providing the necessary mass spectrometers and the accompany biotechnology tools funded in part by the Yale School of Medicine and by the Office of The Director, National Institutes of Health (S10OD02365101A1, S10OD019967, and S10OD018034). The funders had no role in study design, data collection and analysis, decision to publish, or preparation of the manuscript.

## Author Contributions

**Conceptualization**: K.S, M.C.J; **Methodology**: K.S., M.T.A.N.; **Software**: P.I.P.;**Validation**: K.S., M.T.A.N.; **Formal analysis**: K.S., M.T.A.N., P.I.P.; **Investigation**: K.S., M.T.A.N., P.I.P.; **Resources**: P.I.P.; **Data curation**: P.I.P.; **Writing**: K.S., M.C.J.; **Visualization**: K.S., M.C.J.; **Supervision**: M.C.J., J.F.B., F.J.I.; **Project administration**: M.C.J., J.F.B., F.J.I.; **Funding acquisition**: M.C.J., J.F.B., F.J.I.

## Data and materials availability

All data is available upon request.

## Supplementary Information

### Supplementary Figures

**Figure S1:**
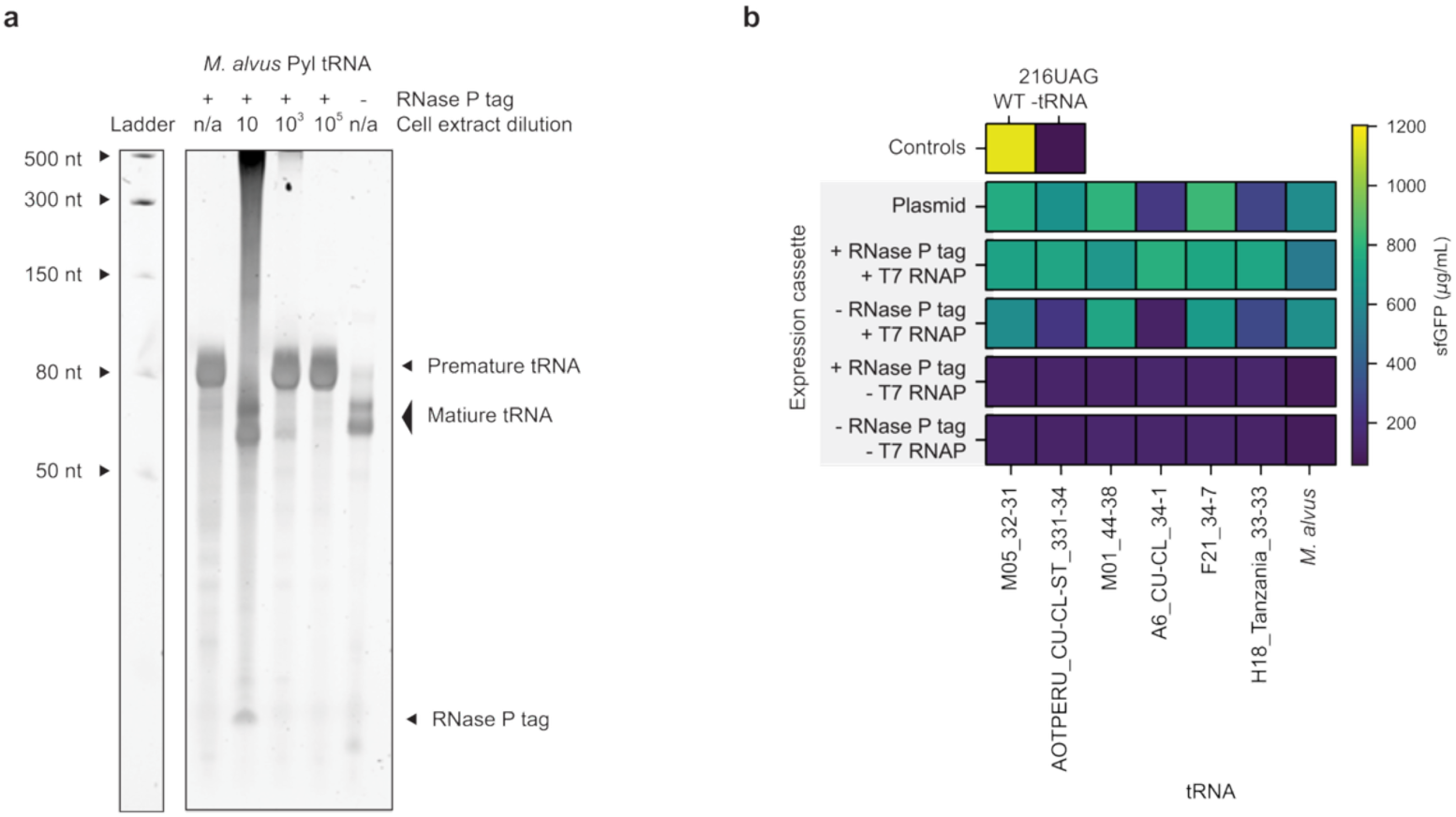
A cell-free RNase-P based tRNA expression platform efficiently transcribes and matures functional tRNAs. (**a**) The *M. alvus* tRNA^Pyl^_CUA_ can be cleaved into a mature tRNA by the action of endogenous RNase P in cell extracts. *In vitro* transcribed *M. alvus* tRNA^Pyl^_CUA_, with and without the RNase P tag, was incubated the *E. coli* S30 extracts and then analyzed by denaturing urea PAGE to visualize cleavage. We note that the mature tRNA band in the presence of cell extracts consists of a mixture of endogenous *E. coli* tRNAs and the *M. alvus* tRNA^Pyl^_CUA_ due to the presence endogenous tRNAs in *E. coli* S30 extracts. A representative gel from two experiments is shown. (**b**) Evaluation of a panel of metagenomically-identified tRNAs_CUA_ shows efficient expression of all tRNAs when using the RNase P tag compared to plasmid-based expression and constructs lacking the RNase P tag. *In vitro* transcribed *M. alvus* tRNA^Pyl^_CUA_, using a variety of experimental conditions, were supplemented into CFE reactions expressing 216UAG-sfGFP. Average expression from n = 3 replicates is plotted in the heatmap.

**Figure S2:**
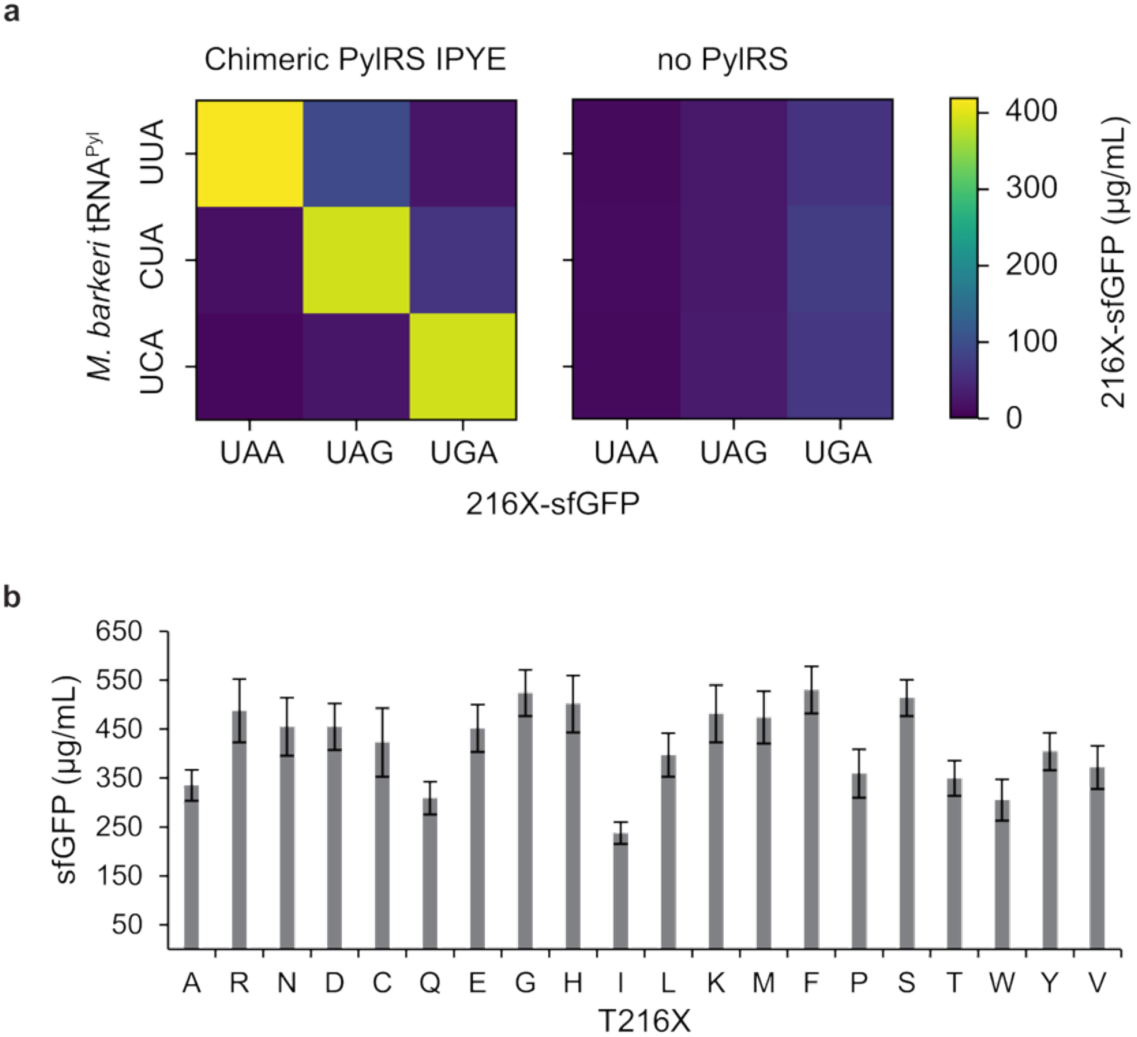
tRNA orthogonality and activity can be robustly detected in CFE workflows. (**a**) Cell free reactions cannot synthesize 216UAG-sfGFP in the presence of an orthogonal tRNA (no PylRS). Addition of a cognate aaRS (chimeric PylRS IPYE) restores suppression activity. tRNAs were expressed in cells prior to CFE. Y-axis is labelled with anticodon of tRNA and X-axis is labelled with the premature stop codon in 216X-sfGFP. Data shows the average of n = 3 replicates. (**b**) Position T216 of sfGFP is robust to all 20 canonical amino acid mutations, indicating a suitable position for assessing tRNA orthogonality. T216 was mutated to every canonical amino acid and expressed in CFE. Each bar represents the average of n = 3 data points and the error bar represents one standard deviation.

**Figure S3:**
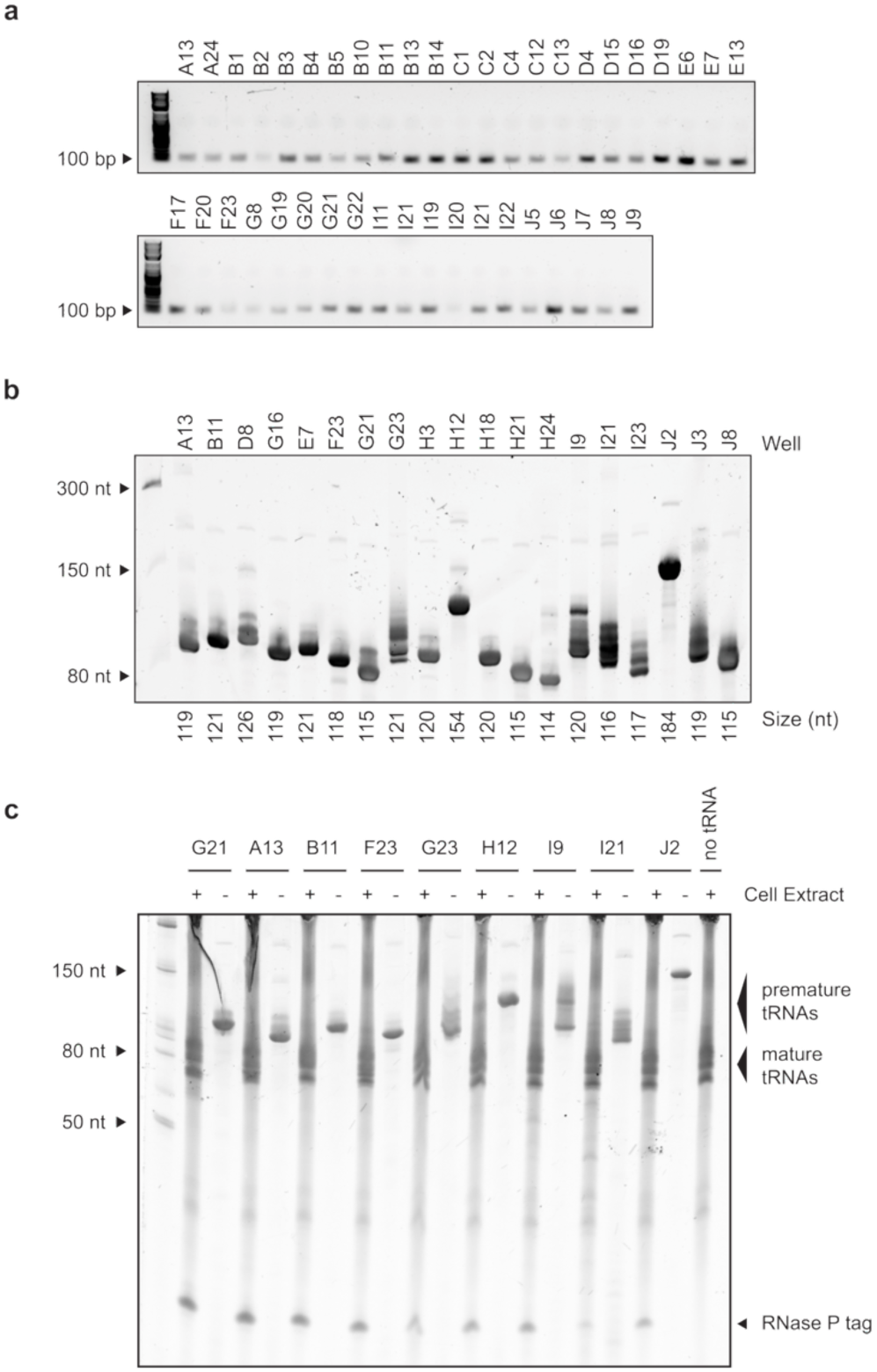
Analysis of products from tRNA expression and maturation support conclusion of orthogonality. (**a**) DNA templates were correctly amplified from commercial templates as measured by agarose gel electrophoresis. (**b**) *in vitro* transcription products are correctly synthesized from PCR-amplified templates as measured by denaturing urea PAGE. (**c**) *In vitro* transcription products are processed by RNase P to remove the tag as measured by denaturing urea PAGE. Depletion of the premature tRNA band along with emergence of RNase P tag suggests correct processing of tRNAs. Gels were collected and analyzed once.

**Figure S4:**
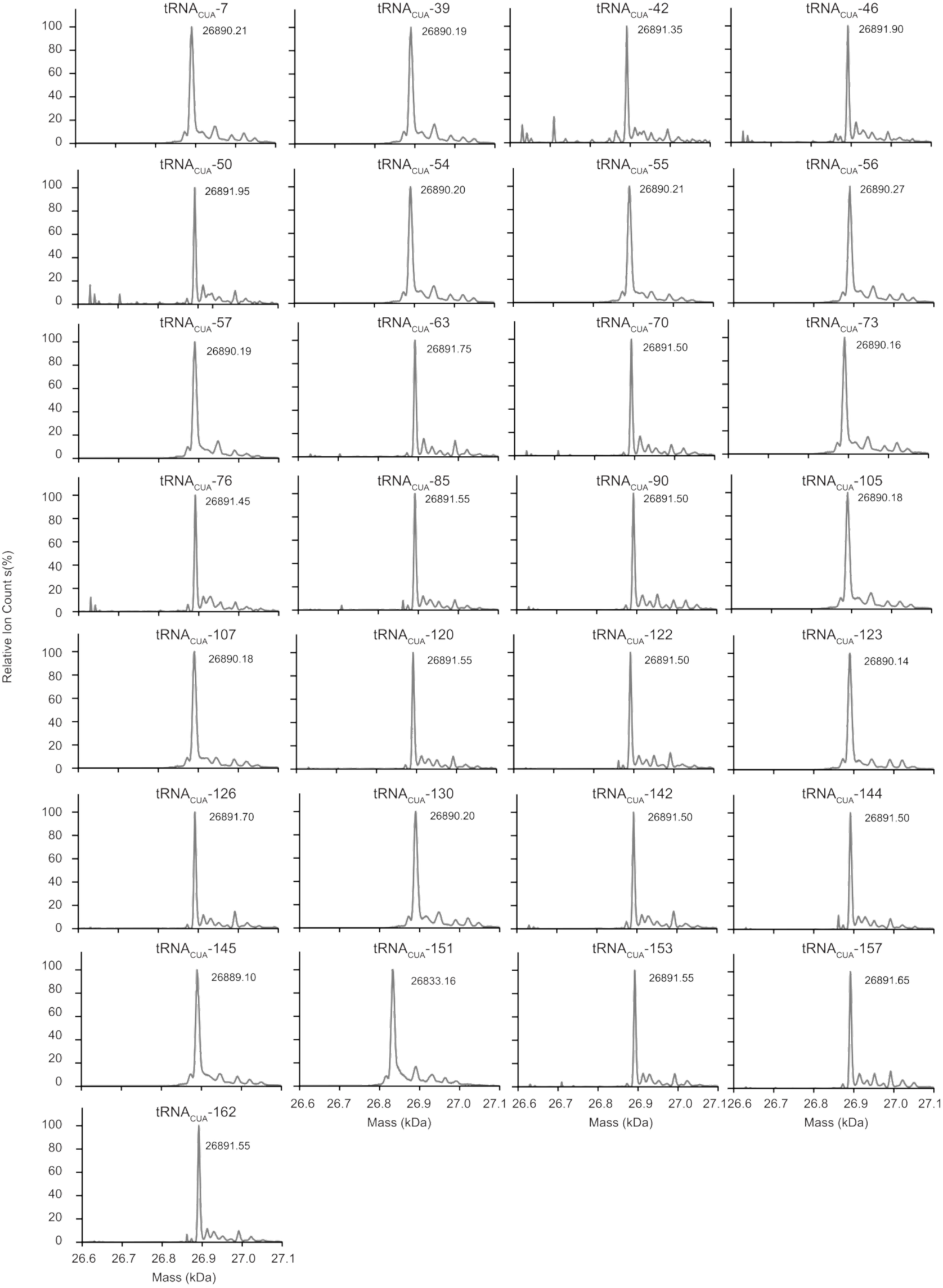
ESI-MS of purified 216UAG-sfGFP purified proteins from CFE reactions with non-orthogonal tRNAs. 216UAG-sfGFP expressed when using a non-orthogonal suppressor tRNA were purified and analyzed by ESI-MS. These tRNAs span 216X-sfGFP synthesis yields from 10-100% of WT sfGFP. Most (28/29) tRNAs enable the incorporation of Gln or Lys in 216UAG-sfGFP. tRNA_CUA_-151 enables the incorporation of Ala in 216UAG-sfGFP. Spectra were collected twice and a representative spectra is shown.

**Figure S5:**
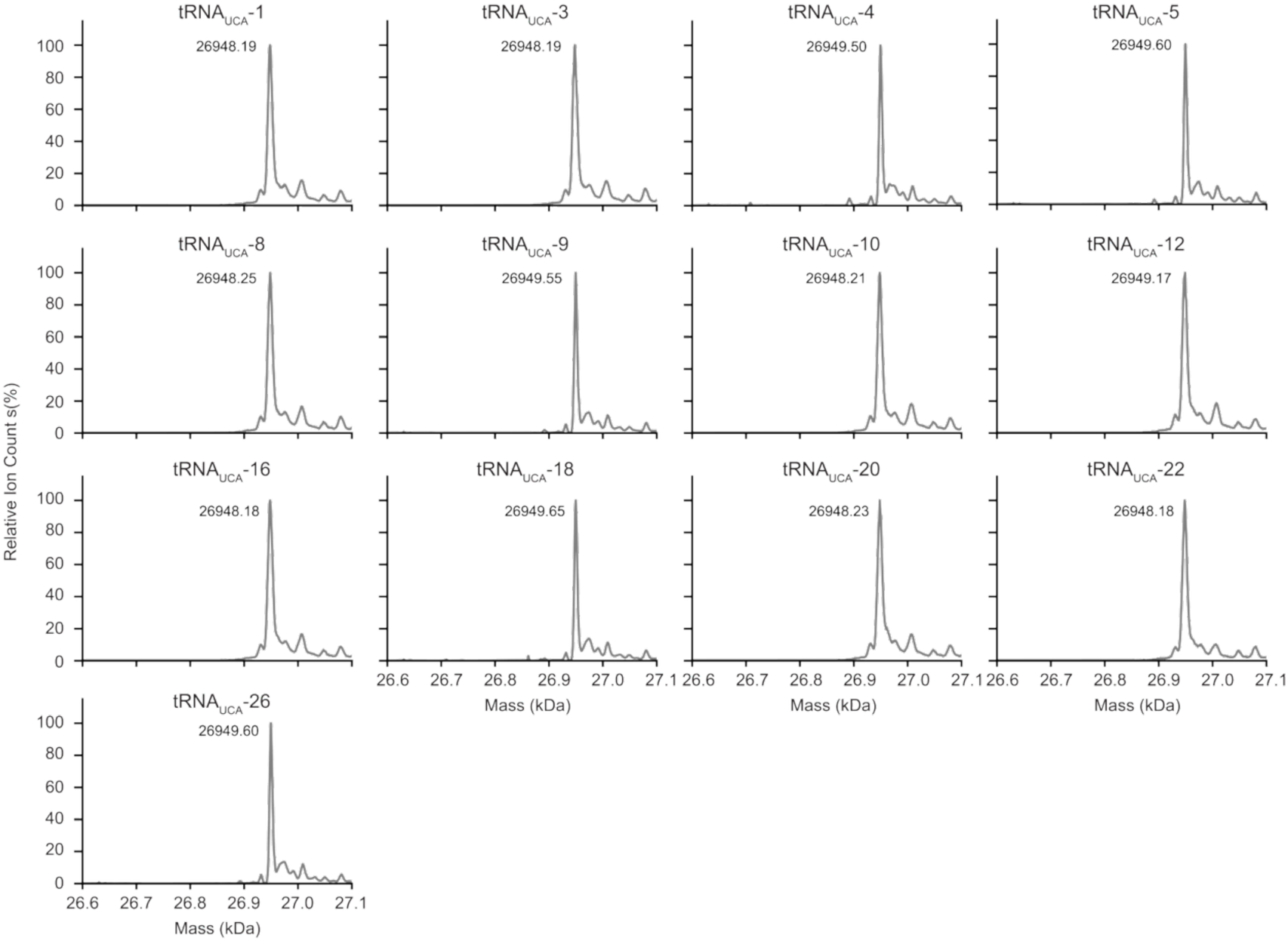
ESI-MS of 216UGA-sfGFP from CFE reactions with non-orthogonal tRNAs show incorporation of Trp in all cases. 216UGA-sfGFP was expressed in CFE reactions supplemented with a non-orthogonal tRNA_UCA_. Spectra were collected twice and a representative spectra is shown.

**Figure S6:**
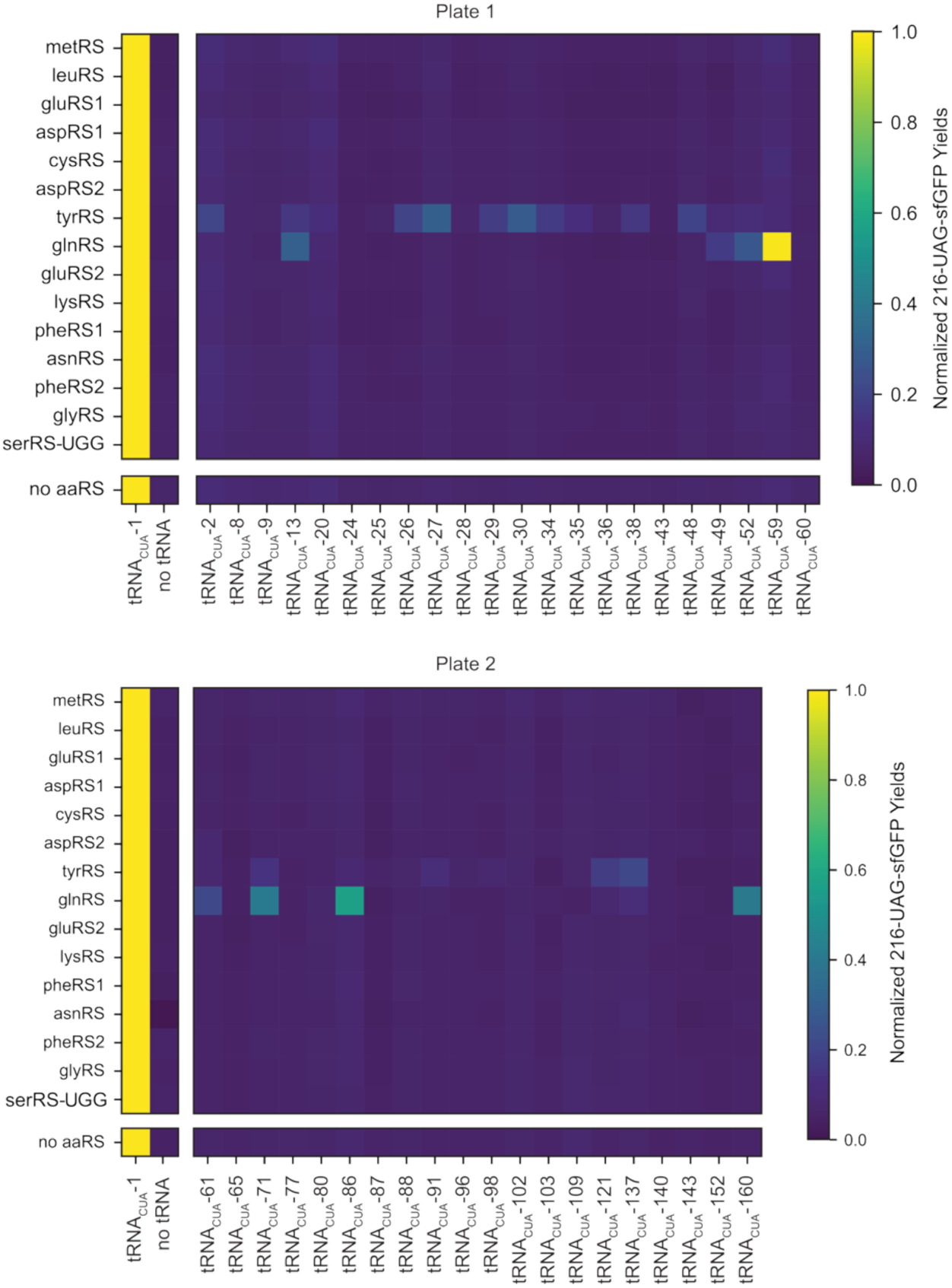
Full screening results from CFE reactions to identify functional aaRS:tRNA pairs using metagenomically identified aaRSs and orthogonal tRNAs_CUA._ 216UAG-sfGFP is normalized to 216UAG-sfGFP signal from tRNA_CUA_-1 in each row. tRNA_CUA_-1 is a non-orthogonal tRNA used as a positive control. Each heatmap data point is n = 1 and the experiment was repeated twice with similar results.

**Figure S7:**
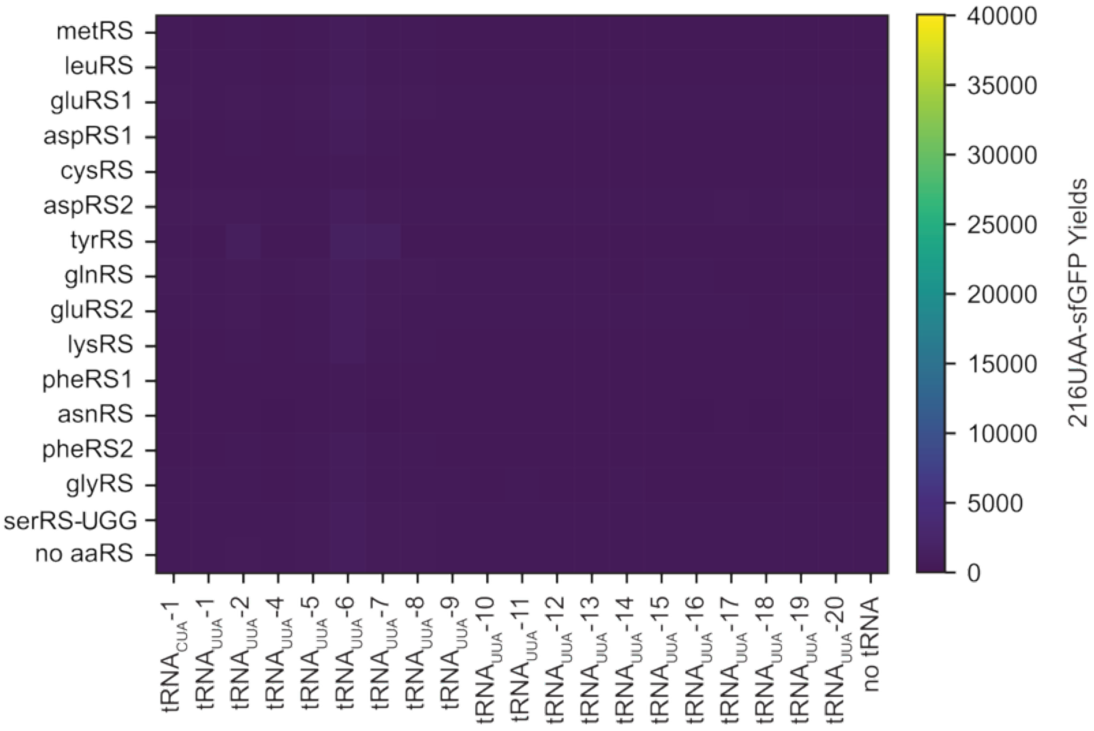
Screening results from CFE reactions to identify functional aaRS:tRNA pairs using metagenomically identified aaRSs and orthogonal tRNAs_UUA_. No positive tRNA control was included because an active tRNAs_UUA_ was not identified in Figure 1b. tRNA_CUA_-1 is used as an additional negative control in this experiment. Unnormalized data is shown to highlight that no tRNAs resulted in any 216UAA-sfGFP signal above background.

**Figure S8:**
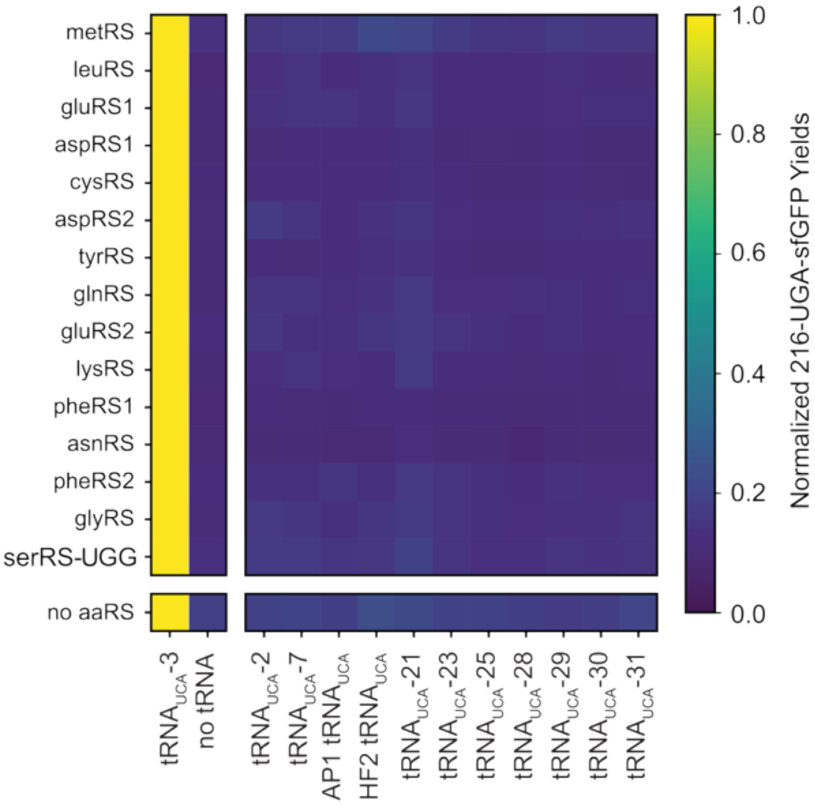
Screening results from CFE reactions to identify functional aaRS:tRNA pairs using metagenomically identified aaRSs and orthogonal tRNAs_UCA_. tRNA_UCA_-3 is used as a positive control in this experiment.

**Figure S9:**
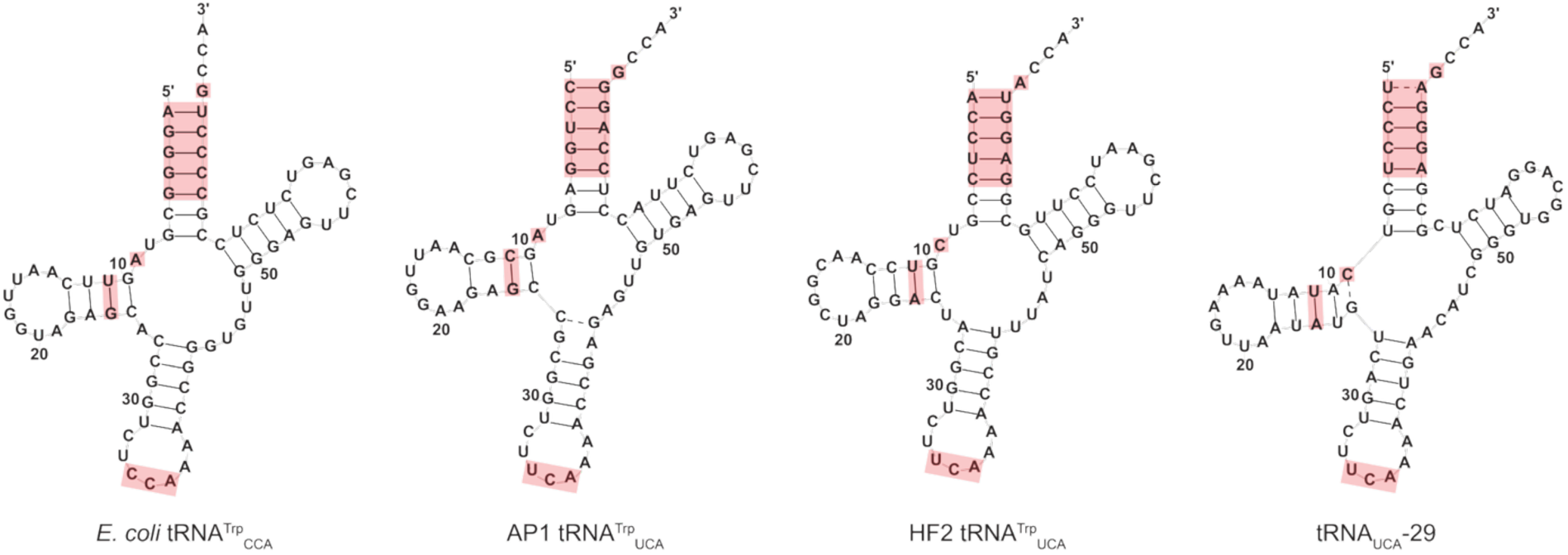
Comparison of tRNA secondary structures as predicted by R2DT.^1^ Identity elements for *E. coli* tRNA^Trp^_CCA_ are highlighted in red, and the corresponding nucleotides for the AP1, HF2, and tRNA_UCA_-29 tRNAs_UCA_ are highlighted in red.

**Figure S10:**
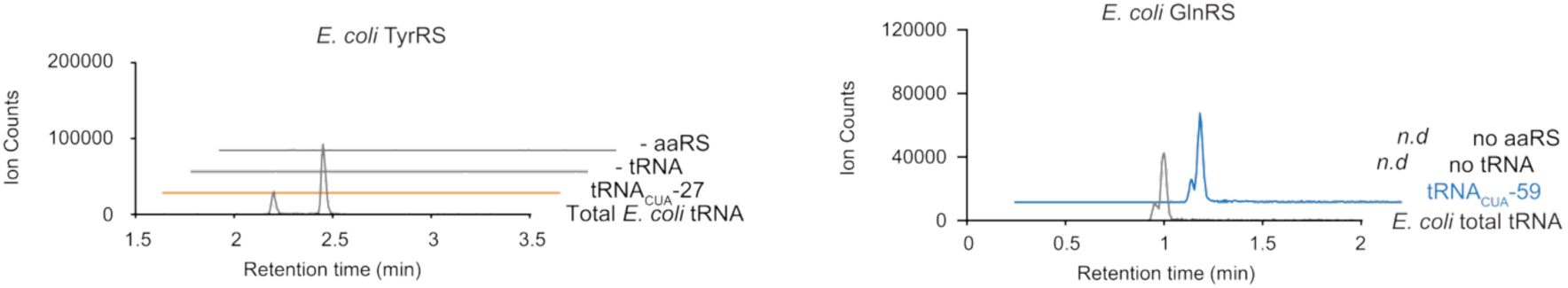
Aminoacylation assays using endogenous *E. coli* aaRSs to identify Gln-A and Tyr-A products and to test activity against tRNA_CUA_-27 and tRNA_CUA_-59. Tyr-A elutes at retention times of 2.20 and 2.45 min and and Gln-A elutes at retention times of 0.96 and 0.99 min. tRNA_CUA_-27 is orthogonal to *E. coli* TyrRS, but tRNA_CUA_-59 is aminoacylated by *E. coli* GlnRS in *in vitro* conditions. A representative spectra from n = 2 experiments is shown.

**Figure S11:**
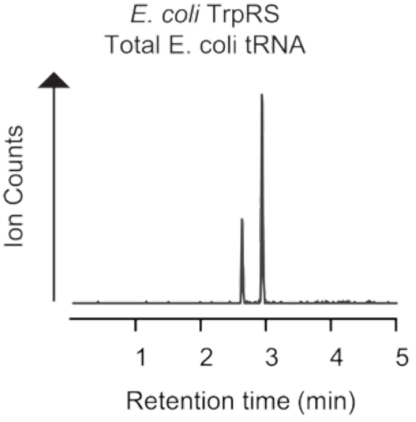
Positive control aminoacylation reactions using *E. coli* TrpRS and total *E. coli* tRNA show elution of Trp-A at retention times of 2.63 min and 2.93 min as measured by LC-MS. Representative spectra from n = 3 experiments is shown.

**Figure S12:**
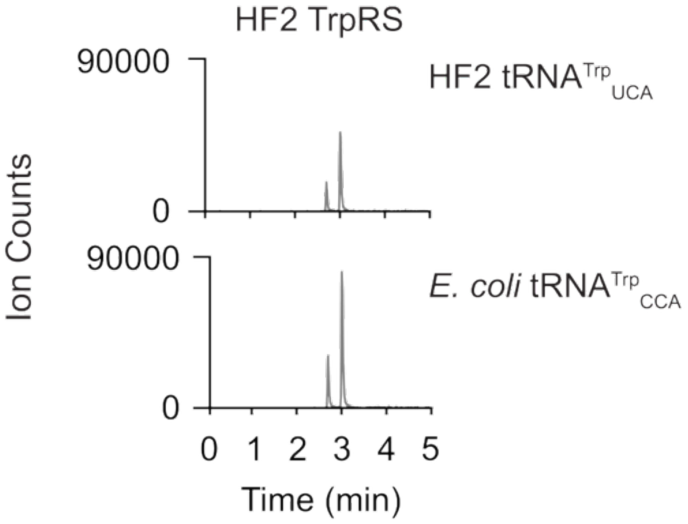
HF2 TrpRS nonspecifically aminoacylates *E. coli* tRNA^Tro^_CCA_ with Trp as measured by LC-MS of aminoacylation assays. Spectra was collected once.

**Figure S13:**
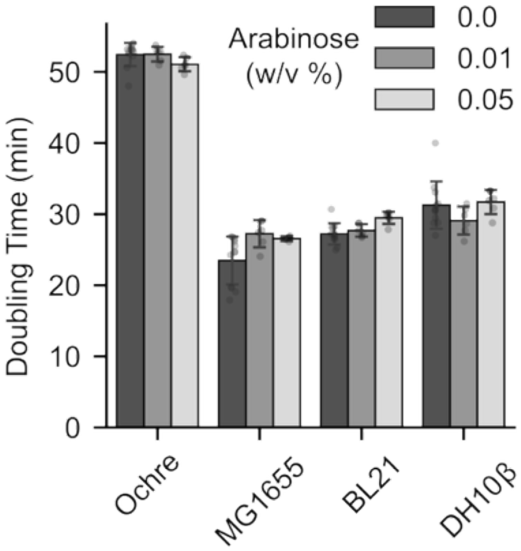
Bar chart of doubling time during induction of AP1 TrpRS: AP1 tRNA_UCA_ in common *E. coli* laboratory strains shows no observable growth defect during exponential growth. AP1 OTS is induced by addition of 0.05% w/v arabinose. Bars represent n = 12 replicates and error bars represent one standard deviation.

**Figure S14:**
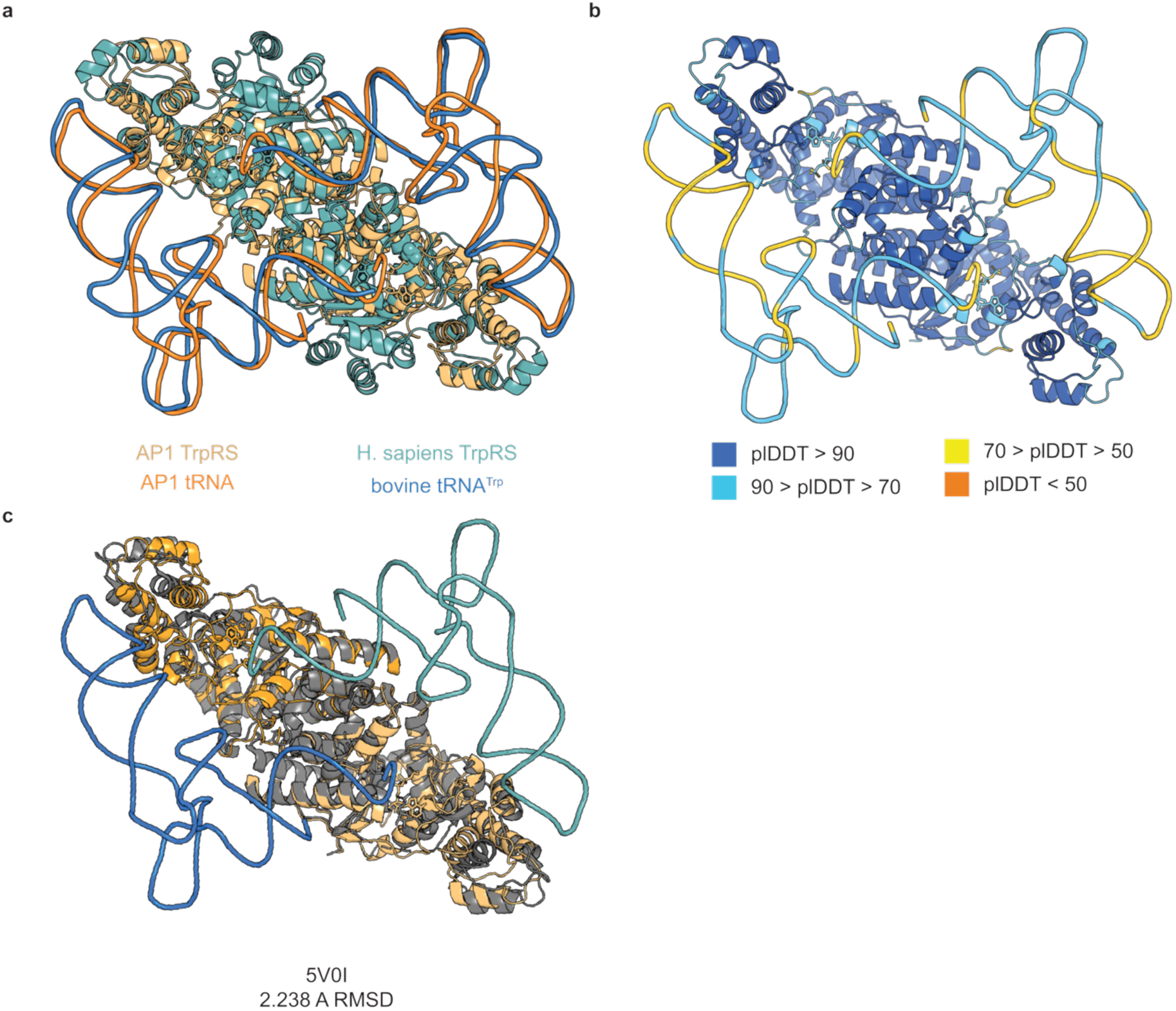
Analysis of the AlphaFold3 structural model of the AP1 TrpRS:tRNA complex. (**a**) The AlphaFold3 prediction of the AP1 TrpRS:tRNA complex overlays well with the crystal structure of the human TrpRS complexed with a bovine tRNA^Trp^. The AP1 complex is shown in shades of orange, and the human TrpRS:bovine tRNA complex is shown in shades of blue. (**b**) AlphaFold3 predicts the structure of the AP1 TrpRS:tRNA complex with high confidence metrics. pLDDT for each residue and nucleotide in the TrpRS and tRNA are shown, respectively. The TrpRS is confidently predicted with an average pLDDT = 89.8, and the tRNA is predicted with moderate confidence pLDDT = 72.0. (**c**) The AP1 TrpRS structure aligns with an RMSD of 2.24 Å against a crystal structure of the *E. coli* TrpRS (PDB ID: 5v0i). The *E. coli* TrpRS is shown in gray, the AP1 TrpRS is shown in shades of orange, and the AP1 tRNA is shown in shades of blue.

### Supplementary Tables

**Table S1:**
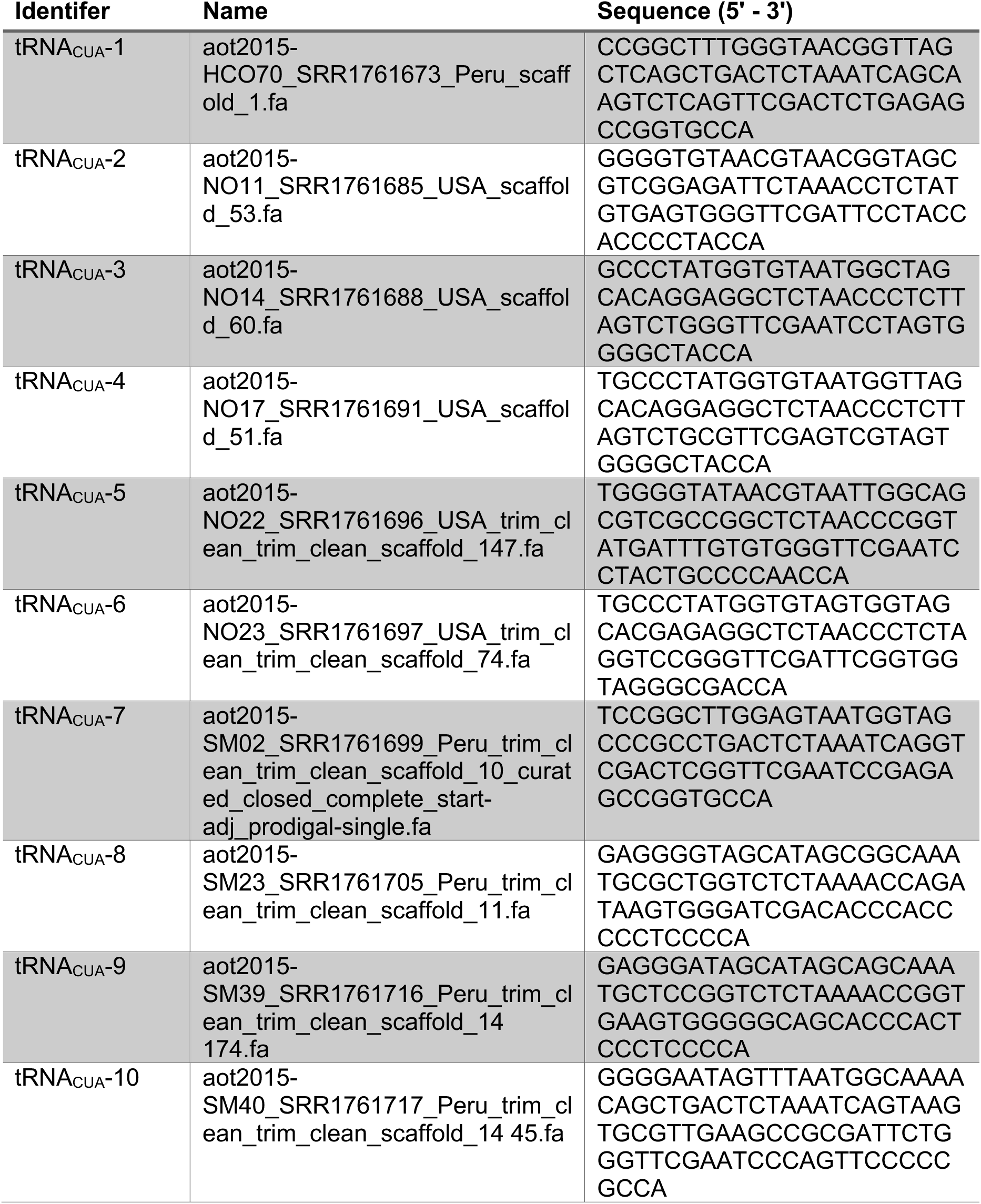
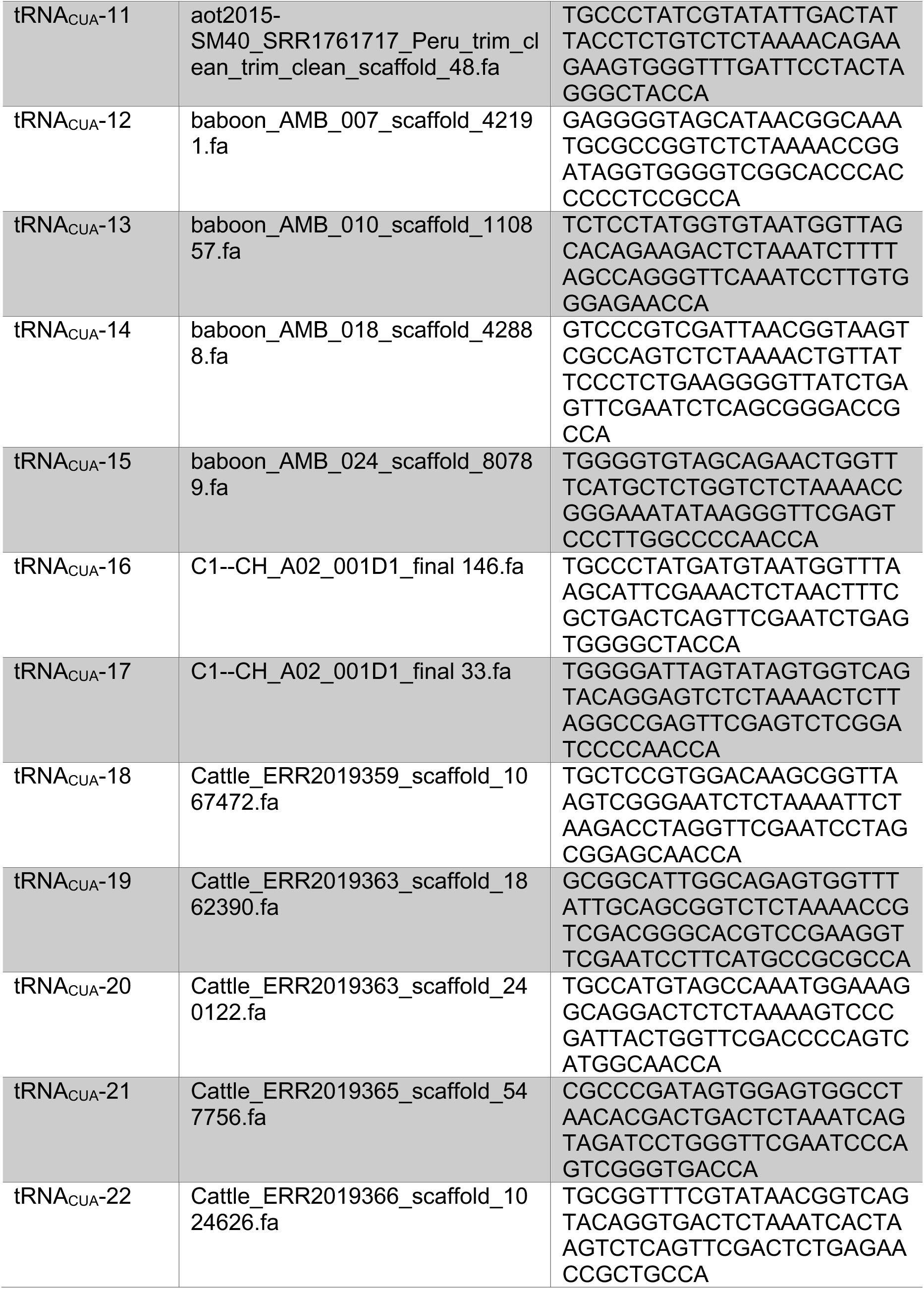
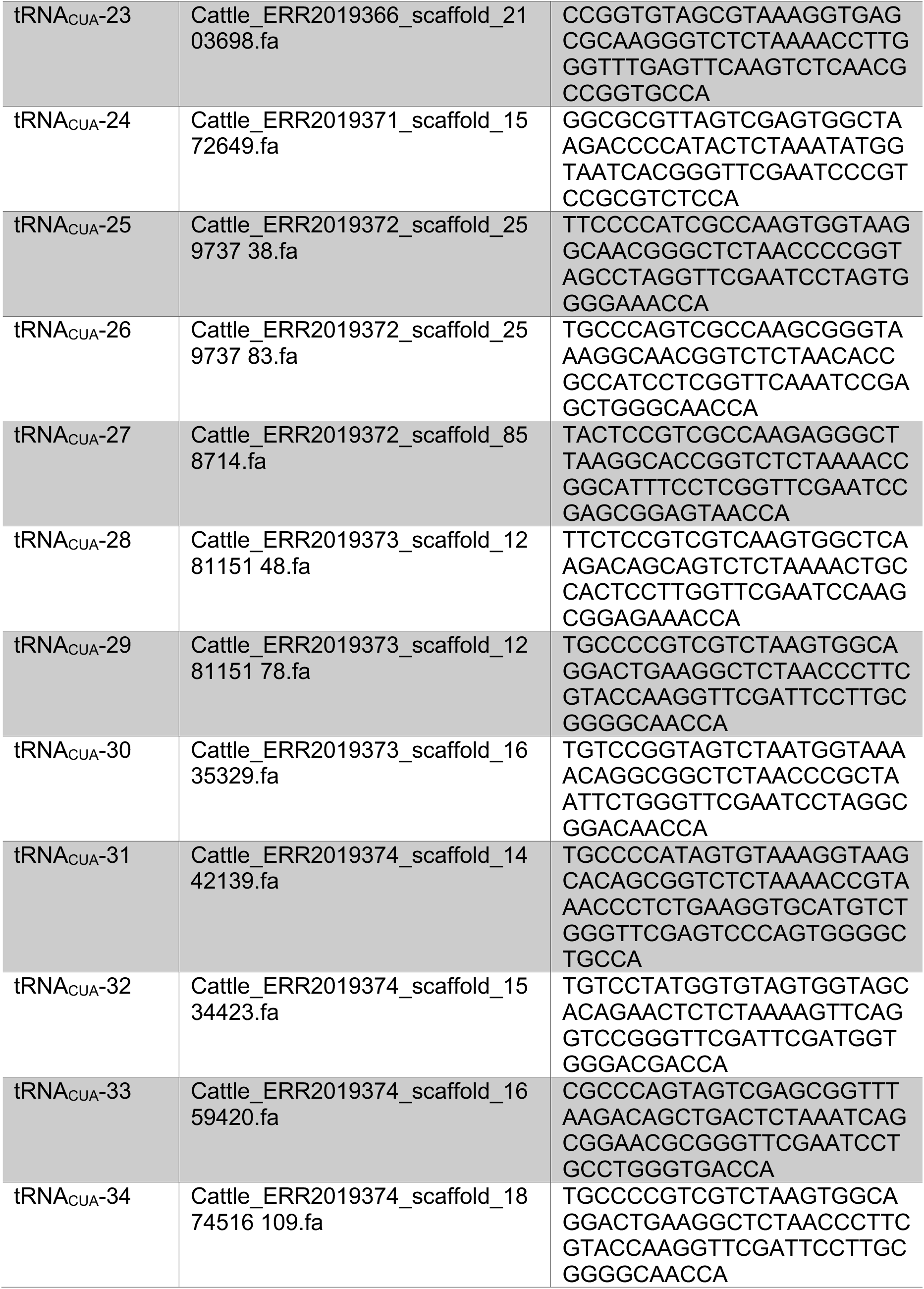
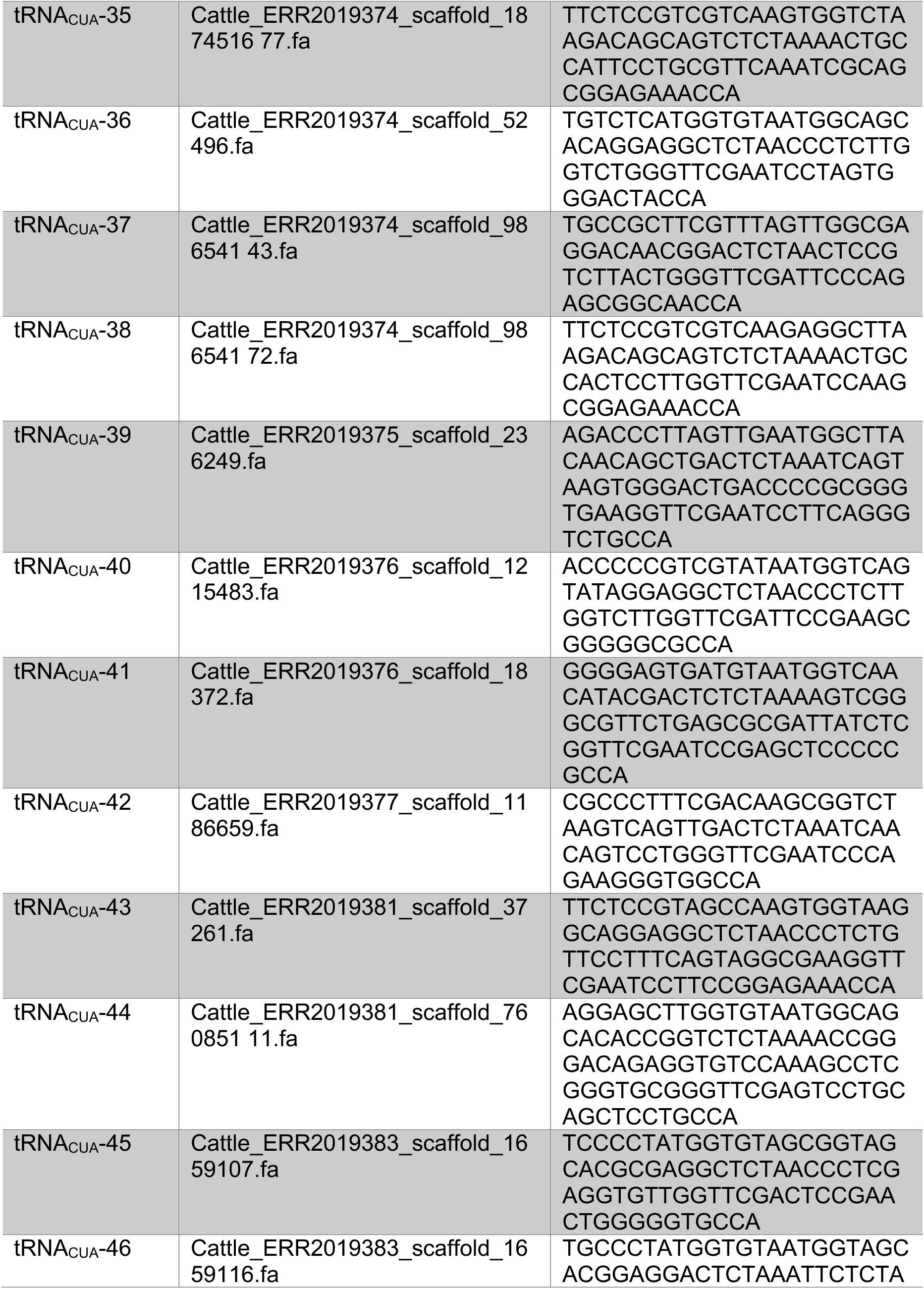
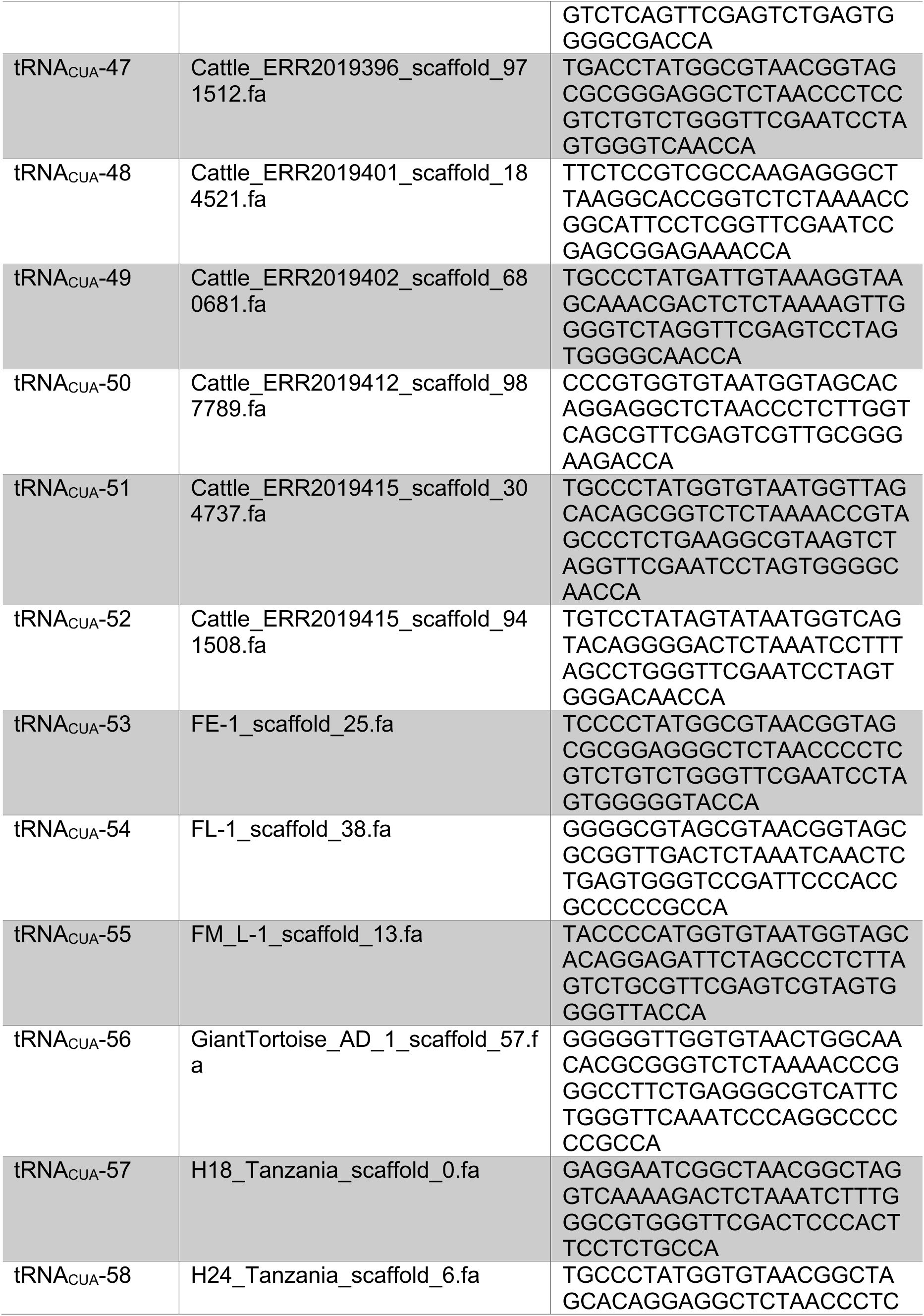
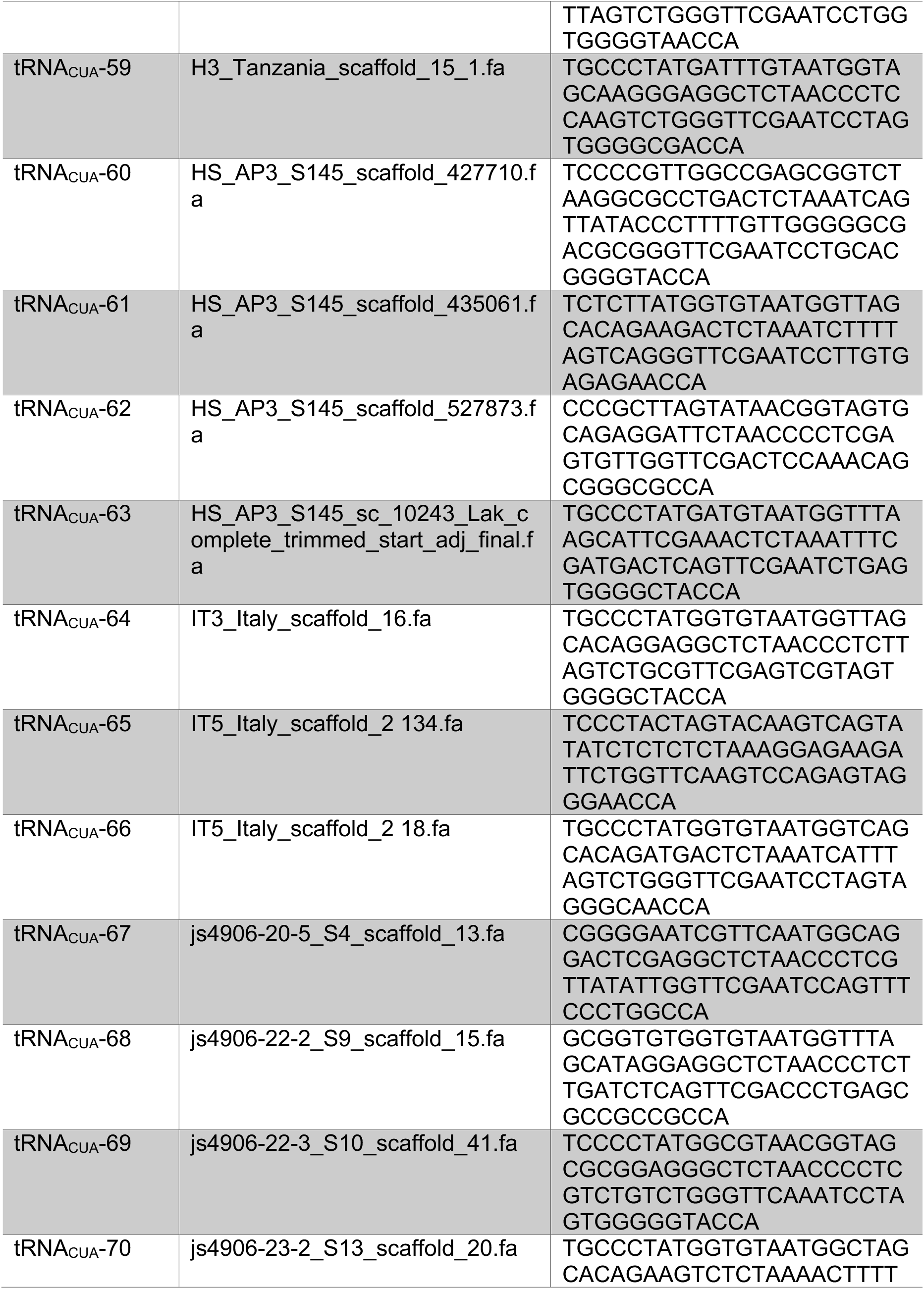
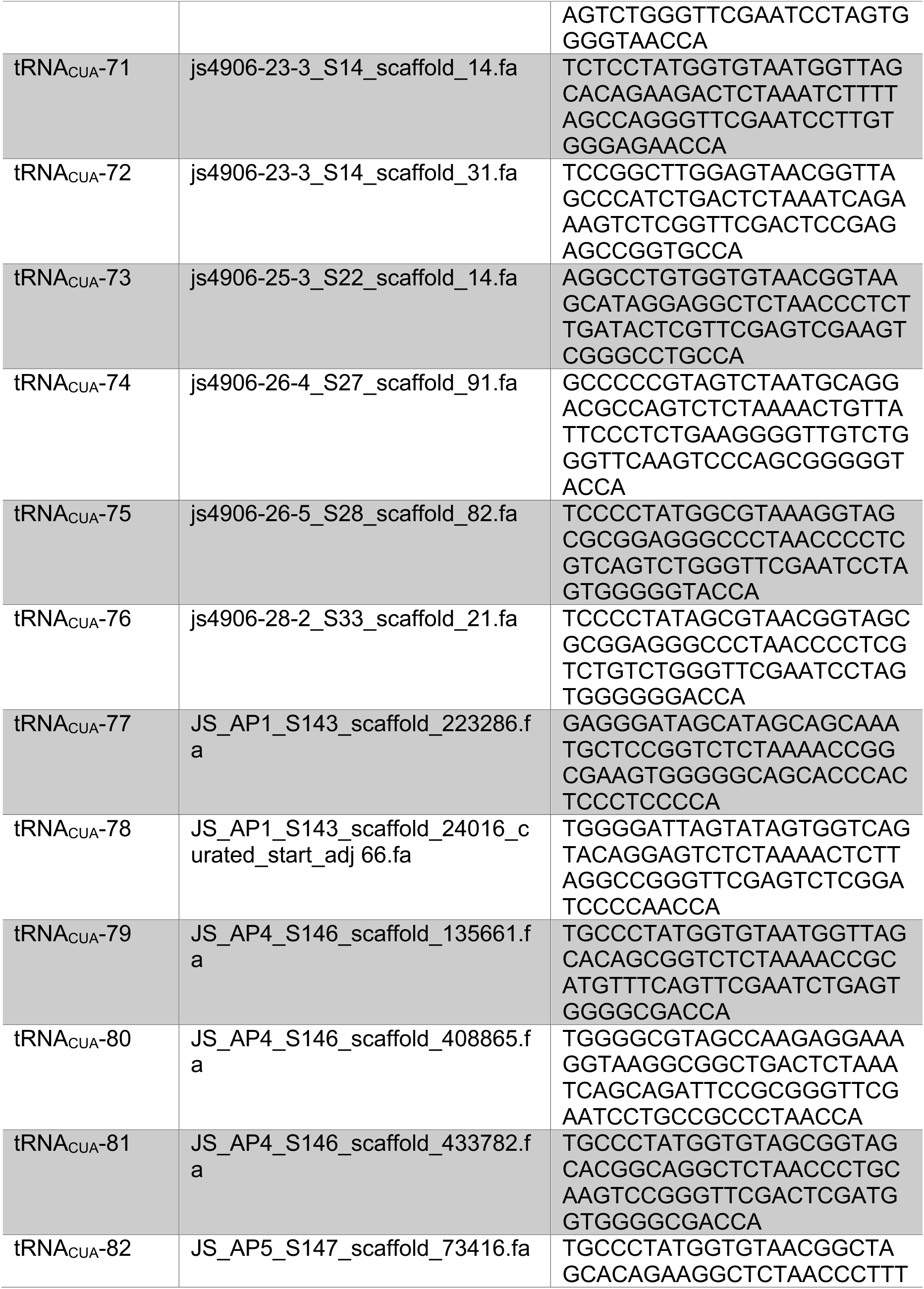
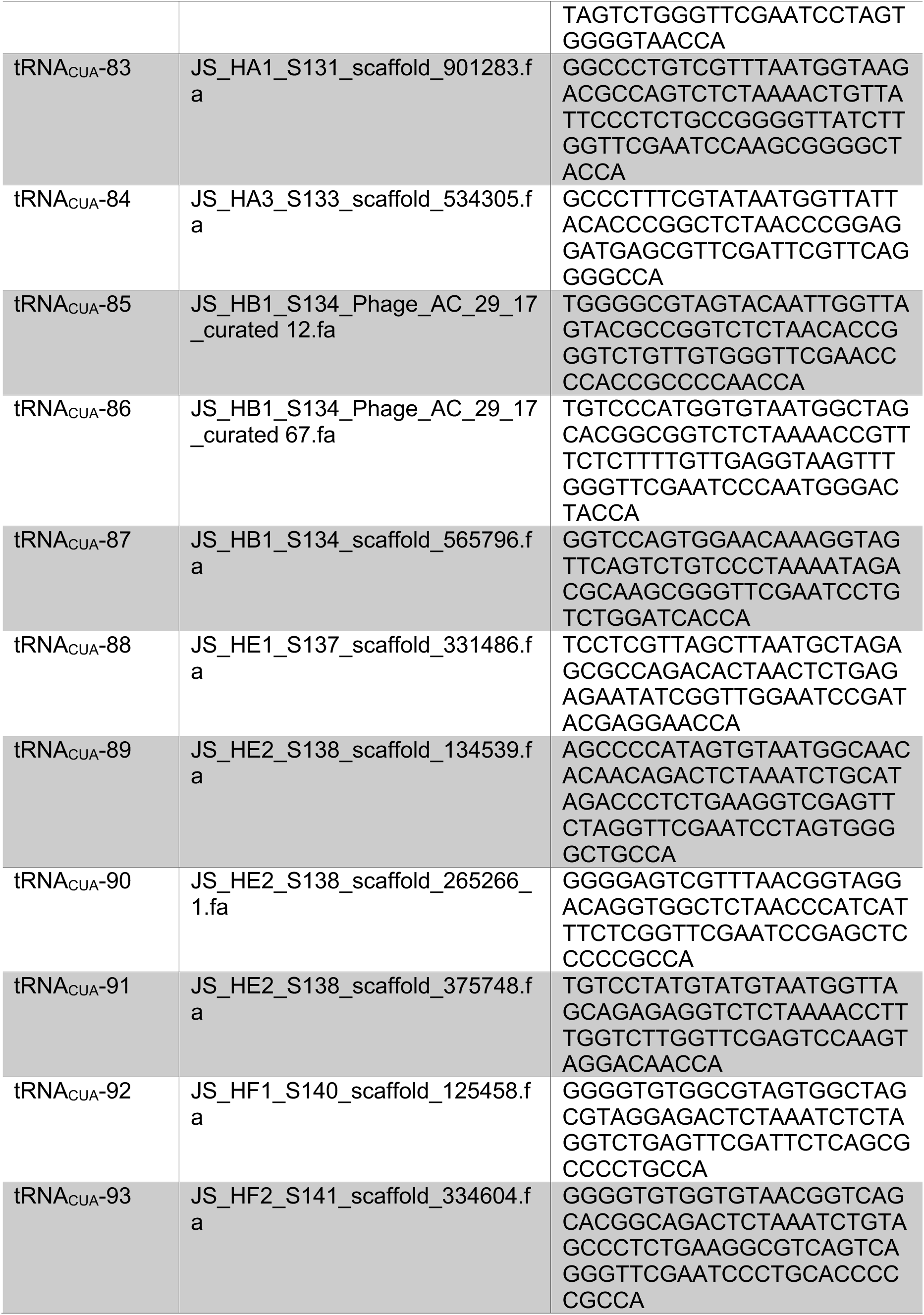
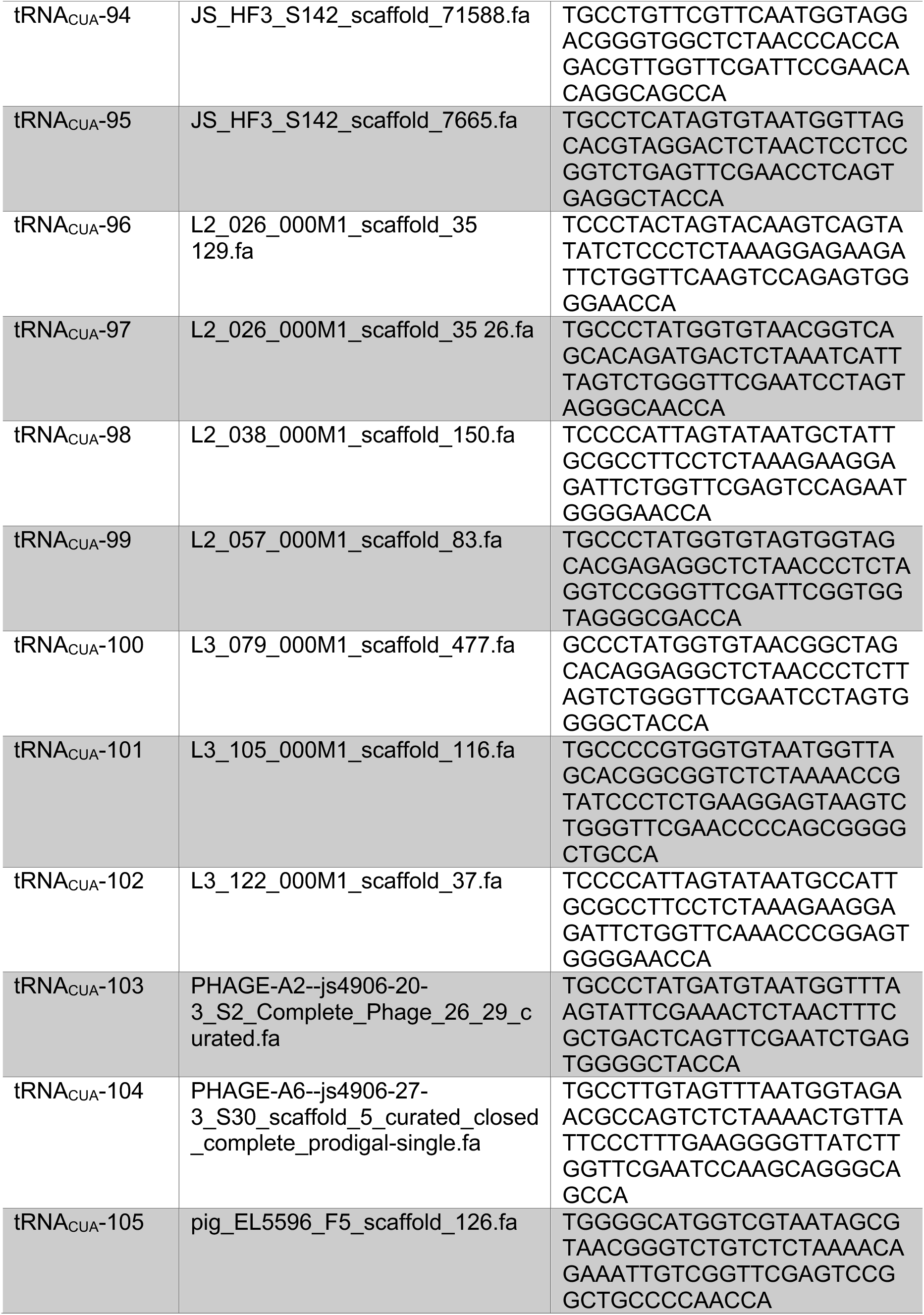
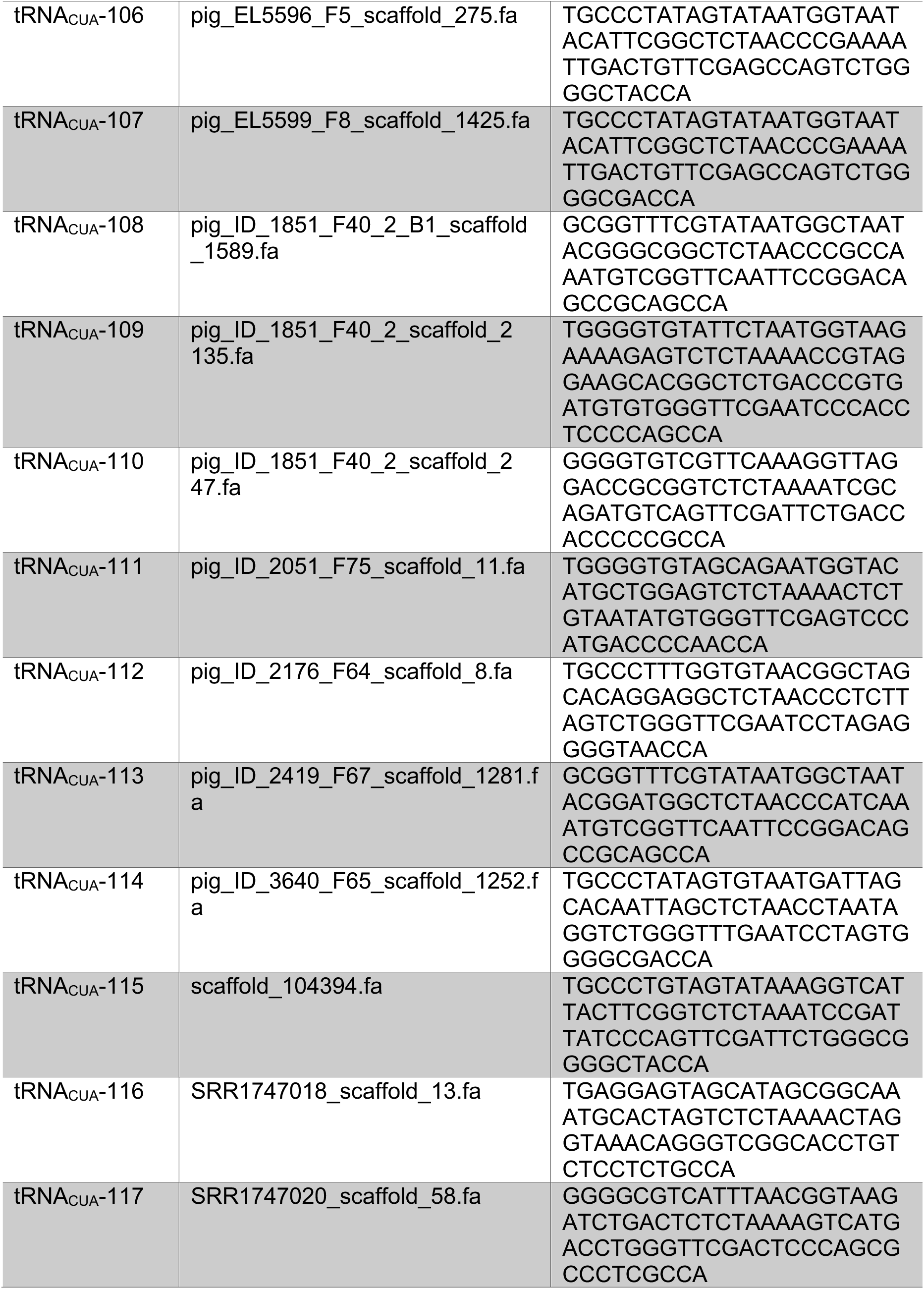
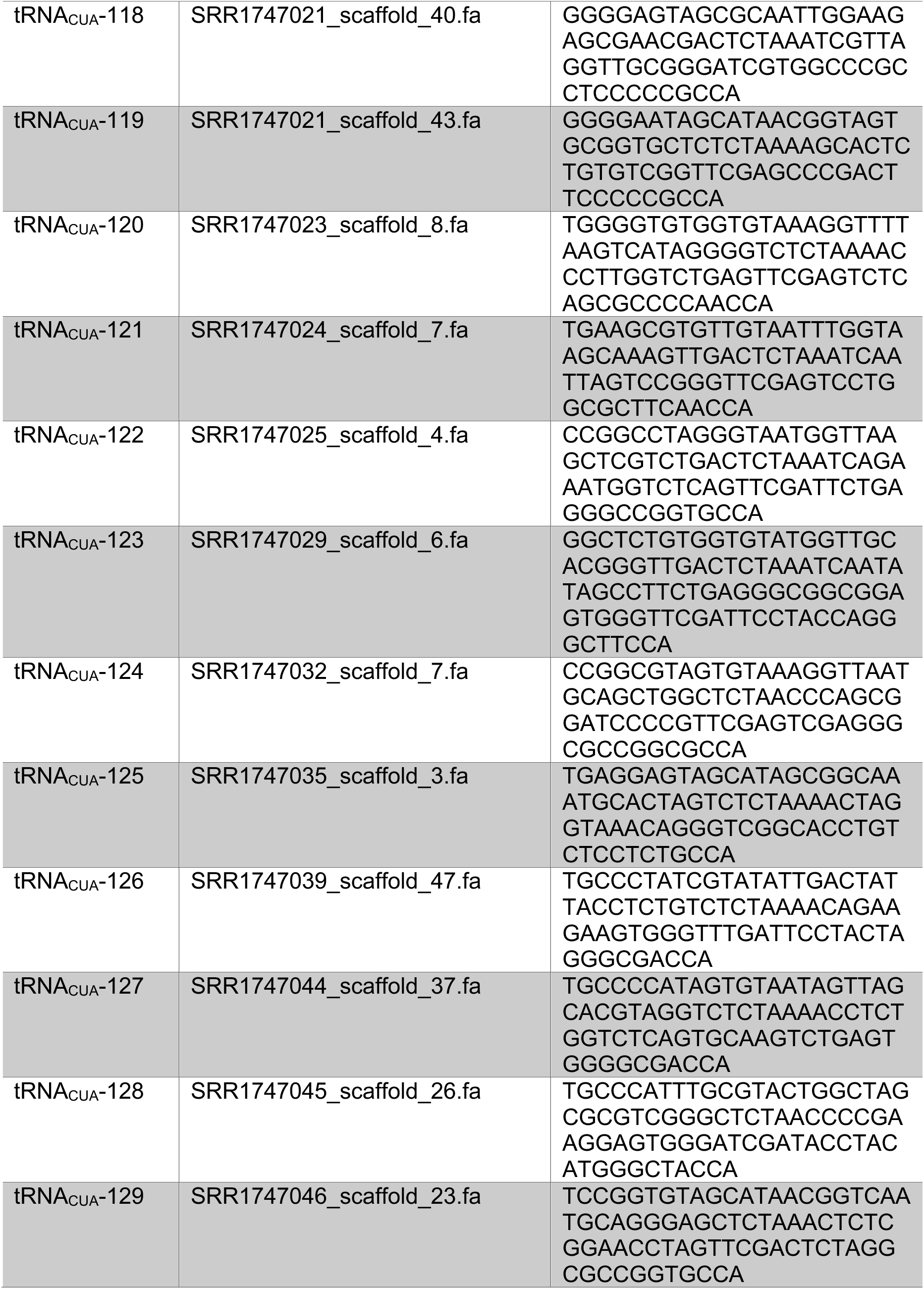
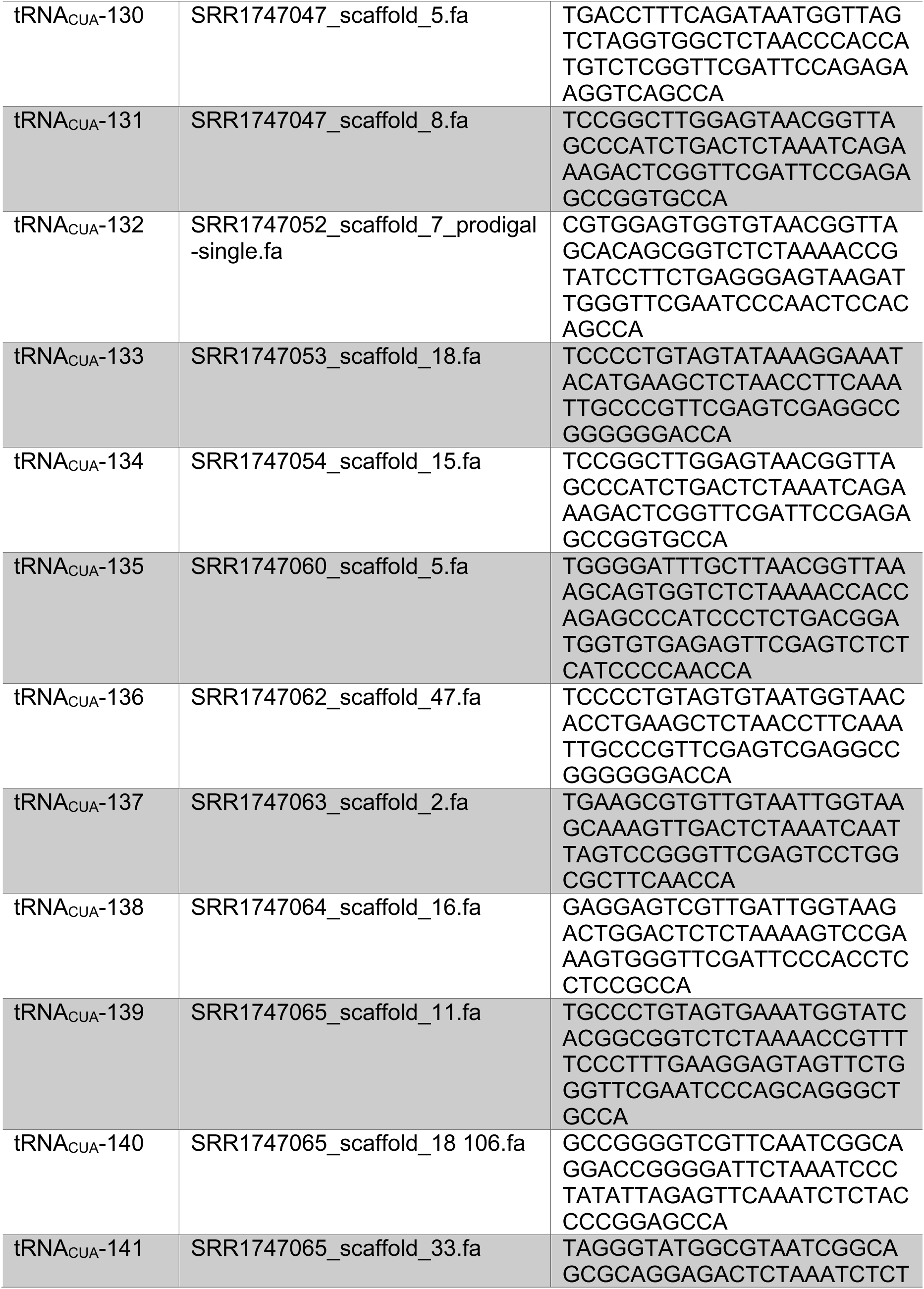
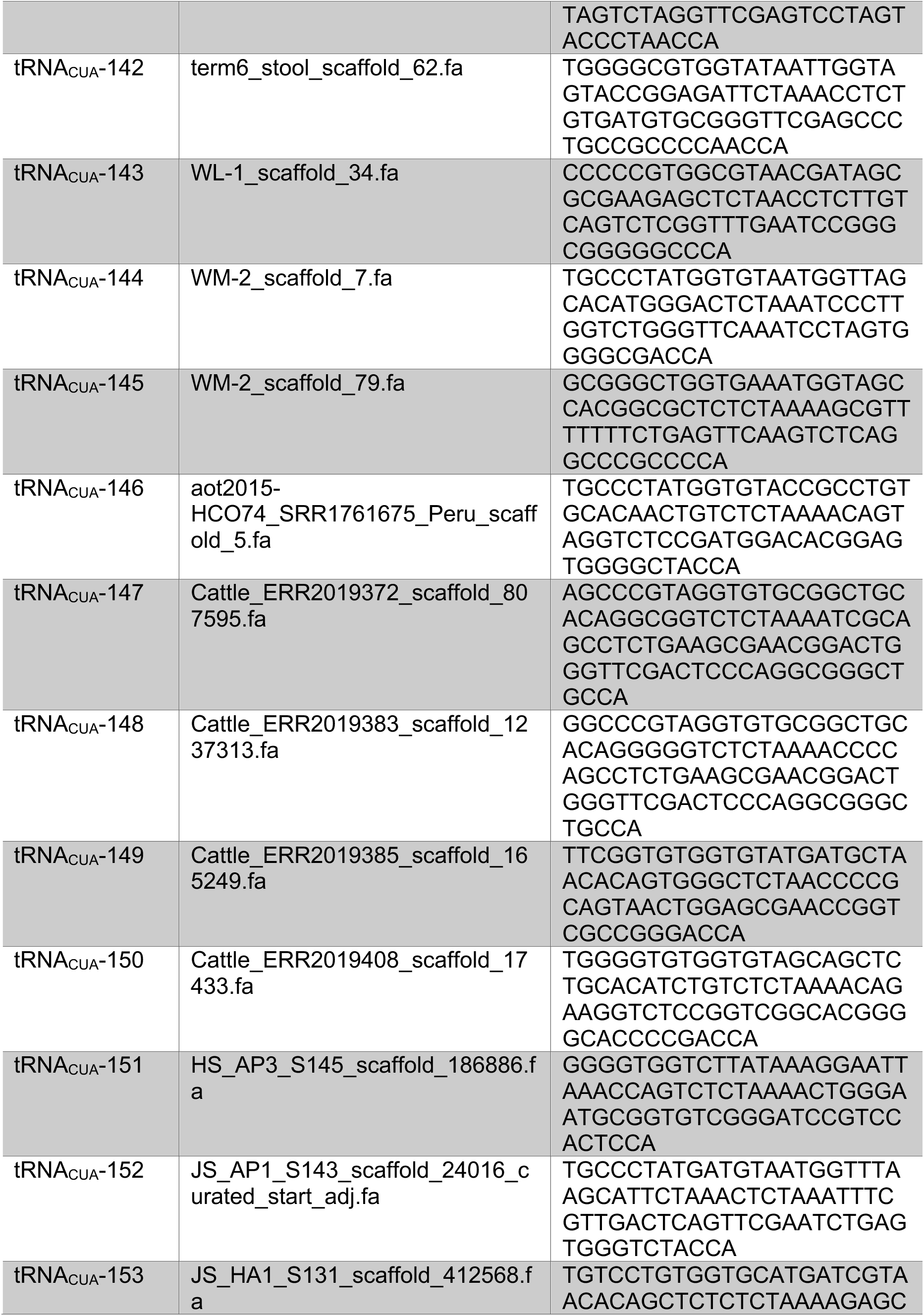
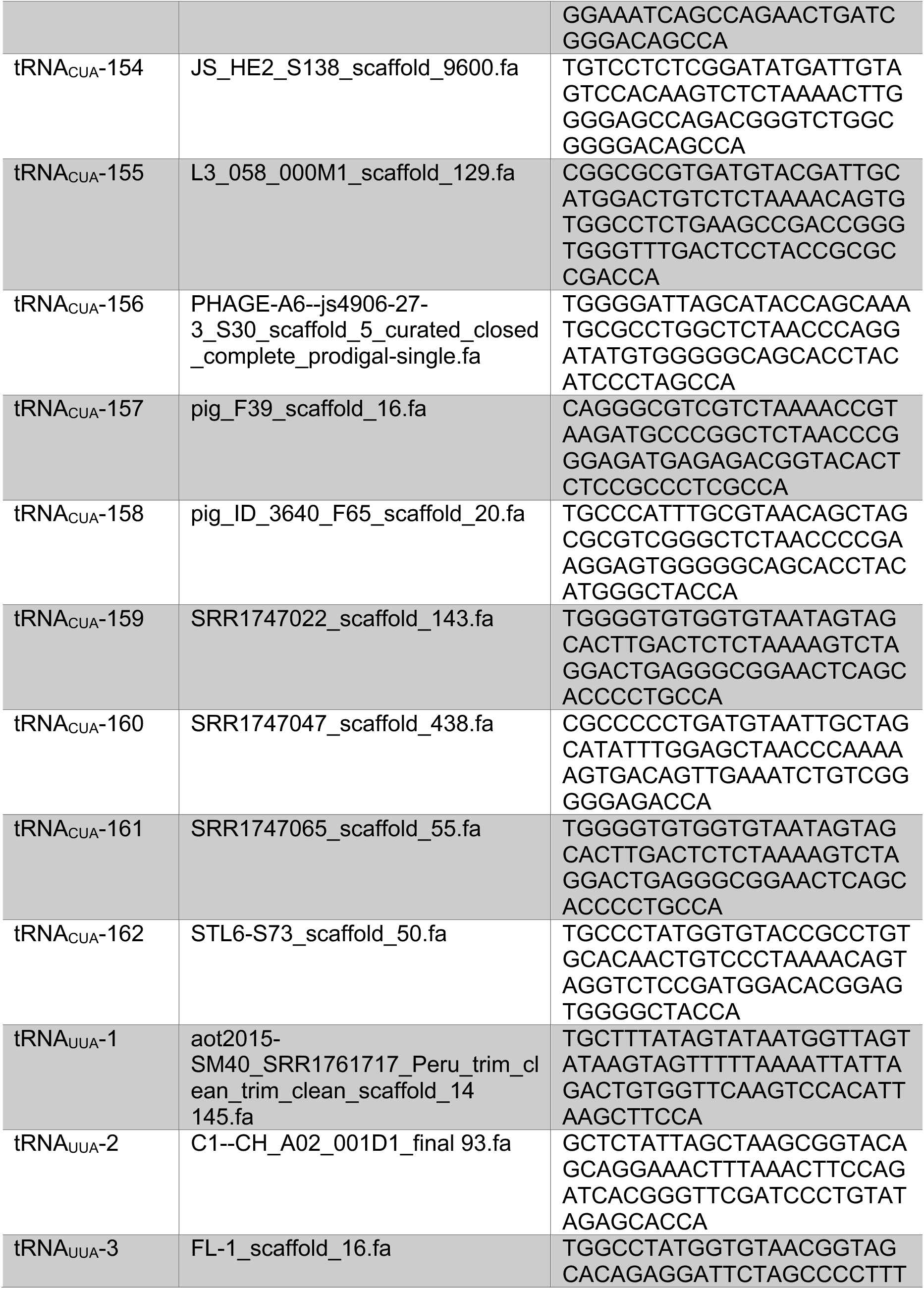
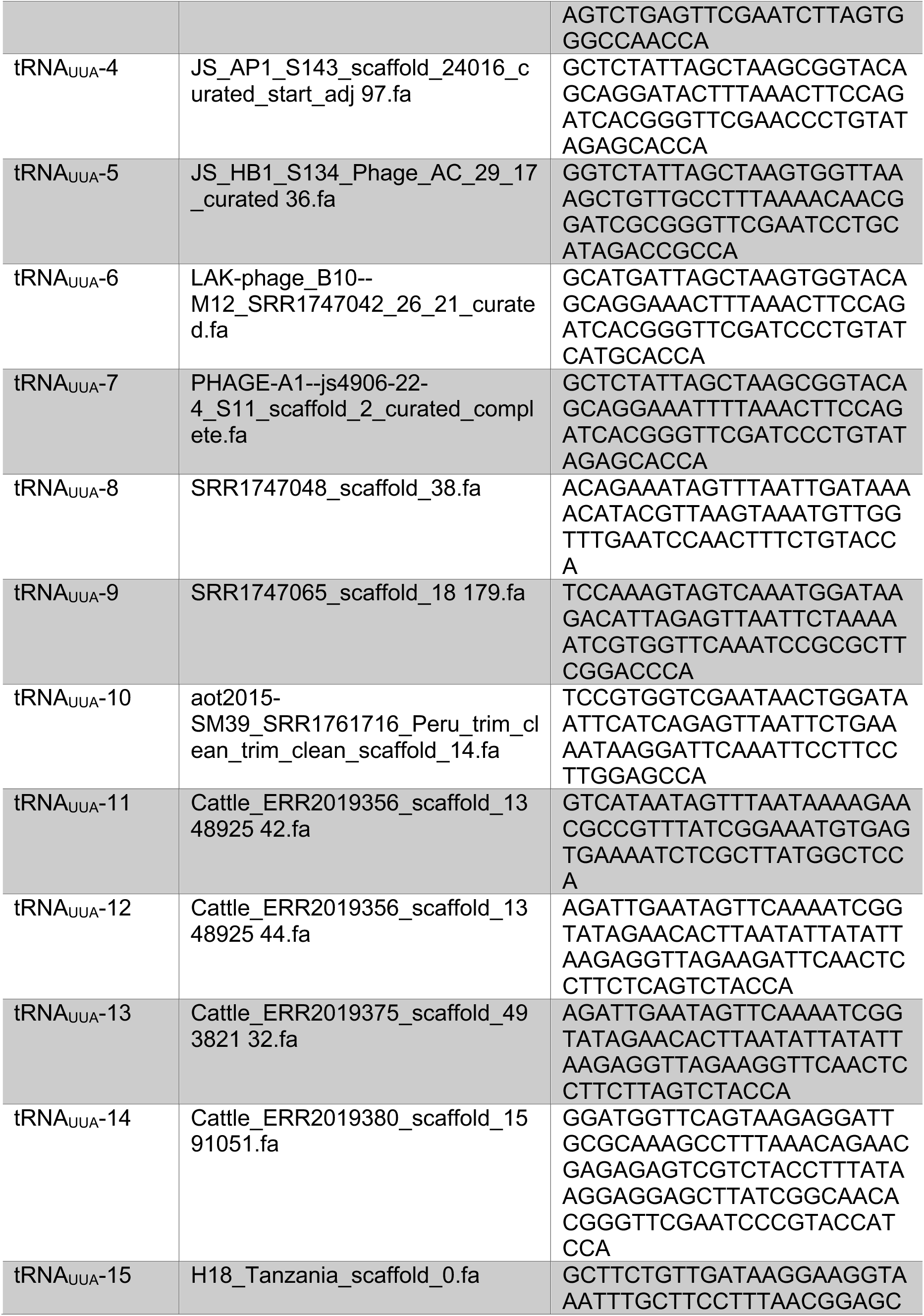
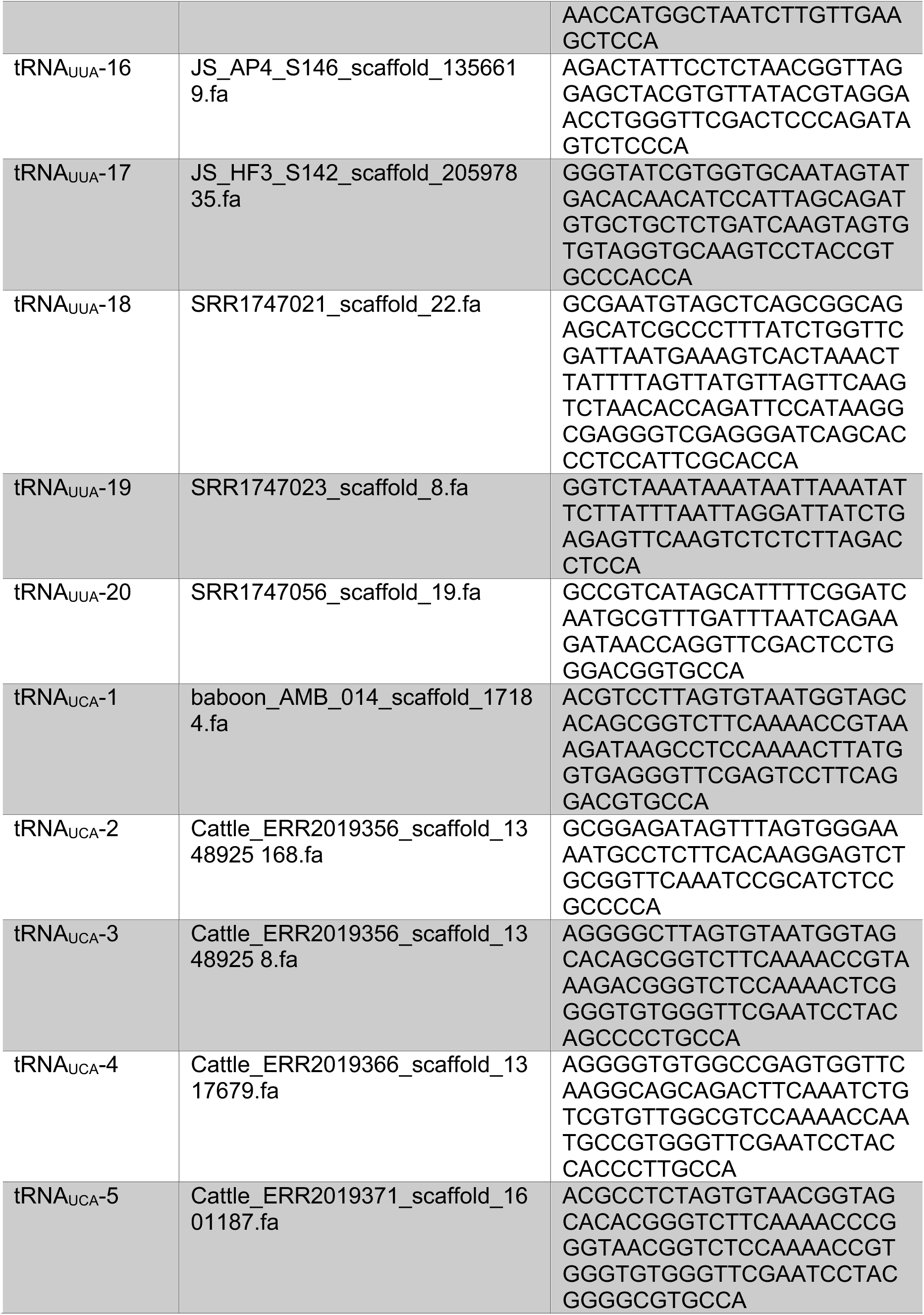
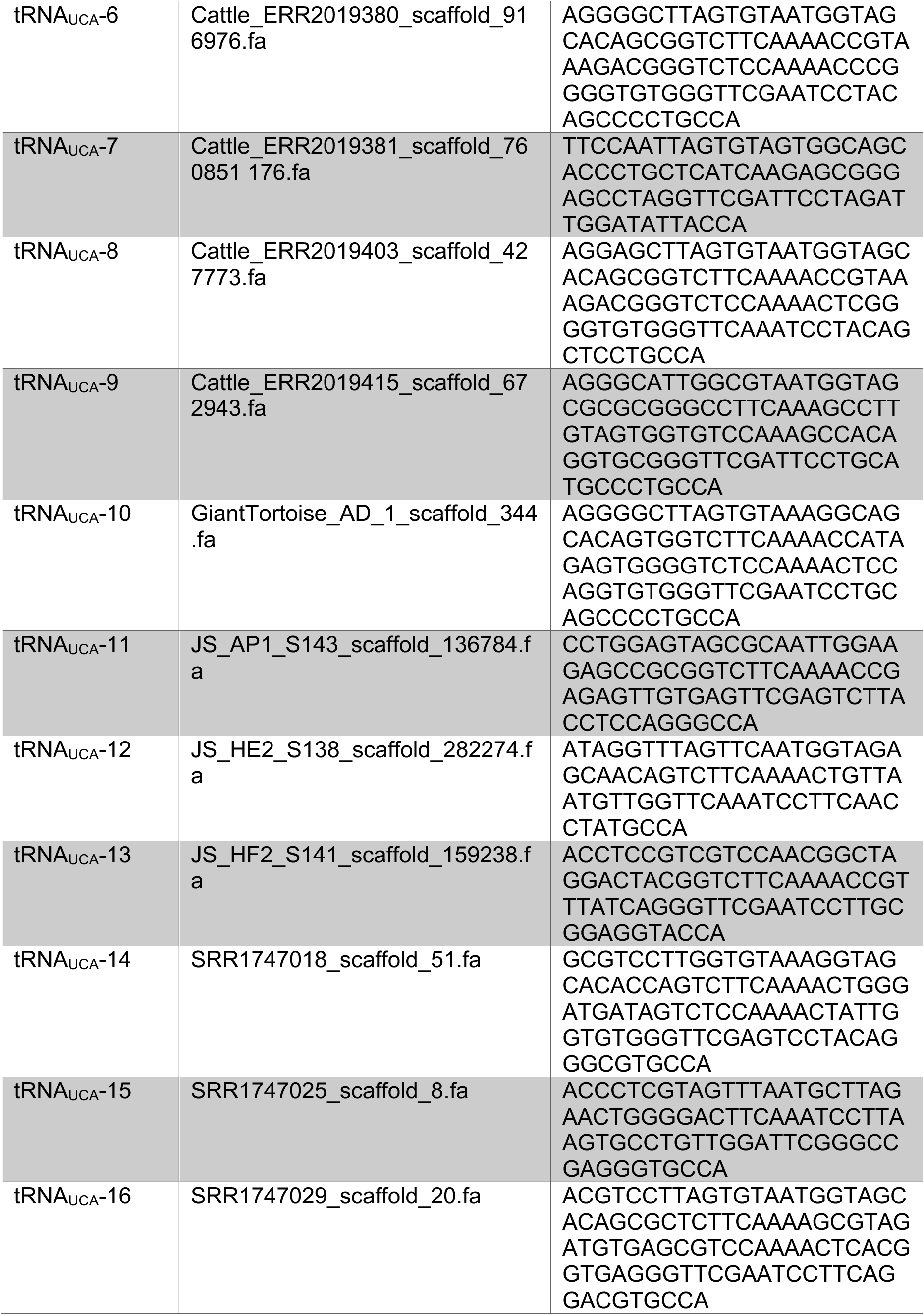
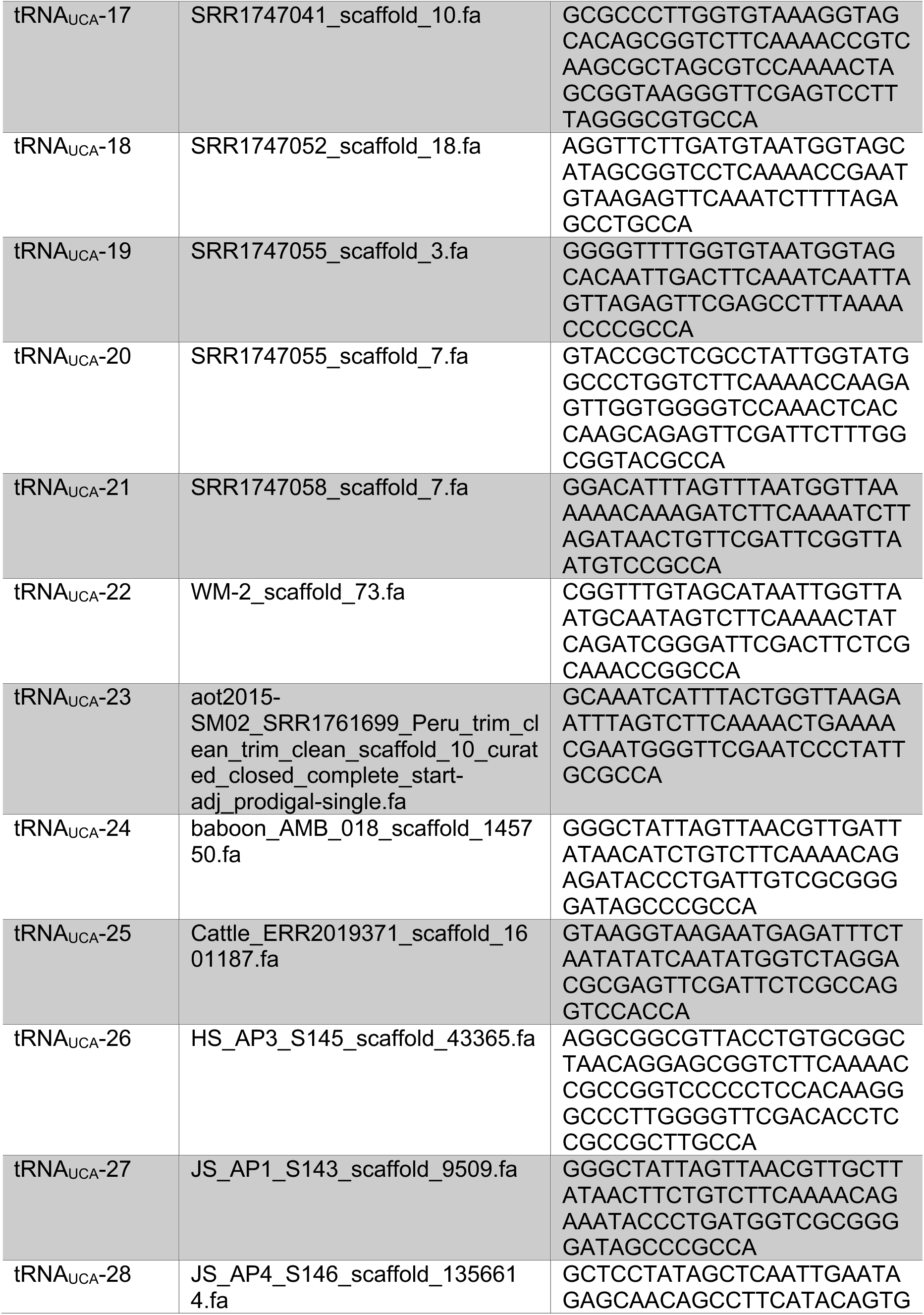
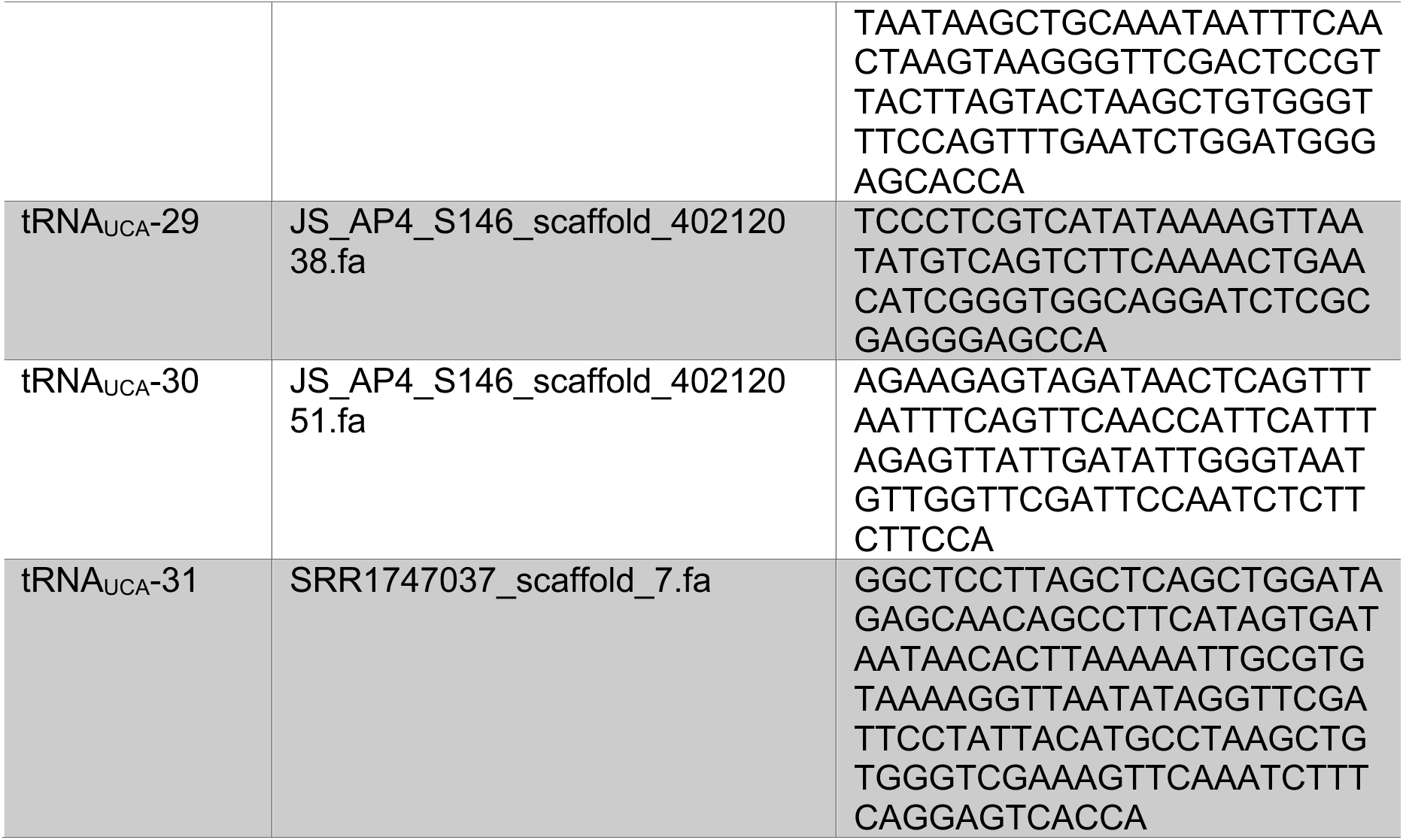
All bioinformatically identified suppressor tRNAs found in the genomes of bacteriophages.

**Table S2:**
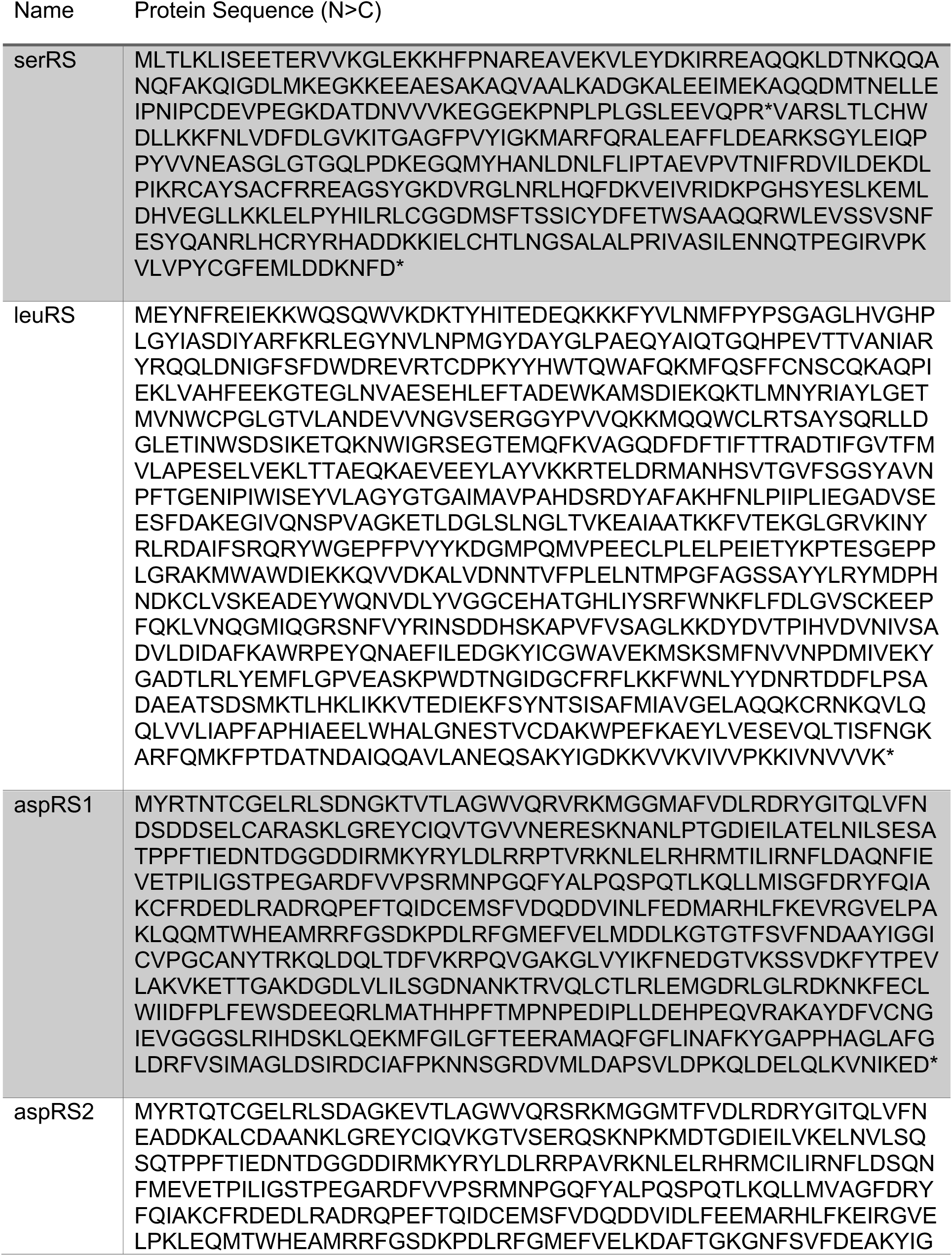
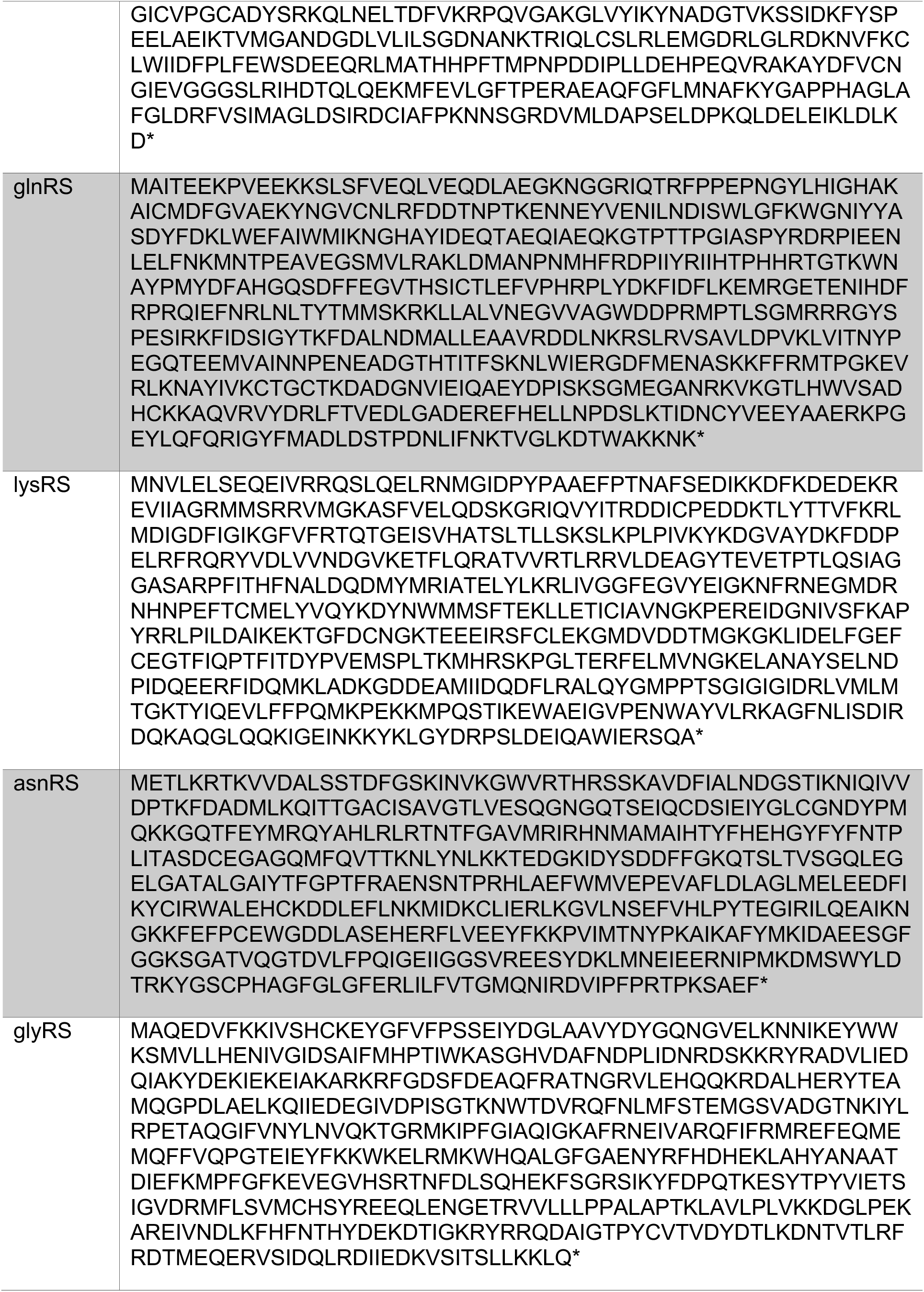
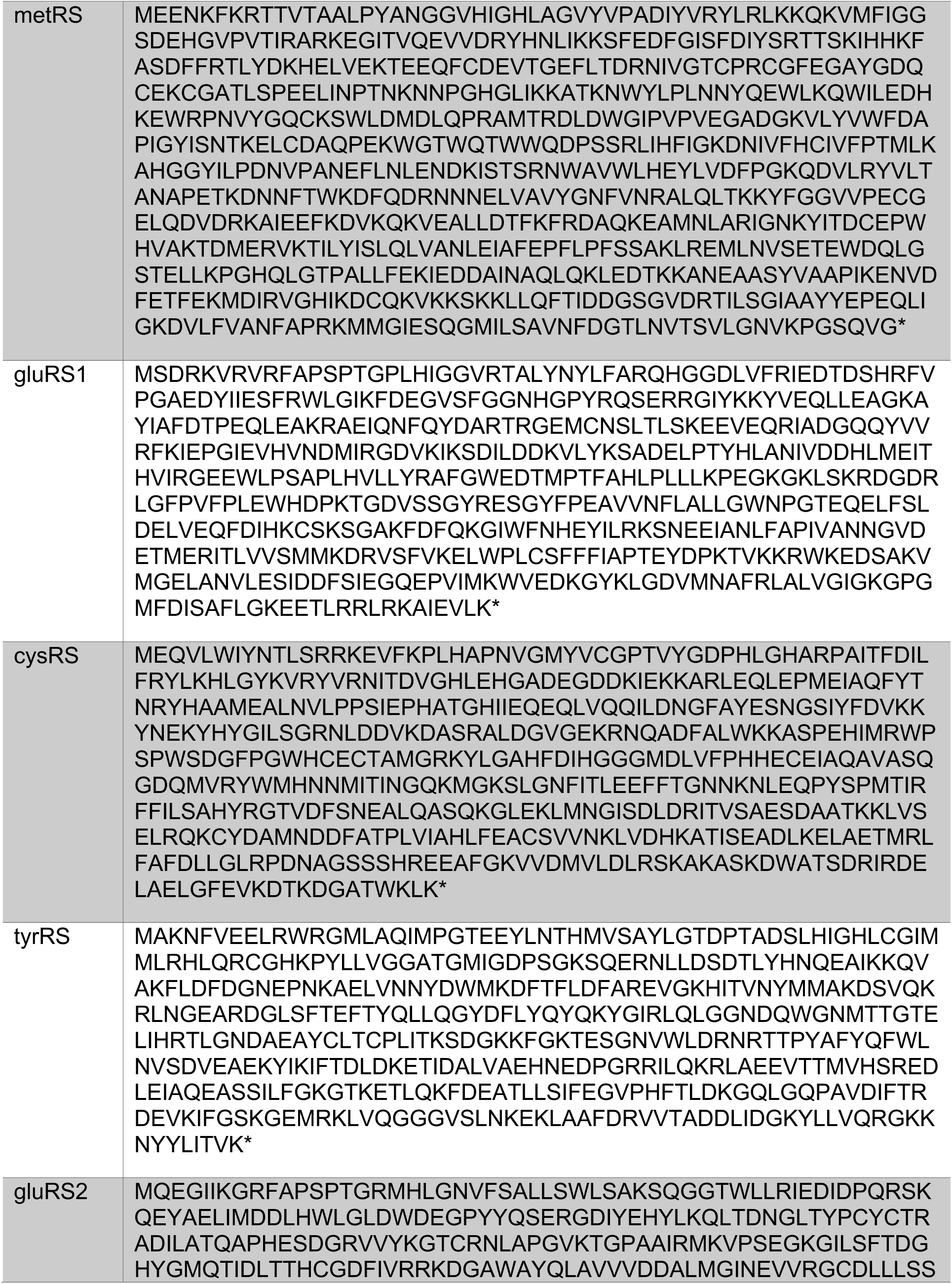
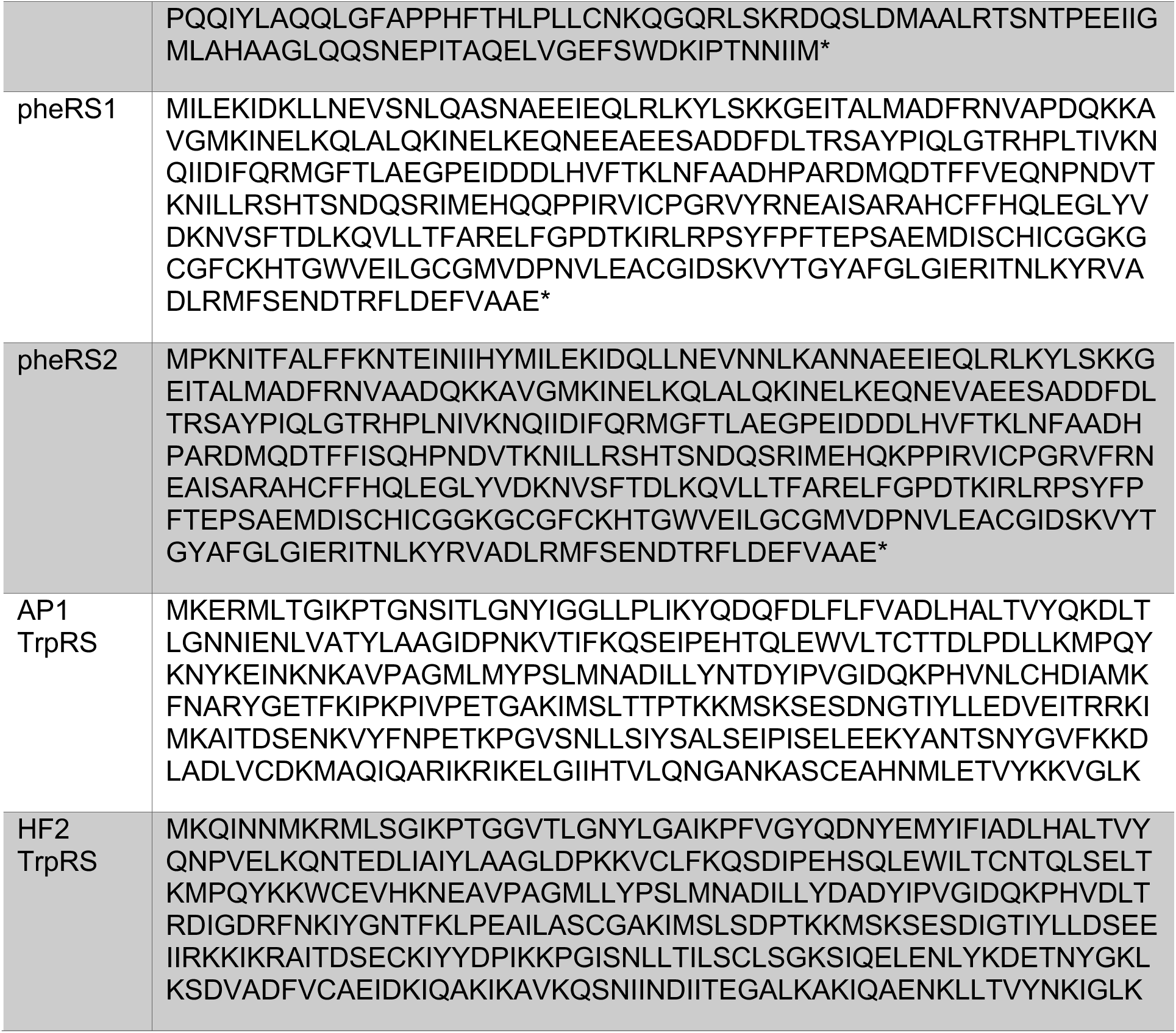
Protein sequences for bioinformatically identified aaRSs.

**Table S3:** Primers used for IVT template amplification.

| Identifier | Name | Sequence (5' - 3') |
| --- | --- | --- |
| tRNA<sub>CUA</sub>-1 | aot2015-HCO70_SRR1761673_Peru_scaffold_1.fa | TmGGCACCG<br>GCTCTCA |
| tRNA<sub>CUA</sub>-2 | aot2015-NO11_SRR1761685_USA_scaffold_53.fa | TmGGTAGGG<br>GTGGTAG |
| tRNA<sub>CUA</sub>-3 | aot2015-NO14_SRR1761688_USA_scaffold_60.fa | TmGGTAGCC<br>CCACTAG |
| tRNA<sub>CUA</sub>-4 | aot2015-NO17_SRR1761691_USA_scaffold_51.fa | TmGGTAGCC<br>CCACTAC |
| tRNA<sub>CUA</sub>-5 | aot2015-NO22_SRR1761696_USA_trim_clean_trim_clean_scaffo<br>ld_147.fa | TmGGTTGGG<br>GCAGTAG |
| tRNA<sub>CUA</sub>-6 | aot2015-NO23_SRR1761697_USA_trim_clean_trim_clean_scaffo<br>ld_74.fa | TmGGTCGCC<br>CTACCAC |
| tRNA<sub>CUA</sub>-7 | aot2015-SM02_SRR1761699_Peru_trim_clean_trim_clean_scaffo<br>ld_10_curated_closed_complete_start-adj_prodigal-single.fa | TmGGCACCG<br>GCTCTCG |
| tRNA<sub>CUA</sub>-8 | aot2015-SM23_SRR1761705_Peru_trim_clean_trim_clean_scaffo<br>ld_11.fa | TmGGGGAGG<br>GGGTGGG |
| tRNA<sub>CUA</sub>-9 | aot2015-SM39_SRR1761716_Peru_trim_clean_trim_clean_scaffo<br>ld_14_174.fa | TmGGGGAGG<br>GAGTGGG |
| tRNA<sub>CUA</sub>-10 | aot2015-SM40_SRR1761717_Peru_trim_clean_trim_clean_scaffo<br>ld_14_45.fa | TmGGCGGGG<br>GAAGTGG |
| tRNA<sub>CUA</sub>-11 | aot2015-SM40_SRR1761717_Peru_trim_clean_trim_clean_scaffo<br>ld_48.fa | TmGGTAGCC<br>CTAGTAG |
| tRNA<sub>CUA</sub>-12 | baboon_AMB_007_scaffold_42191.fa | TmGGCGGAG<br>GGGGTGG |
| tRNA<sub>CUA</sub>-13 | baboon_AMB_010_scaffold_110857.fa | TmGGTTCTCC<br>CACAAG |
| tRNA<sub>CUA</sub>-14 | baboon_AMB_018_scaffold_42888.fa | TmGGCGGTC<br>CCGCTGA |
| tRNA<sub>CUA</sub>-15 | baboon_AMB_024_scaffold_80789.fa | TmGGTTGGG<br>GCCAAGG |
| tRNA<sub>CUA</sub>-16 | C1--CH_A02_001D1_final_146.fa | TmGGTAGCC<br>CCACTCA |
| tRNA<sub>CUA</sub>-17 | C1--CH_A02_001D1_final_33.fa | TmGGTTGGG<br>GATCCGA |
| tRNA<sub>CUA</sub>-18 | Cattle_ERR2019359_scaffold_1067472.fa | TmGGTTGCTC<br>CGCTAG |
| tRNA<sub>CUA</sub>-19 | Cattle_ERR2019363_scaffold_1862390.fa | TmGGCGCGG<br>CATGAAG |
| tRNA<sub>CUA</sub>-20 | Cattle_ERR2019363_scaffold_240122.fa | TmGGTTGCCA<br>TGA CTG |
| tRNA<sub>CUA</sub>-21 | Cattle_ERR2019365_scaffold_547756.fa | TmGGTCACCC<br>GACTGG |
| tRNA<sub>CUA</sub>-22 | Cattle_ERR2019366_scaffold_1024626.fa | TmGGCAGCG<br>GTTCTCA |
| tRNA<sub>CUA</sub>-23 | Cattle_ERR2019366_scaffold_2103698.fa | TmGGCACCG<br>GCGTTGA |
| tRNA<sub>CUA</sub>-24 | Cattle_ERR2019371_scaffold_1572649.fa | TmGGAGACG<br>CGGACGG |
| tRNA<sub>CUA</sub>-25 | Cattle_ERR2019372_scaffold_259737_38.fa | TmGGTTTCCC<br>CACTAG |
| tRNA<sub>CUA</sub>-26 | Cattle_ERR2019372_scaffold_259737_83.fa | TmGGTTGCCC<br>AGCTCG |
| tRNA<sub>CUA</sub>-27 | Cattle_ERR2019372_scaffold_858714.fa | TmGGTTACTC<br>CGCTCG |
| tRNA<sub>CUA</sub>-28 | Cattle_ERR2019373_scaffold_1281151_48.fa | TmGGTTTCTC<br>CGCTTG |
| tRNA<sub>CUA</sub>-29 | Cattle_ERR2019373_scaffold_1281151_78.fa | TmGGTTGCCC<br>CGCAAG |
| tRNA<sub>CUA</sub>-30 | Cattle_ERR2019373_scaffold_1635329.fa | TmGGTTGTCC<br>GCCTAG |
| tRNA<sub>CUA</sub>-31 | Cattle_ERR2019374_scaffold_1442139.fa | TmGGCAGCC<br>CCTACTGG |
| tRNA<sub>CUA</sub>-32 | Cattle_ERR2019374_scaffold_1534423.fa | TmGGTCGTCC<br>CACCAT |
| tRNA<sub>CUA</sub>-33 | Cattle_ERR2019374_scaffold_1659420.fa | TmGGTCACCC<br>AGGCAG |
| tRNA<sub>CUA</sub>-34 | Cattle_ERR2019374_scaffold_1874516_109.fa | TmGGTTGCCC<br>CGCAAG |
| tRNA<sub>CUA</sub>-35 | Cattle_ERR2019374_scaffold_1874516_77.fa | TmGGTTTCTC<br>CGCTGC |
| tRNA<sub>CUA</sub>-36 | Cattle_ERR2019374_scaffold_52496.fa | TmGGTAGTCC<br>CACTAG |
| tRNA<sub>CUA</sub>-37 | Cattle_ERR2019374_scaffold_986541_43.fa | TmGGTTGCC<br>GCTCTGG |
| tRNA<sub>CUA</sub>-38 | Cattle_ERR2019374_scaffold_986541_72.fa | TmGGTTTCTC<br>CGCTTG |
| tRNA<sub>CUA</sub>-39 | Cattle_ERR2019375_scaffold_236249.fa | TmGGCAGAC<br>CCTGAAG |
| tRNA<sub>CUA</sub>-40 | Cattle_ERR2019376_scaffold_1215483.fa | TmGGCGCCC<br>CCGCTTC |
| tRNA<sub>CUA</sub>-41 | Cattle_ERR2019376_scaffold_18372.fa | TmGGCGGGG<br>GAGCTCG |
| tRNA<sub>CUA</sub>-42 | Cattle_ERR2019377_scaffold_1186659.fa | TmGGCCACC<br>CTTCTGG |
| tRNA<sub>CUA</sub>-43 | Cattle_ERR2019381_scaffold_37261.fa | TmGGTTTCTC<br>CGGAAG |
| tRNA<sub>CUA</sub>-44 | Cattle_ERR2019381_scaffold_760851_11.fa | TmGGCAGGA<br>GCTGCAG |
| tRNA<sub>CUA</sub>-45 | Cattle_ERR2019383_scaffold_1659107.fa | TmGGCACCC<br>CCAGTTC |
| tRNA<sub>CUA</sub>-46 | Cattle_ERR2019383_scaffold_1659116.fa | TmGGTCGCC<br>CCACTCA |
| tRNA<sub>CUA</sub>-47 | Cattle_ERR2019396_scaffold_971512.fa | TmGGTTGACC<br>CACTAG |
| tRNA<sub>CUA</sub>-48 | Cattle_ERR2019401_scaffold_184521.fa | TmGGTTTCTC<br>CGCTCG |
| tRNA<sub>CUA</sub>-49 | Cattle_ERR2019402_scaffold_680681.fa | TmGGTTGCCC<br>CACTAG |
| tRNA<sub>CUA</sub>-50 | Cattle_ERR2019412_scaffold_987789.fa | TmGGTCTTCC<br>CGCAAC |
| tRNA<sub>CUA</sub>-51 | Cattle_ERR2019415_scaffold_304737.fa | TmGGTTGCCC<br>CACTAG |
| tRNA<sub>CUA</sub>-52 | Cattle_ERR2019415_scaffold_941508.fa | TmGGTTGTCC<br>CACTAG |
| tRNA<sub>CUA</sub>-53 | FE-1_scaffold_25.fa | TmGGTACCCC<br>CACTAG |
| tRNA<sub>CUA</sub>-54 | FL-1_scaffold_38.fa | TmGGCGGGG<br>GCGGTGG |
| tRNA<sub>CUA</sub>-55 | FM_L-1_scaffold_13.fa | TmGGTAACCC<br>CACTAC |
| tRNA<sub>CUA</sub>-56 | GiantTortoise_AD_1_scaffold_57.fa | TmGGCGGGG<br>GGCCTGG |
| tRNA<sub>CUA</sub>-57 | H18_Tanzania_scaffold_0.fa | TmGGCAGAG<br>GAAGTGG |
| tRNA<sub>CUA</sub>-58 | H24_Tanzania_scaffold_6.fa | TmGGTTACCC<br>CACCAG |
| tRNA<sub>CUA</sub>-59 | H3_Tanzania_scaffold_15_1.fa | TmGGTCGCC<br>CCACTAG |
| tRNA<sub>CUA</sub>-60 | HS_AP3_S145_scaffold_427710.fa | TmGGTACCCC<br>GTGCAG |
| tRNA<sub>CUA</sub>-61 | HS_AP3_S145_scaffold_435061.fa | TmGGTTCTCT<br>CACAAG |
| tRNA<sub>CUA</sub>-62 | HS_AP3_S145_scaffold_527873.fa | TmGGCGCCC<br>GCTGTTT |
| tRNA<sub>CUA</sub>-63 | HS_AP3_S145_sc_10243_Lak_complete_trimmed_start<br>_adj_final.fa | TmGGTAGCC<br>CCACTCA |
| tRNA<sub>CUA</sub>-64 | IT3_Italy_scaffold_16.fa | TmGGTAGCC<br>CCACTAC |
| tRNA<sub>CUA</sub>-65 | IT5_Italy_scaffold_2 134.fa | TmGGTTCCT<br>ACTCTG |
| tRNA<sub>CUA</sub>-66 | IT5_Italy_scaffold_2 18.fa | TmGGTTGCCC<br>TACTAG |
| tRNA<sub>CUA</sub>-67 | js4906-20-5_S4_scaffold_13.fa | TmGGCCAGG<br>GAAACTG |
| tRNA<sub>CUA</sub>-68 | js4906-22-2_S9_scaffold_15.fa | TmGGCGGCG<br>GCGCTCA |
| tRNA<sub>CUA</sub>-69 | js4906-22-3_S10_scaffold_41.fa | TmGGTACCCC<br>CACTAG |
| tRNA<sub>CUA</sub>-70 | js4906-23-2_S13_scaffold_20.fa | TmGGTTACCC<br>CACTAG |
| tRNA<sub>CUA</sub>-71 | js4906-23-3_S14_scaffold_14.fa | TmGGTTCTCC<br>CACAAG |
| tRNA<sub>CUA</sub>-72 | js4906-23-3_S14_scaffold_31.fa | TmGGCACCG<br>GCTCTCG |
| tRNA<sub>CUA</sub>-73 | js4906-25-3_S22_scaffold_14.fa | TmGGCAGGC<br>CCGACTT |
| tRNA<sub>CUA</sub>-74 | js4906-26-4_S27_scaffold_91.fa | TmGGTACCCC<br>CGCTGG |
| tRNA<sub>CUA</sub>-75 | js4906-26-5_S28_scaffold_82.fa | TmGGTACCCC<br>CACTAG |
| tRNA<sub>CUA</sub>-76 | js4906-28-2_S33_scaffold_21.fa | TmGGTCCCC<br>CCACTAG |
| tRNA<sub>CUA</sub>-77 | JS_AP1_S143_scaffold_223286.fa | TmGGGGAGG<br>GAGTGGG |
| tRNA<sub>CUA</sub>-78 | JS_AP1_S143_scaffold_24016_curated_start_adj 66.fa | TmGGTTGGG<br>GATCCGA |
| tRNA<sub>CUA</sub>-79 | JS_AP4_S146_scaffold_135661.fa | TmGGTCGCC<br>CCACTCA |
| tRNA<sub>CUA</sub>-80 | JS_AP4_S146_scaffold_408865.fa | TmGGTTAGG<br>GCGGCAG |
| tRNA<sub>CUA</sub>-81 | JS_AP4_S146_scaffold_433782.fa | TmGGTCGCC<br>CCACCAT |
| tRNA<sub>CUA</sub>-82 | JS_AP5_S147_scaffold_73416.fa | TmGGTTACCC<br>CACTAG |
| tRNA<sub>CUA</sub>-83 | JS_HA1_S131_scaffold_901283.fa | TmGGTAGCC<br>CCGCTTG |
| tRNA<sub>CUA</sub>-84 | JS_HA3_S133_scaffold_534305.fa | TmGGCCCCCT<br>GAACGAA |
| tRNA<sub>CUA</sub>-85 | JS_HB1_S134_Phage_AC_29_17_curated 12.fa | TmGGTTGGG<br>GCGGTGG |
| tRNA<sub>CUA</sub>-86 | JS_HB1_S134_Phage_AC_29_17_curated 67.fa | TmGGTAGTCC<br>CATTGG |
| tRNA<sub>CUA</sub>-87 | JS_HB1_S134_scaffold_565796.fa | TmGGTGATCC<br>AGACAG |
| tRNA<sub>CUA</sub>-88 | JS_HE1_S137_scaffold_331486.fa | TmGGTTCCTC<br>GTATCG |
| tRNA<sub>CUA</sub>-89 | JS_HE2_S138_scaffold_134539.fa | TmGGCAGCC<br>CCACTAG |
| tRNA<sub>CUA</sub>-90 | JS_HE2_S138_scaffold_265266_1.fa | TmGGCGGGG<br>GAGCTCG |
| tRNA<sub>CUA</sub>-91 | JS_HE2_S138_scaffold_375748.fa | TmGGTTGTCC<br>TACTTG |
| tRNA<sub>CUA</sub>-92 | JS_HF1_S140_scaffold_125458.fa | TmGGCAGGG<br>GCGCTGA |
| tRNA<sub>CUA</sub>-93 | JS_HF2_S141_scaffold_334604.fa | TmGGCGGGG<br>GTGCAGG |
| tRNA<sub>CUA</sub>-94 | JS_HF3_S142_scaffold_71588.fa | TmGGCTGCCT<br>GTGTTC |
| tRNA<sub>CUA</sub>-95 | JS_HF3_S142_scaffold_7665.fa | TmGGTAGCCT<br>CACTGA |
| tRNA<sub>CUA</sub>-96 | L2_026_000M1_scaffold_35 129.fa | TmGGTTCCCC<br>ACTCTG |
| tRNA<sub>CUA</sub>-97 | L2_026_000M1_scaffold_35 26.fa | TmGGTTGCCC<br>TACTAG |
| tRNA<sub>CUA</sub>-98 | L2_038_000M1_scaffold_150.fa | TmGGTTCCCC<br>ATTCTG |
| tRNA<sub>CUA</sub>-99 | L2_057_000M1_scaffold_83.fa | TmGGTCGCC<br>CTACCAC |
| tRNA<sub>CUA</sub>-100 | L3_079_000M1_scaffold_477.fa | TmGGTAGCC<br>CCACTAG |
| tRNA<sub>CUA</sub>-101 | L3_105_000M1_scaffold_116.fa | TmGGCAGCC<br>CCGCTGG |
| tRNA<sub>CUA</sub>-102 | L3_122_000M1_scaffold_37.fa | TmGGTTCCCC<br>ACTCCG |
| tRNA<sub>CUA</sub>-103 | PHAGE-A2--js4906-20-<br>3_S2_Complete_Phage_26_29_curated.fa | TmGGTAGCC<br>CCACTCA |
| tRNA<sub>CUA</sub>-104 | PHAGE-A6--js4906-27-<br>3_S30_scaffold_5_curated_closed_complete_prodigal-single.fa | TmGGCTGCC<br>CTGCTTG |
| tRNA<sub>CUA</sub>-105 | pig_EL5596_F5_scaffold_126.fa | TmGGTTGGG<br>GCAGCCG |
| tRNA<sub>CUA</sub>-106 | pig_EL5596_F5_scaffold_275.fa | TmGGTAGCC<br>CCAGACT |
| tRNA<sub>CUA</sub>-107 | pig_EL5599_F8_scaffold_1425.fa | TmGGTCGCC<br>CCAGACT |
| tRNA<sub>CUA</sub>-108 | pig_ID_1851_F40_2_B1_scaffold_1589.fa | TmGGCTGCG<br>GCTGTCC |
| tRNA<sub>CUA</sub>-109 | pig_ID_1851_F40_2_scaffold_2 135.fa | TmGGCTGGG<br>GAGGTGG |
| tRNA<sub>CUA</sub>-110 | pig_ID_1851_F40_2_scaffold_2 47.fa | TmGGCGGGG<br>GTGGTCA |
| tRNA<sub>CUA</sub>-111 | pig_ID_2051_F75_scaffold_11.fa | TmGGTTGGG<br>GTCATGG |
| tRNA<sub>CUA</sub>-112 | pig_ID_2176_F64_scaffold_8.fa | TmGGTTACCC<br>CTCTAG |
| tRNA<sub>CUA</sub>-113 | pig_ID_2419_F67_scaffold_1281.fa | TmGGCTGCG<br>GCTGTCC |
| tRNA<sub>CUA</sub>-114 | pig_ID_3640_F65_scaffold_1252.fa | TmGGTCGCC<br>CCACTAG |
| tRNA<sub>CUA</sub>-115 | scaffold_104394.fa | TmGGTAGCC<br>CCGCCA |
| tRNA<sub>CUA</sub>-116 | SRR1747018_scaffold_13.fa | TmGGCAGAG<br>GAGACAG |
| tRNA<sub>CUA</sub>-117 | SRR1747020_scaffold_58.fa | TmGGCGAGG<br>GCGCTGG |
| tRNA<sub>CUA</sub>-118 | SRR1747021_scaffold_40.fa | TmGGCGGGG<br>GAGGCGG |
| tRNA<sub>CUA</sub>-119 | SRR1747021_scaffold_43.fa | TmGGCGGGG<br>GAAGTCG |
| tRNA<sub>CUA</sub>-120 | SRR1747023_scaffold_8.fa | TmGGTTGGG<br>GCGCTGA |
| tRNA<sub>CUA</sub>-121 | SRR1747024_scaffold_7.fa | TmGGTTGAAG<br>CGCCAG |
| tRNA<sub>CUA</sub>-122 | SRR1747025_scaffold_4.fa | TmGGCACCG<br>GCCCTCA |
| tRNA<sub>CUA</sub>-123 | SRR1747029_scaffold_6.fa | TmGGAAGCC<br>CTGGTAG |
| tRNA<sub>CUA</sub>-124 | SRR1747032_scaffold_7.fa | TmGGCGCCG<br>GCGCCCT |
| tRNA<sub>CUA</sub>-125 | SRR1747035_scaffold_3.fa | TmGGCAGAG<br>GAGACAG |
| tRNA<sub>CUA</sub>-126 | SRR1747039_scaffold_47.fa | TmGGTCGCC<br>CTAGTAG |
| tRNA<sub>CUA</sub>-127 | SRR1747044_scaffold_37.fa | TmGGTCGCC<br>CCACTCA |
| tRNA<sub>CUA</sub>-128 | SRR1747045_scaffold_26.fa | TmGGTAGCC<br>CATGTAG |
| tRNA<sub>CUA</sub>-129 | SRR1747046_scaffold_23.fa | TmGGCACCG<br>GCGCCTA |
| tRNA<sub>CUA</sub>-130 | SRR1747047_scaffold_5.fa | TmGGCTGAC<br>CTTCTCT |
| tRNA<sub>CUA</sub>-131 | SRR1747047_scaffold_8.fa | TmGGCACCG<br>GCTCTCG |
| tRNA<sub>CUA</sub>-132 | SRR1747052_scaffold_7_prodigal-single.fa | TmGGCTGTG<br>GAGTTGG |
| tRNA<sub>CUA</sub>-133 | SRR1747053_scaffold_18.fa | TmGGTCCCC<br>CCGGCCT |
| tRNA<sub>CUA</sub>-134 | SRR1747054_scaffold_15.fa | TmGGCACCG<br>GCTCTCG |
| tRNA<sub>CUA</sub>-135 | SRR1747060_scaffold_5.fa | TmGGTTGGG<br>GATGAGA |
| tRNA<sub>CUA</sub>-136 | SRR1747062_scaffold_47.fa | TmGGTCCCC<br>CCGGCCT |
| tRNA<sub>CUA</sub>-137 | SRR1747063_scaffold_2.fa | TmGGTTGAAG<br>CGCCAG |
| tRNA<sub>CUA</sub>-138 | SRR1747064_scaffold_16.fa | TmGGCGGAG<br>GAGGTGG |
| tRNA<sub>CUA</sub>-139 | SRR1747065_scaffold_11.fa | TmGGCAGCC<br>CTGCTGG |
| tRNA<sub>CUA</sub>-140 | SRR1747065_scaffold_18_106.fa | TmGGCTCCG<br>GGGTAGA |
| tRNA<sub>CUA</sub>-141 | SRR1747065_scaffold_33.fa | TmGGTTAGG<br>GTACTAG |
| tRNA<sub>CUA</sub>-142 | term6_stool_scaffold_62.fa | TmGGTTGGG<br>GCGGCAG |
| tRNA<sub>CUA</sub>-143 | WL-1_scaffold_34.fa | TmGGGCCCC<br>CGCCCGG |
| tRNA<sub>CUA</sub>-144 | WM-2_scaffold_7.fa | TmGGTCGCC<br>CCACTAG |
| tRNA<sub>CUA</sub>-145 | WM-2_scaffold_79.fa | TmGGGGCGG<br>GCCTGAG |
| tRNA<sub>CUA</sub>-146 | aot2015-HCO74_SRR1761675_Peru_scaffold_5.fa | TmGGTAGCC<br>CCACTCC |
| tRNA<sub>CUA</sub>-147 | Cattle_ERR2019372_scaffold_807595.fa | TmGGCAGCC<br>CGCCTGG |
| tRNA<sub>CUA</sub>-148 | Cattle_ERR2019383_scaffold_1237313.fa | TmGGCAGCC<br>CGCCTGG |
| tRNA<sub>CUA</sub>-149 | Cattle_ERR2019385_scaffold_165249.fa | TmGGTCCCG<br>GCGACCG |
| tRNA<sub>CUA</sub>-150 | Cattle_ERR2019408_scaffold_17433.fa | TmGGTCGGG<br>GTGCCCC |
| tRNA<sub>CUA</sub>-151 | HS_AP3_S145_scaffold_186886.fa | TmGGAGTGG<br>ACGGATC |
| tRNA<sub>CUA</sub>-152 | JS_AP1_S143_scaffold_24016_curated_start_adj.fa | TmGGTAGACC<br>CACTCA |
| tRNA<sub>CUA</sub>-153 | JS_HA1_S131_scaffold_412568.fa | TmGGCTGTCC<br>CGATCA |
| tRNA<sub>CUA</sub>-154 | JS_HE2_S138_scaffold_9600.fa | TmGGCTGTCC<br>CCGCCA |
| tRNA<sub>CUA</sub>-155 | L3_058_000M1_scaffold_129.fa | TmGGTCGGC<br>GCGGTAG |
| tRNA<sub>CUA</sub>-156 | PHAGE-A6--js4906-27-<br>3_S30_scaffold_5_curated_closed_complete_prodigal-single.fa | TmGGCTAGG<br>GATGTAG |
| tRNA<sub>CUA</sub>-157 | pig_F39_scaffold_16.fa | TmGGCGAGG<br>GCGGAGA |
| tRNA<sub>CUA</sub>-158 | pig_ID_3640_F65_scaffold_20.fa | TmGGTAGCC<br>CATGTAG |
| tRNA<sub>CUA</sub>-159 | SRR1747022_scaffold_143.fa | TmGGCAGGG<br>GTGCTGA |
| tRNA<sub>CUA</sub>-160 | SRR1747047_scaffold_438.fa | TmGGTCTCCC<br>CCGACA |
| tRNA<sub>CUA</sub>-161 | SRR1747065_scaffold_55.fa | TmGGCAGGG<br>GTGCTGA |
| tRNA<sub>CUA</sub>-162 | STL6-S73_scaffold_50.fa | TmGGTAGCC<br>CCACTCC |
| tRNA<sub>UUA</sub>-1 | aot2015-SM40_SRR1761717_Peru_trim_clean_trim_clean_scaffo<br>ld_14_145.fa | TmGGAAGCTT<br>AATGTG |
| tRNA<sub>UUA</sub>-2 | C1--CH_A02_001D1_final 93.fa | TmGGTGCTCT<br>ATACAG |
| tRNA<sub>UUA</sub>-3 | FL-1_scaffold_16.fa | TmGGTTGGC<br>CCACTAA |
| tRNA<sub>UUA</sub>-4 | JS_AP1_S143_scaffold_24016_curated_start_adj 97.fa | TmGGTGCTCT<br>ATACAG |
| tRNA<sub>UUA</sub>-5 | JS_HB1_S134_Phage_AC_29_17_curated 36.fa | TmGGCGGTC<br>TATGCAG |
| tRNA<sub>UUA</sub>-6 | LAK-phage_B10--M12_SRR1747042_26_21_curated.fa | TmGGTGCATG<br>ATACAG |
| tRNA<sub>UUA</sub>-7 | PHAGE-A1--js4906-22-<br>4_S11_scaffold_2_curated_complete.fa | TmGGTGCTCT<br>ATACAG |
| tRNA<sub>UUA</sub>-8 | SRR1747048_scaffold_38.fa | TmGGTACAGA<br>AAGTTG |
| tRNA<sub>UUA</sub>-9 | SRR1747065_scaffold_18 179.fa | TmGGGTCCG<br>AAGCGCG |
| tRNA<sub>UUA</sub>-10 | aot2015-SM39_SRR1761716_Peru_trim_clean_trim_clean_scaffo<br>ld_14.fa | TmGGCTCCAA<br>GGAAGG |
| tRNA<sub>UUA</sub>-11 | Cattle_ERR2019356_scaffold_1348925 42.fa | TmGGAGCCAT<br>AAGCGA |
| tRNA<sub>UUA</sub>-12 | Cattle_ERR2019356_scaffold_1348925 44.fa | TmGGTAGACT<br>GAGAAG |
| tRNA<sub>UUA</sub>-13 | Cattle_ERR2019375_scaffold_493821 32.fa | TmGGTAGACT<br>AAGAAG |
| tRNA<sub>UUA</sub>-14 | Cattle_ERR2019380_scaffold_1591051.fa | TmGGATGGTA<br>CGGGAT |
| tRNA<sub>UUA</sub>-15 | H18_Tanzania_scaffold_0.fa | TmGGAGCTTC<br>AACAAG |
| tRNA<sub>UUA</sub>-16 | JS_AP4_S146_scaffold_135661 9.fa | TmGGGAGAC<br>TATCTGG |
| tRNA<sub>UUA</sub>-17 | JS_HF3_S142_scaffold_205978 35.fa | TmGGTGGGC<br>ACGGTAG |
| tRNA<sub>UUA</sub>-18 | SRR1747021_scaffold_22.fa | TmGGTGCGA<br>ATGGAGG |
| tRNA<sub>UUA</sub>-19 | SRR1747023_scaffold_8.fa | TmGGAGGTCT<br>AAGAGA |
| tRNA<sub>UUA</sub>-20 | SRR1747056_scaffold_19.fa | TmGGCACCG<br>TCCCAGG |
| tRNA<sub>UCA</sub>-1 | baboon_AMB_014_scaffold_17184.fa | TmGGCACGT<br>CCTGAAG |
| tRNA<sub>UCA</sub>-2 | Cattle_ERR2019356_scaffold_1348925 168.fa | TmGGGGCGG<br>AGATGCG |
| tRNA<sub>UCA</sub>-3 | Cattle_ERR2019356_scaffold_1348925 8.fa | TmGGCAGGG<br>GCTGTAG |
| tRNA<sub>UCA</sub>-4 | Cattle_ERR2019366_scaffold_1317679.fa | TmGGCAAGG<br>GTGGTAG |
| tRNA<sub>UCA</sub>-5 | Cattle_ERR2019371_scaffold_1601187.fa | TmGGCACGC<br>CCCGTAG |
| tRNA<sub>UCA</sub>-6 | Cattle_ERR2019380_scaffold_916976.fa | TmGGCAGGG<br>GCTGTAG |
| tRNA<sub>UCA</sub>-7 | Cattle_ERR2019381_scaffold_760851 176.fa | TmGGTAATAT<br>CCAATC |
| tRNA<sub>UCA</sub>-8 | Cattle_ERR2019403_scaffold_427773.fa | TmGGCAGGA<br>GCTGTAG |
| tRNA<sub>UCA</sub>-9 | Cattle_ERR2019415_scaffold_672943.fa | TmGGCAGGG<br>CATGCAG |
| tRNA<sub>UCA</sub>-10 | GiantTortoise_AD_1_scaffold_344.fa | TmGGCAGGG<br>GCTGCAG |
| tRNA<sub>UCA</sub>-11 | JS_AP1_S143_scaffold_136784.fa | TmGGCCCTG<br>GAGGTAA |
| tRNA<sub>UCA</sub>-12 | JS_HE2_S138_scaffold_282274.fa | TmGGCATAG<br>GTTGAAG |
| tRNA<sub>UCA</sub>-13 | JS_HF2_S141_scaffold_159238.fa | TmGGTACCTC<br>CGCAAG |
| tRNA<sub>UCA</sub>-14 | SRR1747018_scaffold_51.fa | TmGGCACGC<br>CCTGTAG |
| tRNA<sub>UCA</sub>-15 | SRR1747025_scaffold_8.fa | TmGGCACCCCT<br>CGGCCC |
| tRNA<sub>UCA</sub>-16 | SRR1747029_scaffold_20.fa | TmGGCACGT<br>CCTGAAG |
| tRNA<sub>UCA</sub>-17 | SRR1747041_scaffold_10.fa | TmGGCACGC<br>CCTAAAG |
| tRNA<sub>UCA</sub>-18 | SRR1747052_scaffold_18.fa | TmGGCAGGC<br>TCTAAAA |
| tRNA<sub>UCA</sub>-19 | SRR1747055_scaffold_3.fa | TmGGCGGGG<br>TTTTAAA |
| tRNA<sub>UCA</sub>-20 | SRR1747055_scaffold_7.fa | TmGGCGTAC<br>CGCCAAA |
| tRNA<sub>UCA</sub>-21 | SRR1747058_scaffold_7.fa | TmGGCGGAC<br>ATTAACC |
| tRNA<sub>UCA</sub>-22 | WM-2_scaffold_73.fa | TmGGCCGGT<br>TTGCGAG |
| tRNA<sub>UCA</sub>-23 | aot2015-SM02_SRR1761699_Peru_trim_clean_trim_clean_scaffo<br>ld_10_curated_closed_complete_start-adj_prodigal-single.fa | TmGGCGCAAT<br>AGGGAT |
| tRNA<sub>UCA</sub>-24 | baboon_AMB_018_scaffold_145750.fa | TmGGCGGGC<br>TATCCCC |
| tRNA<sub>UCA</sub>-25 | Cattle_ERR2019371_scaffold_1601187.fa | TmGGTGGAC<br>CTGGCGA |
| tRNA<sub>UCA</sub>-26 | HS_AP3_S145_scaffold_43365.fa | TmGGCAAGC<br>GGCGGAG |
| tRNA<sub>UCA</sub>-27 | JS_AP1_S143_scaffold_9509.fa | TmGGCGGGC<br>TATCCCC |
| tRNA<sub>UCA</sub>-28 | JS_AP4_S146_scaffold_135661 4.fa | TmGGTGCTCC<br>CATCCA |
| tRNA<sub>UCA</sub>-29 | JS_AP4_S146_scaffold_402120 38.fa | TmGGCTCCCT<br>CGCGAG |
| tRNA<sub>UCA</sub>-30 | JS_AP4_S146_scaffold_402120 51.fa | TmGGAAGAA<br>GAGATTG |
| tRNA<sub>UCA</sub>-31 | SRR1747037_scaffold_7.fa | TmGGTGACTC<br>CTGAAA |

